# Insights into mechanisms of ATM activation via constitutively active mutants

**DOI:** 10.64898/2026.07.15.738190

**Authors:** Md Saiful Islam, Verena G. Gautsch, Rimma Belotserkovskaya, Almudena Serrano-Benitez, Dénes Buzafalvi, Olga Perisic, Stephen P. Jackson, Roger L. Williams

**Affiliations:** MRC Laboratory of Molecular Biology, Francis Crick Avenue, Cambridge, CB2 0QH, UK; Cancer research UK (CRUK) Cambridge Institute, University of Cambridge, Cambridge, CB2 0RE, UK

## Abstract

The Ser/Thr kinase ATM orchestrates cellular responses to DNA double-strand breaks (DSBs) and promotes DSB repair by homologous recombination. In this process, ATM is activated by DNA and the MRN (MRE11, RAD50, and NBS1) complex. Here we show that mutations of the conserved PIKK regulatory domain (PRD) within ATM’s kinase domain can confer a maximally active state that no longer requires MRN/DNA. In ATM-knockout human cells, the PRD mutants display substantially higher phosphorylation of histone H2AX, KAP1, and CHK2 than wild-type ATM, with or without IR-induced DNA damage. Cryo-EM structures of two PRD mutants each revealed basal or activated conformations depending on bound ligands, suggesting that disrupting the ordered portion of the PRD results in an enzyme poised to transition to the active conformation. However, the identity of the active-site nucleotide is a key driver of the conformational switching. We speculate that this plasticity might be exploited to develop small-molecule ATM modulators for therapeutic applications.

## Introduction

DNA double-strand breaks (DSBs) are the most lethal form of DNA damage and in eukaryotes are mainly repaired by non-homologous end joining (NHEJ) or homologous recombination (HR) associated with activation of the DNA-damage signaling kinases ATM (ataxia-telangiectasia mutated), ATR (ATM- and Rad3-related protein), and DNA-PK (DNA-dependent protein kinase) (*1, 2*). During DSB repair, ATM signals the presence of DSBs by catalyzing phosphorylation of proteins involved in checkpoint activation, DNA repair, apoptosis, and senescence (*3, 4*). A critical step is the activation of ATM, which is mediated by the heterotrimeric MRN complex consisting of the nuclease MRE11, the ATPase RAD50, and the adaptor NBS1 (Nibrin) (*5–7*) (fig. S1A). Additionally, ATM is activated by reactive oxygen species (ROS) and has roles in intracellular redox management (*4, 8–10*) (fig. S1A). This ATM function also has clinical implications in the rare autosomal recessive disorder, ataxia-telangiectasia (A-T). A-T patients have mutations in *ATM*, leading to non-functional or hypofunctional ATM protein, which causes radiosensitivity, premature ageing, cerebellar dysfunction, and cancer predisposition (*11*).

ATM is a Ser/Thr protein kinase that belongs to the phosphoinositide-3-kinase (PI3K) related protein kinase (PIKK) family, which also includes ATR, DNA-PK, mTOR, TRRAP, and SMG-1 (*1, 12, 13*) (fig. S1B). PIKKs share a homologous C-terminal region known as the FATKIN, consisting of a FAT domain (∼ 700 residues) followed by a kinase domain (∼400 residues) (Fig. 1A, fig. S1B). The kinase domain has a typical bilobal structure, with a catalytic cleft formed between the N-lobe and the C-lobe, and a PIKK regulatory domain (PRD) of a variable length within the C-lobe (*1, 3, 7, 10*). Unlike their C-termini, the N-terminal regions differ greatly among the PIKKs and consist of α-solenoid structures built of mainly HEAT repeats, which in ATM are primarily involved in interactions with DNA and protein partners (*3, 7, 10, 14*). All PIKKs undergo a large change in catalytic activity from basal to activated state, with basal activities so low that it is difficult to measure, while the activated states have *k*_cat_ values similar to an average enzyme (*15*). Among the six PIKKs, both basal/inactive and activated structures have been reported for ATM (*7, 10, 14, 16*), mTORC1 (*17, 18*), DNA-PK (*19–21*) and ATR (*22, 23*). Upon activation, all structures show large conformational changes occurring outside the kinase domain. In addition, there are smaller, but critically important, changes within the kinase domain, shifting the kinase N-lobe relative to the C-lobe, bringing the N-lobe and bound ATP into better register with the catalytic residues in the C-lobe, thereby increasing catalytic rate. The shifts of the N-lobe relative to the C-lobe seem to be a universal feature of the active state of PIKKs, including ATM.

**Fig. 1.**
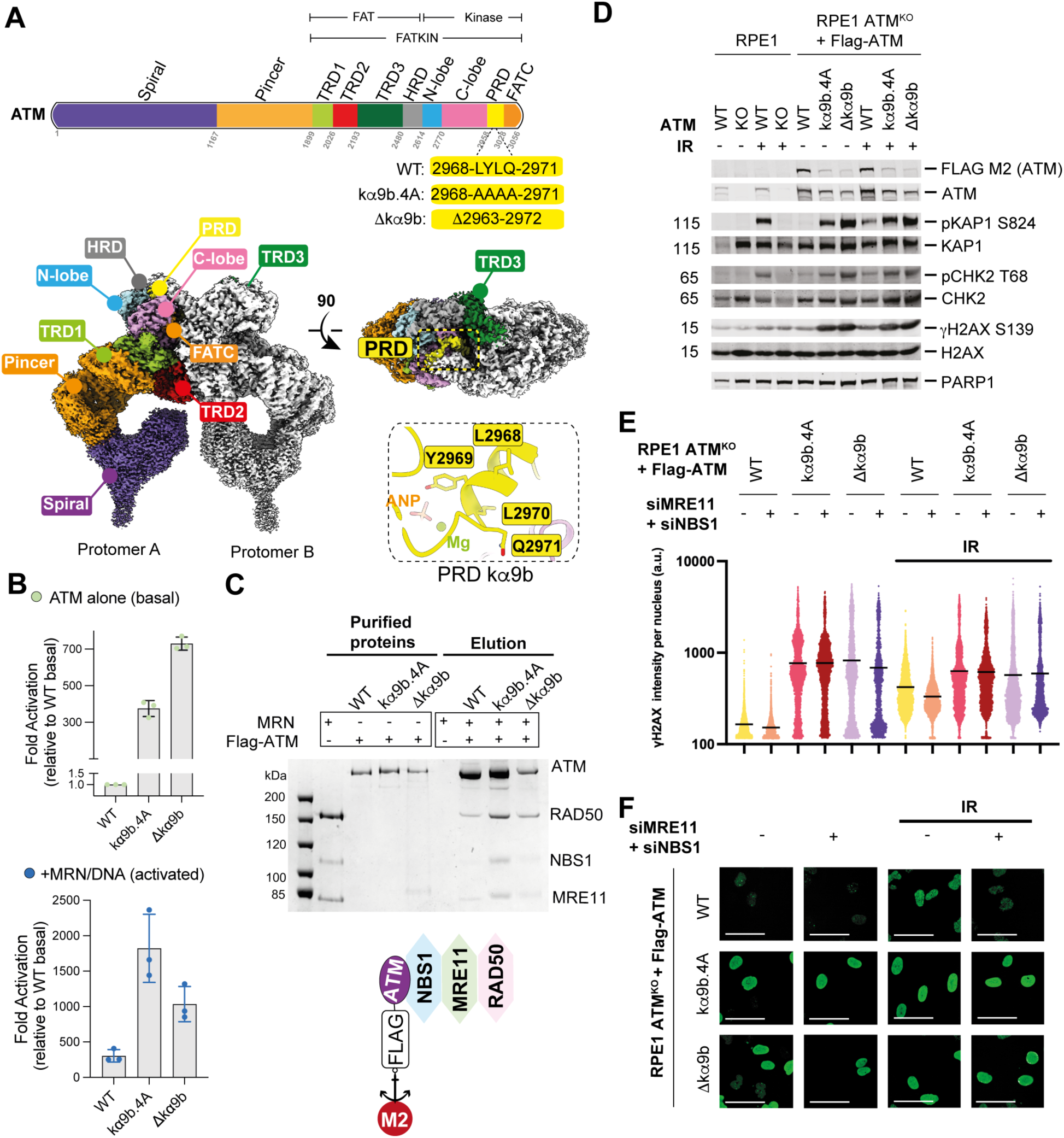
PRD mutations constitutively activate ATM. (A) A schematic diagram of ATM domain organization (top), highlighting the PRD mutations described in this study. Shown below the diagram are two orthogonal views of the cryo-EM density map of a wild-type ATM dimer (WT ATM, 7SIC), with one protomer colored as in the bar diagram. Also shown is the expanded view of PRD helix kα9b. (B) Comparison of the catalytic activity (*k*_cat_) values (n = 3) of ATM WT and PRD mutants in the presence or absence of MRN/DNA relative to the basal activity of WT ATM, based on data shown in fig. S2B. (C) A protein gel showing MRN complex pulled down on ATM. Flag-ATM variants were bound to anti-Flag M2 affinity beads (see fig. S2, C and D) and following incubation with MRN/DNA, bound components were eluted with a Flag peptide. Purified proteins and eluted samples were analyzed by SDS-PAGE gel. (D) Western blot analysis of phosphorylation of ATM targets, Ser^824^ of KAP1, Thr^68^ of CHK2, and Ser^139^ of H2AX, from wild-type and ATM KO RPE1 cells (lanes 1-5) and ATM KO RPE1 cells expressing inducible Flag-tagged ATM WT or ATM PRD mutants (lanes 6-10). We observed higher levels of target phosphorylation for the PRD mutants than the WT, and unlike WT, PRD mutants were active even without IR exposure (see fig. S3A). Cells were transduced with a lentiviral vector expressing doxycycline-inducible Flag-ATM constructs. Where indicated, cells were exposed to 2 Gy ionizing radiation (IR) and harvested after 1h recovery. IR = Ionizing radiation of 2 Gy, 1h recovery. Cells were treated with doxycycline prior to the experiment (0.1 µg/mL, 24h). (E) γH2AX mean intensity (n = 3) measured in *ATM* and *TP53* double knockout (KO) RPE1 cells with and without the depletion of MRE11/NBS1 transduced with Flag-tagged ATM variants. ATM PRD mutants show a higher basal activity compared to the WT, and γH2AX signal for the PRD mutants could not be decreased to baseline by MRE11/NBS1 depletion. Horizontal line indicates mean. Depletion efficiency is shown in fig. S3C. (F) Representative confocal images of the *ATM* and *TP53* double knockout (KO) RPE1 cells used to measure the γH2AX signal for the construction of the plots (Fig. 1E). Scale bar, 50 µm.

Human and yeast ATM orthologues form a butterfly-shaped homodimer where PRDs from the two protomers adopt an “autoinhibitory” conformation by occupying the substrate binding pockets (Fig. 1A, *movie clip 1*). Recently, Howes *et al.* reported an active-state structure of ATM elicited by oxidation (*10*). Here, we characterized structures and function of two constitutively active PRD-mutated ATMs. In both a cell free reconstituted system and in cellular assays with *ATM* knockout (KO) human cells, we show that ATM variants harbouring PRD kα9b mutations are catalytically more active than the wild-type enzyme and that these hyperactive ATM mutants efficiently phosphorylate their substrates without requiring MRN complex. The cryo-EM structures of two PRD variants show that the autoinhibitory PRD helix kα9b is displaced from the substrate binding pocket, rendering the enzyme poised for a basal-to-active-state transition. The PRD displacement elicits global conformational changes that facilitate efficient substrate-binding and catalysis, while still maintaining the ATM dimeric state. Furthermore, our cryo-EM structures of the most active of the mutants demonstrates that in the presence of a model substrate, ATM can form a stable active-state dimer when the nucleotide pocket is occupied by a transition state mimic (ADP/MgF3^-^), substrate (ATP) or product (ADP).

## Results

### ATM PRD mutants are hyperactive and function in an MRN-independent manner in vitro

We purified ATM variants from transiently transfected human Expi293F cells. Kinase activity was monitored using an *in vitro* reconstituted system consisting of purified ATM, MRN, linear double-stranded DNA (dsDNA), and p53^102^ as a substrate (MBP-p53, residues 1-102, p53^102^) (*10*). The basal activity of the wild-type ATM (WT) was very low, but it could be potently stimulated (∼270-fold) by MRN/dsDNA (basal *k*_cat_, 0.003 s^-1^; MRN/DNA activated *k*_cat_, 0.8 s^-1^) (Fig. 1B and fig. S2B). However, a variant with a mutation of PRD kα9b residues ^2968^LYLQ^2971^ to ^2968^AAAA^2971^ (kα9b.4A) (Fig. 1A) had ∼330-fold higher basal activity than the WT enzyme, and its *k*_cat_ could be activated only 4.8-fold by MRN/dsDNA (basal *k*_cat_, 1.0 s^-1^; MRN/DNA activated *k*_cat_, 4.8 s^-1^) (Fig. 1B and fig. S2, A and B). When we tested the ability of kα9b.4A to bind MRN in the presence of dsDNA, we found that it could bind MRN more efficiently than WT ATM (Fig. 1C and fig. S2, C and D), demonstrating that the mutation did not disrupt ATM/MRN interactions. Further, a PRD variant with deletion of PRD residues 2963-2972, including helix kα9b (Δkα9b), (Fig. 1A) had ∼630 fold higher basal activity relative to WT and could be only marginally activated by MRN/dsDNA (basal *k*_cat_, 1.9 s^-1^; MRN/DNA activated *k*_cat_, 2.7 s^-1^) (Fig. 1B and fig. S2A, B), corroborating what was reported previously by Howes *et al.* (*10*). Like kα9b.4A, Δkα9b also displayed enhanced binding of MRN relative to the WT enzyme (Fig. 1C and fig. S2, C and D), demonstrating again that the PRD kα9b mutation did not prevent ATM/MRN interactions.

### ATM PRD mutants are hyperactive and function in an MRN-independent manner in cells

To ascertain whether ATM PRD mutants are hyperactive in a cellular environment, we complemented *ATM TP53* double knockout (KO) non-transformed human RPE1 cells (*24*) with doxycycline-inducible Flag-tagged ATM variants and examined their kinase activity under basal conditions and after ionizing radiation (IR)-induced DNA damage by monitoring the phosphorylation of the ATM targets CHK2, histone H2AX, and KAP1 (Fig. 1D and fig. S3A). Notably, WT ATM promoted phosphorylation of KAP1 on Ser^824^, H2AX on Ser^139^ (γH2AX) and CHK2 on Thr^68^ only after IR-induced DNA damage. In contrast, cells expressing PRD mutants kα9b.4A and Δkα9b exhibited substantially higher phosphorylation levels even in the absence of DNA damage, and these levels remained similar following irradiation (Fig. 1D and fig. S3A). Because the RPE1 cell line used for ATM complementation experiments was *ATM⁻/⁻ TP53⁻/⁻*, a different cell line was used to assay ATM-dependent phosphorylation of p53 on Ser^15^. For this, we transiently transfected Flag-tagged ATM constructs into HEK293 cells and detected p53 Ser^15^ phosphorylation by immunoblot (fig. S3B). Compared to WT ATM, both PRD mutants were hyperactive towards p53 Ser^15^. Collectively, these data showed that the mutants were constitutively active in cells and, in contrast to the WT ATM, do not require activation by IR-induced DNA damage to phosphorylate cellular substrates. To test whether kα9b.4A and Δkα9b function in cells in an MRN-independent manner, we depleted MRE11 and NBS1 using short-interfering RNAs (siRNAs) and measured the mean nuclear intensity of γH2AX before and after IR induction (Fig. 1, E and F and figs. S3C, S4, and S5). Our immunofluorescence analyses of WT ATM revealed that γH2AX levels decreased upon depletion of MRE1 and NBS1. In contrast, for the kα9b.4A and Δkα9b mutants, there was a comparable γH2AX signal intensity in the presence or absence of MRE11 and NBS1 (Fig. 1, E and F and figs. S3C, S4, S5). IR exposure increased γH2AX for WT but had almost no influence for the mutants.

### The PRD is a critical block to ATM activation and binding of p53

To understand how PRD mutants overcome normal regulation and achieve constitutive activation, we determined their cryo-EM structures. Because we believed that the hyperactive PRD mutants might provide a good starting point for understanding ATM’s phosphoryl transfer, we determined cryo-EM structures of the kα9b.4A and Δkα9b in the presence of a p53 substrate peptide (p53^11-22^: EPPLSQETFSDL, p53^12^) and a transition-state mimic (TS, ADP/MgF_3_^-^) (*25–27*). For comparison, we also determined a structure of the WT ATM treated in the same fashion (with ADP/MgF_3_^-^/p53^12^).

Cryo-EM maps enabled us to resolve ATM homodimer structures in the presence of TS mimic, ADP/MgF_3_^-^, and p53^12^ for WT, kα9b.4A, and Δkα9b at 3.1 Å, 3.4 Å, and 3.3 Å resolutions, respectively (Fig. 2 and figs. S6-S12, table S1). For all three constructs, the ATM dimeric interfaces are formed by contacts between the symmetry-related FATKINs (Fig. 2A). These dimeric contacts involve four sets of interactions: TRD2/TRD2’ (I), TRD2/TRD3’ (II) and kinase domain interactions with TRD3’ (III & IV) (Fig. 2). The first two sets of interactions involving TRD2 with TRD2’ (I) and TRD2 with TRD3’ (II) are very similar among the WT and the mutants (Fig. 2A). In contrast, the other two sets of dimeric interactions (III and IV) formed between the kinase domain of one protomer and TRD3’ (helices fα19’ to fα22’, residues 2327-2476) differ markedly between the WT and the mutants (Fig. 2A). Because of the PRD kα9b mutations (kα9b.4A or Δkα9b), the PRD helix absent from the substrate binding pocket (PRD Out) and the contacts between the TRD3’ long coiled coil helices fα21’/fα22’ (residues 2377-2476) and the PRD helix are lost in the mutant structures (Lid off) (Fig. 2B and fig. S13, B and C). Conversely, in WT ATM, treated with the TS mimic and p53^12^ in the same way as the PRD mutants, the PRD helix kα9b occupies the substrate-binding pocket (PRD In), with the fα21’/fα22’ of the paired protomer leaning against the kα9b (Lid on) and stabilizing the “PRD In” conformation (Fig. 2B and fig. S13A). The presence of the PRD helix kα9b in the substrate-binding pocket (Fig. 2 and fig. S13) prevents the p53^12^ peptide binding in the active site, as suggested previously (*7, 10, 14, 16, 28*). This explains why the WT ATM is less active in the basal state. The opening of the substrate-binding sites for kα9b.4A and Δkα9b variants, in the presence of a TS mimic enables the p53^12^ peptide to bind to the active-site (Fig. 2, B and C and fig. S13).

**Fig. 2.**
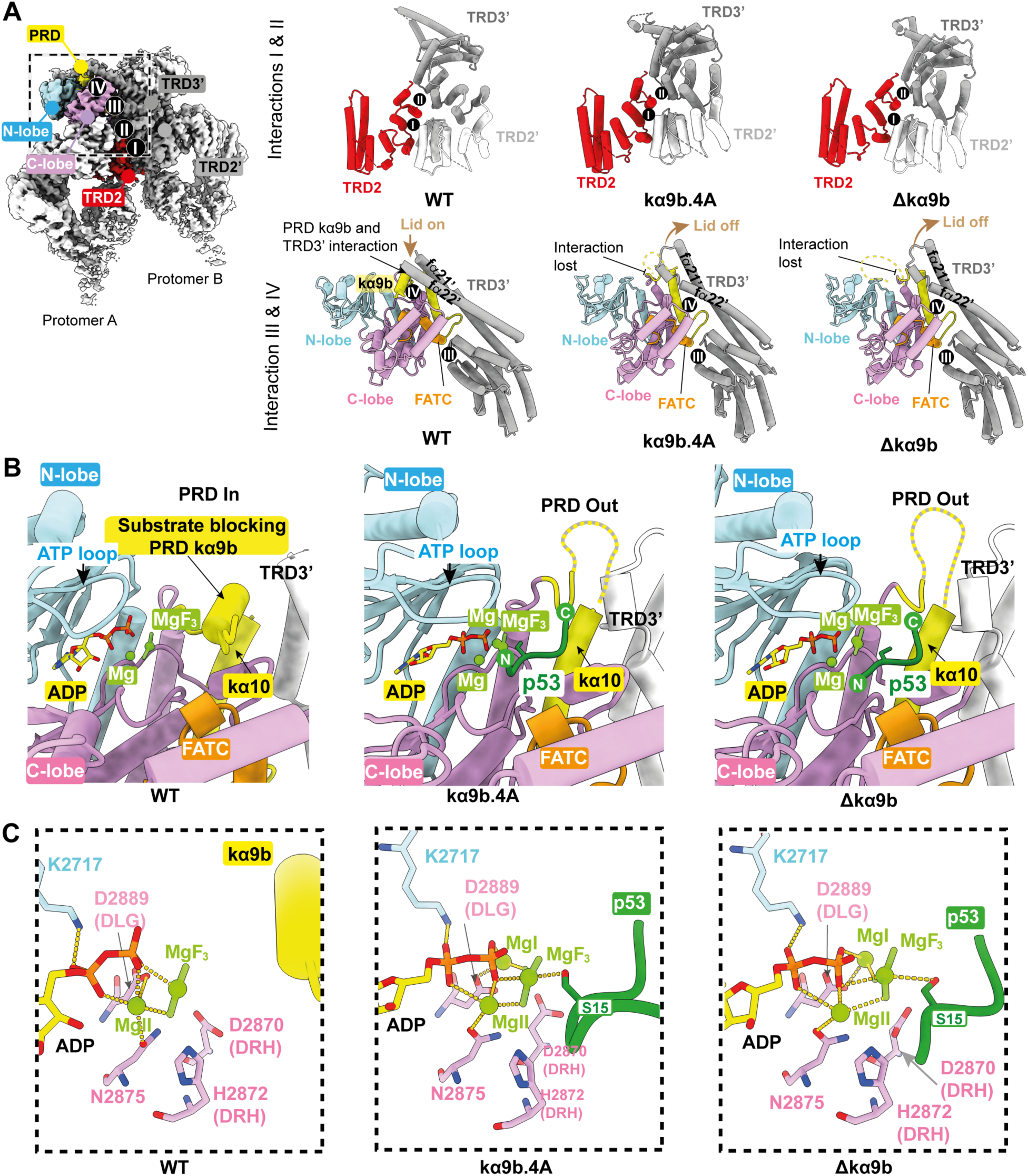
TS structures and interactions of ATM WT and PRD mutants. (A) Cryo-EM density map of a WT ATM dimer (left panel), with the interfaces between the two protomers for WT, kα9b.4A and Δkα9b shown on the right panels. The upper three panels show that interactions of the TRD2 domain of one protomer (red) with TRD2’ (white) (interaction I) and TRD3’ (gray) (interaction II) are very similar among the wild-type and the mutants. The three lower panels show that the interactions of the kinase domain of one protomer (colored as in a bar diagram of Fig. 1A) with TRD3’ (gray) (helices fα19’ to fα22’, residues 2327-2476) of the symmetry-related molecule (interactions III and IV) differ greatly between wild-type and the mutants. (B) Cryo-EM models of the kinase domain of the ATM variants, colored as in Fig. 1A. Samples were prepared in the presence of p53^12^ peptide and TS mimic ADP/MgF_3_^-^, showing that p53^12^ (dark green) binds to ATM when the PRD kα9b does not occupy the substrate binding pocket (“PRD Out”) in kα9b.4A and Δkα9b structures, whereas PRD kα9b still occupies the substrate pocket (“PRD In”) in WT ATM, preventing binding of the p53^12^ peptide. (C) Zoom into the active site of WT, kα9b.4A, and Δkα9b treated with p53^12^ peptide and TS mimic ADP/MgF_3_^-^. While WT ATM shows only ordered ADP/MgF_3_^-^ and one Mg^2+^ in the active site, kα9b.4A and Δkα9b have ADP/MgF_3_^-^, p53^12^ peptide and two Mg^2+^ ions bound in the active site. ADP is colored yellow, MgF_3_^-^ and Mg^2+^ ions light green, and p53^12^ peptide dark green.

### Ligand binding to the active site of PRD mutants

The architectures of the active-site of kα9b.4A and Δkα9b with bound ADP, MgF_3_^-^, p53 peptide, and Mg^2+^ ions are shown in Figure 2C. The trigonal-planar MgF_3_^-^ is positioned between the β-phosphate of ADP and the phospho-acceptor Ser^15^ of the p53 substrate. In the case of Δkα9b, we could model four (^14^LSQE^17^) out of twelve residues of the p53 peptide, whereas for kα9b.4A seven residues (^11^EPPLSQE^17^) were modelled. Compared with the WT TS structure and previous structures of ATM and the yeast homologue Tel1^ATM^ bound to an ATP mimic, which all have only a single Mg^2+^ bound (MgII), the TS structures of kα9b.4A and Δkα9b have two Mg^2+^ ions (MgI and MgII) in their active sites (Fig. 2C). The second Mg^2+^ (MgI) is coordinated by two oxygens of D2889 (from the DLG motif analogous to the DFG motif of canonical protein kinases), the β-phosphate of ADP, and the γ-phosphate mimic MgF_3_^-^. The binding of a second Mg^2+^ ion (as in kα9b.4A and Δkα9b) is often a rate limiting step in kinase catalysis (*29*), and it is likely that its capture in TS structures was facilitated both by the presence of MgF_3_^-^, which mimics the PO_3_^-^ intermediate, and by the p53 substrate.

In both Δkα9b and kα9b.4A structures, we modelled the phosphoacceptor Ser^15^ of the p53 peptide (PhosSer^15^) next to the ADP/MgF_3_^-^, with the N-terminus of the peptide (^11^EPPL^14^) towards the kinase C-lobe and the C-terminus (^16^QET^18^) towards the TRD3’ (Fig. 2B). This orientation of the p53 peptide, which has higher correlation coefficient and Q-score values than the reverse orientation (fig. S14), agrees with the previously reported p53-bound ATM and Upf1-bound SMG-1 structures (*10, 30*).

### PRD mutants with a bound TS mimic have a global active-state conformation

In addition to remodeling the active site to allow substrate peptide to bind, the kα9b.4A and Δkα9b mutants that remove the PRD elicit global conformational changes relative to the WT in the presence of a TS mimic (Fig. 3). The first global change is that TRD2, TRD3 and HRD regions of the FAT domain (FAT-clamp) slide as a unit (Fig. 3A and *movie clip 2*). The space created by this movement allows the kinase N-lobe to shift into its optimal position relative to the C-lobe (Fig. 3B and C). This N-lobe realignment results in a substantial movement of the ATP loop (residues 2693-2699) and brings the ATP γ-phosphate into better register with the catalytic residues in the C-lobe and with the acceptor hydroxyl group of the peptide substrate (Fig. 3B and C and *movie clip 3*). Conversely, no such N-lobe to C-lobe realignment was observed for the WT ATM, although the cryo-EM sample was treated with ADP/MgF_3_^-^/p53 in the same manner as the kα9b mutants. Coincident with the FAT clamp sliding, helices fα21’/fα22’ from TRD3’ retract toward the two-fold axis of the dimer (Fig. 3D and *movie clip 4*). This has important consequences for the substrate binding site of the symmetry-related protomer and for the dimeric interface.

**Fig. 3.**
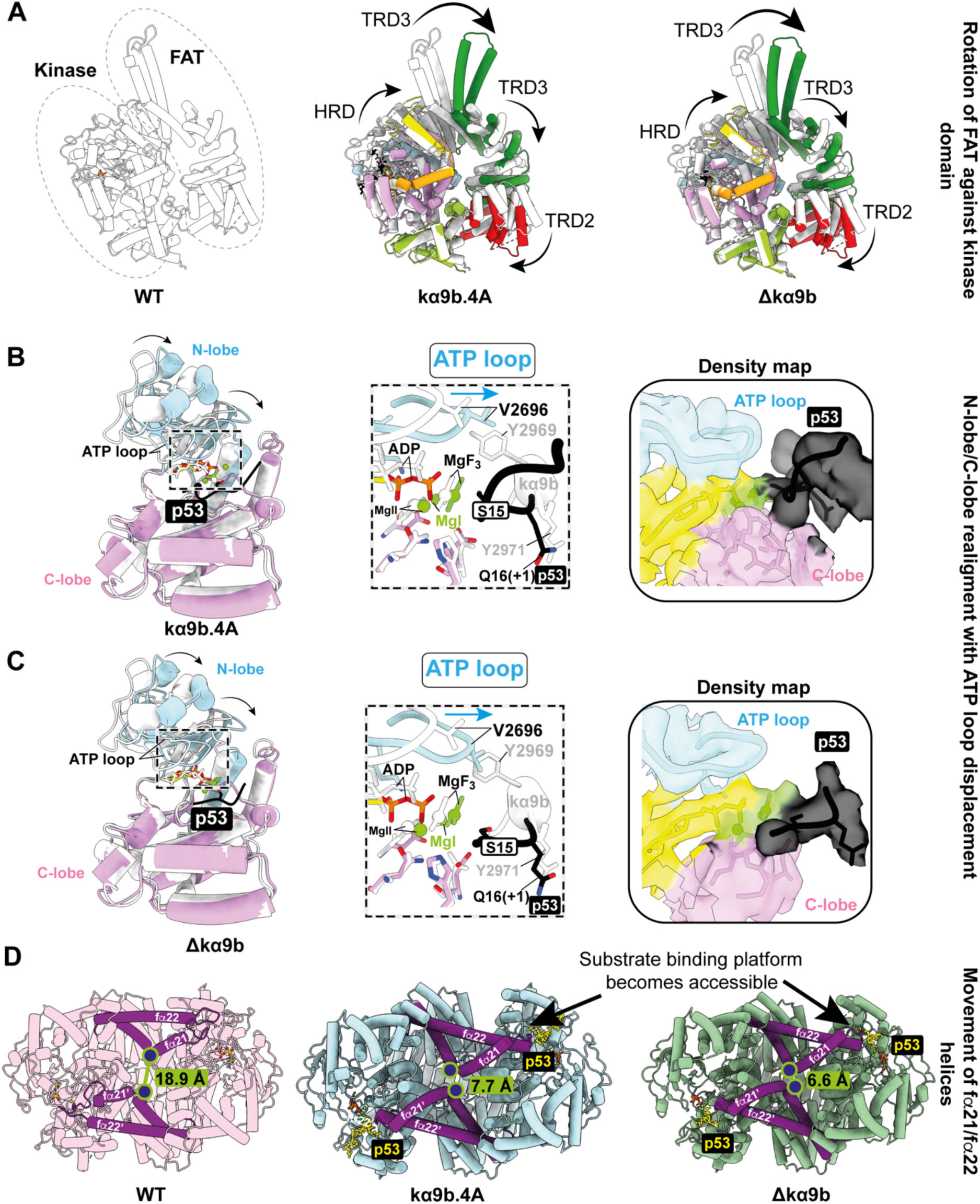
TS structures of the PRD variants are in the active state conformation. The TS structures of kα9b.4A and Δkα9b show distinct global movements relative to TS structure of the WT ATM. (A) Rotation of the FAT TRD2/TRD3/HRD domains relative to the C-lobe of the kinase domain. WT ATM (white) superimposed on the C-lobes of the kinase domains of kα9b.4A and Δkα9b (colored as in the bar diagram of Fig. 1A). (B, C) Kinase domain N-lobe realignment relative to the C-lobe, and large ATP loop movement for kα9b.4A (B) and Δkα9b (C). Structures of the mutants (N- and C-lobe colored blue and pink, respectively) are superimposed on the C-lobe of the WT ATM (white). Middle panels show large movement of the ATP loop, with ADP shown as yellow, MgF_3_^-^ and Mg^2+^ ions light green, and p53^12^ peptide black. Right panels show cryo-EM density for the structural elements shown in the middle inset. These movements are the same as the hallmark features of oxidation (H_2_O_2_)-triggered active-state conformations of ATM (8OXM) (fig. S15). (D) Rotation of the two protomers relative to each other, which brings the two fα21/fα22 elbows (purple) closer together in the kα9b.4A and Δkα9b mutants. Large changes in conformation of the basal (WT) and activated-state (kα9b.4A and Δkα9b) structures are highlighted by showing the distance between symmetry-related Ser^2407^ residue pair (blue sphere).

The same set of global conformational changes were observed previously for H_2_O_2_-activated ATM (with a Q2971A mutation) bound to p53 (fig. S15) (PDB ID 8OXM) (*10*), implying that the conformations of the PRD mutants bound to ADP/MgF_3_^-^/p53 are active-state conformations with the p53^12^ peptide bound. This similarity to the active-state ATM is apparent in a quantitative comparison of the kα9b mutant/TS complexes with the basal WT ATM bound to AMP-PNP/Mg^2+^ (PDB ID 7SIC) (*7*) and with the H_2_O_2_-activated active ATM (Q2971A) bound to AMP-PNP/Mg^2+^/p53^12^ (PDB ID 8OXM) (*10*) (fig. S15). The mutant TS structures differ greatly from the ATM’s basal/inactive conformation by rotation around three virtual hinges (table S2). Upon alignment on the C-lobe, the N-lobe showed a deviation of around ten degrees (∼10°) for the kα9b mutants relative to the WT basal ATM (PDB 7SIC), whereas this deviation for the WT/TS complex is within one degree (∼1°) (table S2). Furthermore, the PRD mutants have a protomer/protomer relationship that differs by ∼13° for the kα9b mutants but by less than a degree (< 1°) for the WT/TS complex with respect to the arrangements for the basal ATM (PDB ID 7SIC) (*7*). The third hinge area is in between the TRD1 and FAT-clamp (table S2), with the FAT-clamp of the kα9b mutants displaying rotations of ∼13° with respect to TRD1, whereas for the WT the rotation is less than a degree (< 1°).

### Complete removal of kα9b facilitates the basal-to-active switch

In the presence of the TS mimic and p53^12^ substrate peptide, Δkα9b was captured only in its active state with ordered, bound TS mimic and p53 (Figs. 2, 3 and figs. S13 and S14). In contrast, kα9b.4A was captured in two major conformations: an active state with bound TS and p53^12^ (Fig. 2 and figs. S8 and S16, Class 1) and a basal state with bound TS only, and no p53 (figs. S8 and S16, Class 2). These results are consistent with our in vitro assays that showed that Δkα9b has higher basal activity than the kα9b.4A. This suggests that Δkα9b is more stable in the active-state conformation than the kα9b.4A. Consequently, we chose Δkα9b to further investigate its structural plasticity in the presence of p53 substrate peptide and other nucleotides.

### Active-site nucleotide dictates the global conformational state of Δkα9b

We determined structures of Δkα9b in the presence of p53^12^ and either substrate or product nucleotides, i.e., ATP or ADP. In the presence of p53 peptide and ATP, Δkα9b took on the active-state conformation, while maintaining its dimeric state (Fig. 4 and figs. S17-S20). Extensive focused 3D classification of ATM particles, followed by local refinement of its FATKIN revealed two conformations of the p53^12^ substrate bound to the Δkα9b: class 1 with the p53^12^ phospho-acceptor Ser^15^ pointing towards the active-site ATP and a second class 2 where Ser^15^ appeared to point away from the active-site ADP (formed in situ from ATP) (Fig. 4 and fig. S20), plausibly representing a post-catalytic state of the p53 peptide. Next, we determined the conformational state of Δkα9b/p53 in the presence of ADP (Fig. 4 and figs. S21-S22). This structure also captured Δkα9b/p53 in the active-state conformation, just as when the ADP was formed in situ from ATP hydrolysis.

**Fig. 4.**
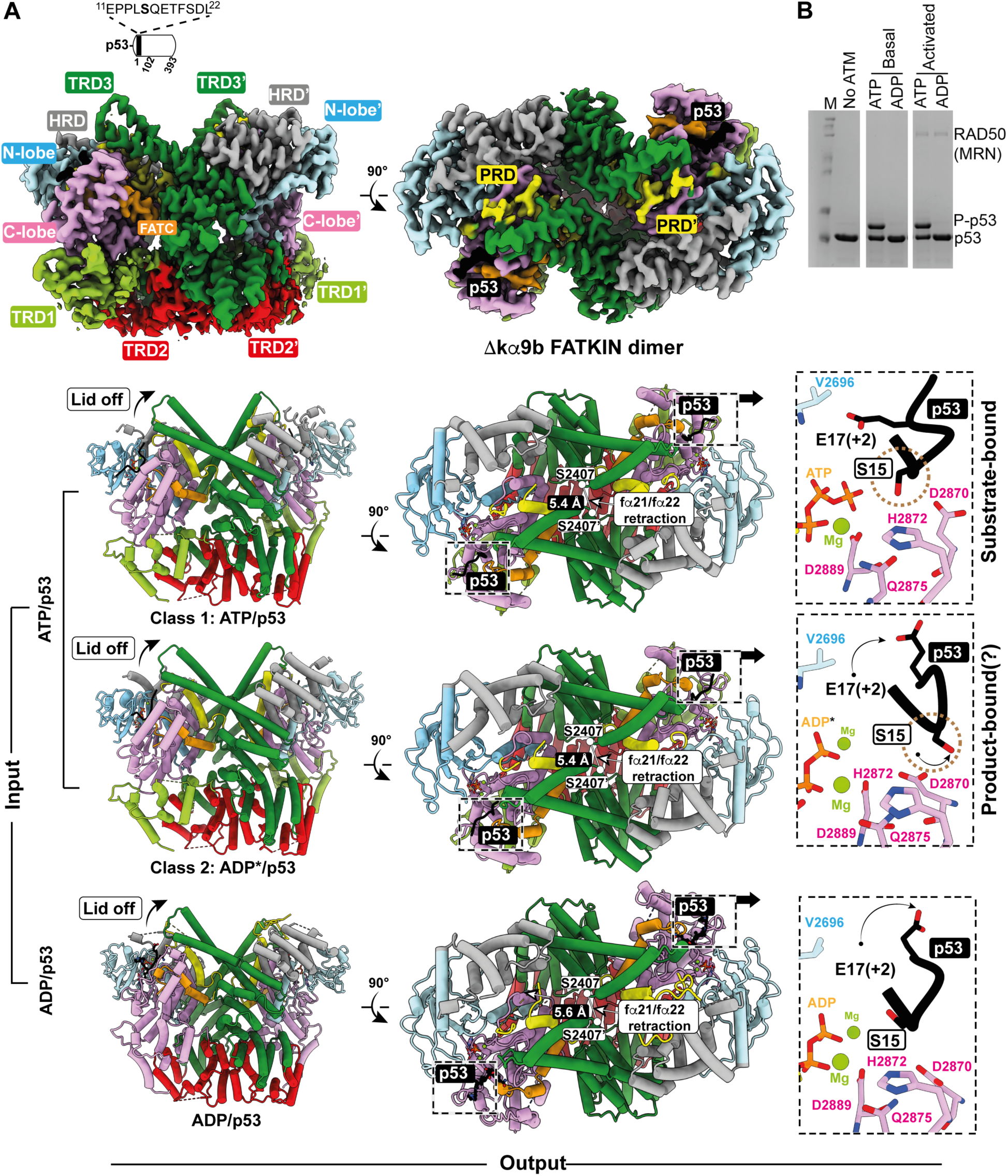
Nucleotide-dependent conformational plasticity of the ATM PRD mutant Δkα9b. (A) Cryo-EM density map of the ATM PRD mutant Δkα9b (left panels, two orthogonal views), with domains colored as in Fig. 1A. Also shown is the sequence of the p53^12^ substrate peptide used for cryo-EM sample preparations. Upper and middle of the bottom three panels show side and top views of a model of Δkα9b obtained from Δkα9b sample in the presence of ATP/p53^12^. There were two cryo-EM classes: ATP/p53^12^ bound to Δkα9b (class 1, upper three panels) and ADP/p53^12^ where ADP was formed in situ from ATP (ADP*) (class 2, middle three panels). The lower of the bottom three panels shows a model of Δkα9b/ADP/p53^12^ obtained from a cryo-EM sample of Δkα9b treated with ADP and p53^12^, showing ADP/p53^12^ bound at the active site. All three structures show helices fα21/fα22 being closer to the dimer interface relative to WT ATM (Fig. 3D, left panel) characteristic of activated conformations. Insets to the right of the bottom three panels show the conformation of p53^12^ substrate peptide (black) in each of the three Δkα9b structures. (B) PhosTag gel analysis of kinase activity of Δkα9b on p53^102^, conducted in the presence or absence of MRN/DNA and with ATP or, as a control, ADP. Only in the presence of ATP is there phosphorylation of p53^102^ (with and without MRN/DNA).

### ATP binding alone can transform the Δkα9b mutant from inactive to active state

Since none of our basal-state cryo-EM structures had the p53 substrate peptide bound (WT ATM, Fig. 2; and kα9b.4A, figs. S8 and S16), we explored whether substrate peptide is necessary for Δkα9b to form a stable active-state conformation or whether nucleotide alone, without p53 substrate, can stabilize it in its active-state conformation. When we treated the Δkα9b mutant with ATP/Mg^2+^ alone, we observed that Δkα9b took on an active-state conformation with ATP bound (Fig. 5, A and B and figs. S23-S24). To see if ATP/Mg^2+^ has a similar effect on kα9b.4A, the other hyperactive mutant of ATM, we also determined its structure in the presence of ATP/Mg^2+^. Like Δkα9b, kα9b.4A also showed bound ATP (Fig. 5, A and B and figs. S25-S26). However, instead of switching to the active-state, the kα9b.4A maintained a homodimeric basal-state conformation. This indicates that peptide substrate binding is critical for the active-state conformation of kα9b.4A but not for Δkα9b. For Δkα9b to adopt its active-state conformation, ATP/Mg^+2^ binding alone was sufficient. Like kα9b.4A, no conformational switch was observed for WT ATM binding to ATP/Mg^+2^ (Fig. 5, A and B and figs. S27-S28). This implies that WT ATM requires assistance from an additional activating influence (oxidation or MRN-DNA) for its transformation to an active conformation (Figs. 1, 2, and 5) (*7, 10*).

**Fig. 5.**
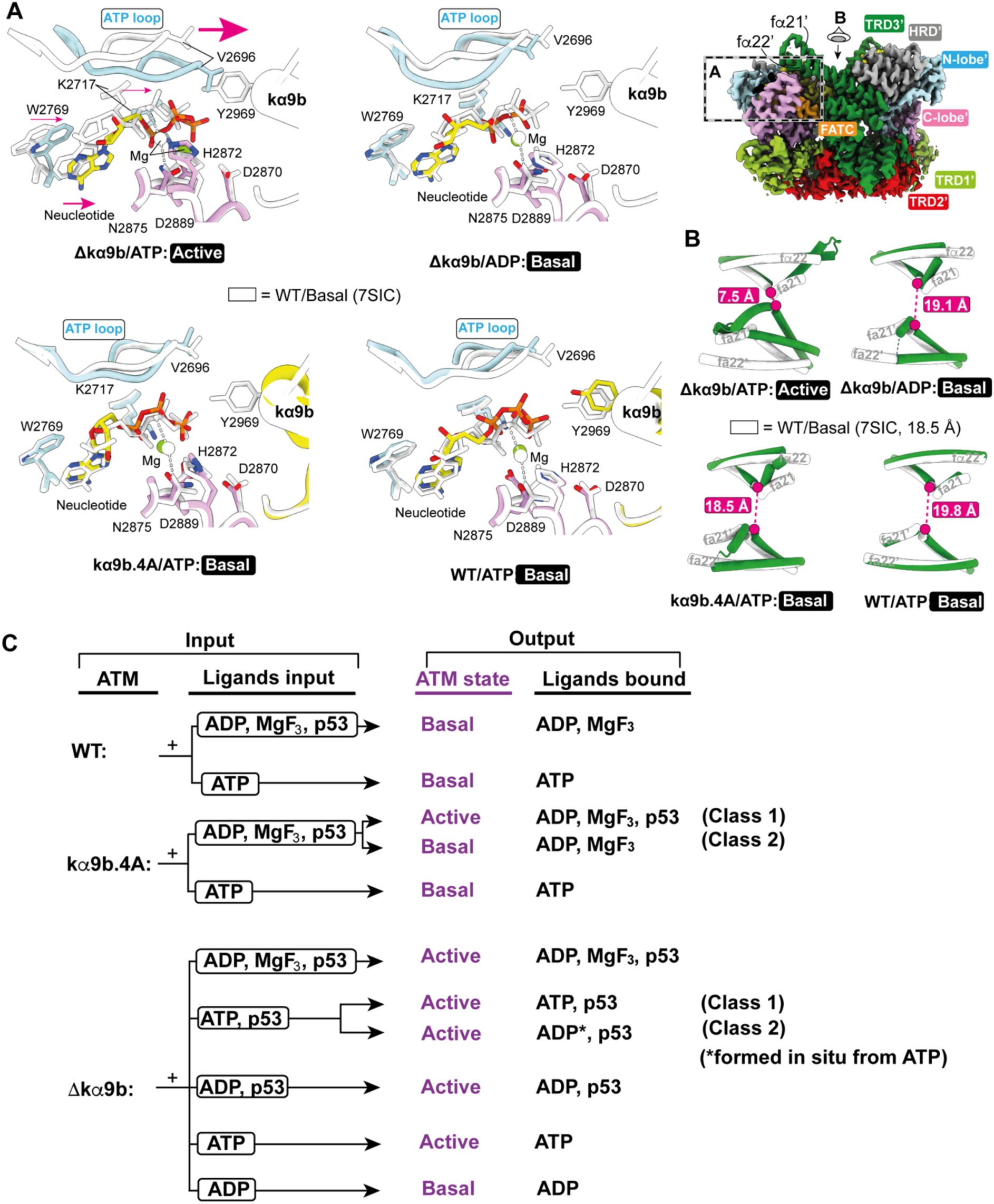
ATP binding alone, but not ADP, can trigger a conformational switch for Δkα9b. (A) Cryo-EM density map of the Δkα9b mutant (upper right), with domains colored as in Fig. 1A, indicating views shown in the rest of the panels. Model of Δkα9b (colored) bound to ATP alone (without p53 substrate peptide) (upper left panel) or ADP alone (upper middle panel), superimposed on the C-lobe of the WT/AMP-PNP ATM (7SIC, white). Only Δkα9b/ATP, but not Δkα9b/ADP, shows ATP-loop movement characteristic of the active-state ATM. Models of kα9b.4A mutant (colored) bound to ATP alone (bottom left panel) or of WT ATM (colored) bound to ATP alone (bottom middle panel), superimposed on the C-lobe of the WT/AMP-PNP ATM (7SIC, white). Both kα9b.4A/ATP and WT/ATP maintain their basal state conformation. (B) Distances between the fα21/fα22 elbows at the dimer interface for the Δkα9b/ATP, Δkα9b/ADP, kα9b.4A/ATP and WT/ATP, as indicated. All four structures are shown in dark green, superimposed on basal WT/AMP-PNP ATM (7SIC, white). Only Δkα9b/ATP has fα21/fα22 elbows close together, characteristic of the active-state ATM (see Figs 3 and 4). (C) ATM variants and the ligands added to ATM before cryo-EM grid preparations are indicated as “Input”. Conformation of ATM, as well as the bound ligands observed in cryo-EM structures are indicated as ‘Output”. Where a single input shows multiple outputs, each output is a cryo-EM class that has a unique bound ligand and/or ATM conformational state (basal or active). All structures that have bound p53 peptide substrate are in the active-state conformation.

On the other hand, when we used ADP/Mg^2+^ (without p53 peptide), Δkα9b took on the basal-state conformation (Fig. 5, A and B and figs. S29-S30), implying that interactions with the γ-phosphate of ATP or interactions from the substrate peptide are essential for the conformational switch. Altogether, our cryo-EM structures suggest that the dimer is poised for activation but does not actually switch to the activated state, unless it is bound to ligands that are conducive to the activated state, such as ATP (not an ATP mimic) along with p53 substrate (Fig. 5C).

### Hyperactivation of Δkα9b is not due to autophosphorylation of PRD residues

Previous reports of post-translational modifications and cancer-associated mutations of the PRD linker implicated the PRD as having a role in ATM activation: PRD residue Cys^2991^ forms an intermolecular disulfide bond that is essential for ROS-induced ATM activation (*9, 10*); Ser^2996^ phosphorylation and Lys^3016^ acetylation are important for DNA damage-induced ATM activation (*31–34*); Arg^3008^ of kα10 is the site of the most common missense cancer mutation (*35*), and the R3008H mutation results in reduced ATM activity and abrogates DNA damage- and oxidative stress-induced activation of ATM (*36*).

Our PhosTag gels revealed that Δkα9b, as purified from human cells is hyperphosphorylated relative to the WT enzyme (fig. S31A). To determine whether Δkα9b phosphorylation contributed to its significantly higher basal activity, we carried out mass spectrometry, which revealed that Ser^3001^ (corresponding to mouse Ser^3021^, figs. S31B, E) of the PRD linker (a 3000-QS-3001 motif) is autophosphorylated in Δkα9b (fig. S32). We mutated Ser^3001^ to alanine, together with Ser^2996^ (Δkα9b^S2A2^) (fig. S31, B and E), which is also in the PRD linker and was reported previously to be phosphorylated. However, the kinase activity of Δkα9b^S2A2^ was comparable to that of Δkα9b (fig. S31C). In addition, a global dephosphorylation of Δkα9b using λ-protein phosphatase (λ-PP) resulted in only about ∼2.0-fold reduction in kinase activity (figs. S31D, S33). Auto re-phosphorylation gained back the original activity (fig. S31D). This suggests that autophosphorylation of the PRD linker accompanies ATM activation but is not responsible for hyperactivation of Δkα9b.

## Discussion

To our knowledge, this is the first time that human ATM has been reported to exhibit constitutive, MRN complex-independent activation in cells. Our structures show concerted conformational transitions that are core to ATM activation (fig. S34): displacement of the PRD from the substrate-binding pocket, realignment of the kinase N-lobe relative to the C-lobe, and sliding of the FAT clamp. These transitions are shared between PRD mutation-driven and oxidative stress-driven activation (*10*), and are closely analogous to the consensus conformational changes observed across the PIKK family upon activation, including mTORC1 activated by RHEB (*18*), Mec1 activated by Ddc2 (*23*), and DNA-PK activated by a nucleotide mimic (*21, 37*) (fig. S35).

PRD mutants that are constitutively active retain robust MRN binding and indeed they bind MRN more efficiently than WT ATM, yet they are not appreciably stimulated by MRN/DNA in vitro or in *ATM/TP53* double-knockout human RPE1 cells. This might be because the mutants are already in a conformation that closely resembles the MRN/DNA-activated conformation, so that binding energy is not dissipated in conformational rearrangement. However, it is also possible that mutations have a unique conformation that coincidentally results in full activation and tighter binding to the MRN/DNA, without the mutants mimicking the natural conformation of the WT enzyme when activated my MRN/DNA. An observation that is consistent with the PRD mutants partially mimicking the activated-WT conformation is a recent cryo-EM study of fission yeast Tel1/ATM from cells that had been pre-treated with methylmethane sulfonate (MMS) to form DSBs. This study captured an asymmetric Tel1 dimer in which one protomer had a fully ejected PRD with re-aligned N- and C-lobes competent for nucleotide and substrate binding, while the other protomer retained a partially disordered PRD and lacked bound peptide (*38*). The authors speculated that after being exposed to DSBs in MMS-treated cells, the Tel1^ATM^ remained active throughout a robust purification protocol, because the activated Tel1^ATM^ retained uncharacterized post-translational modifications, such as phosphorylation and acetylation. In addition to the MMS-treated WT Tel1^ATM^ sharing some structural signatures of activation in common with the PRD mutants, MMS-treated Tel1^ATM^ is persistently active and, like the PRD kα9b variants of ATM, cannot be further activated by MRN/DNA. Nevertheless, a direct structural characterization of ATM in a complex with MRN/DNA will be required to reveal the precise sequence of events leading to MRN/DNA mediated active-site remodeling of ATM. It may be that post-translational modifications in cells lock the active state once the MRN/DNA triggers a switch to the active conformation. There may be additional interactions that MRN/DNA make with ATM that do not activate ATM’s maximal catalytic activity, but which are necessary for recruitment of ATM to sites of DNA damage in cells (*39*).

Our data suggest that both WT and PRD mutants of ATM exist in equilibrium between basal and active conformations (*movie clip 5)*, but the PRD mutants appear to have a significantly lower barrier to switching to an active state. Mutation or deletion of the PRD on its own does not automatically elicit the active state conformation: for the kα9b.4A mutant, both reactants (ATP and peptide substrate) are needed to shift the enzyme to the active state, while Δkα9b (complete deletion of kα9b) needs only ATP to assume the active conformation (Fig. 5). Basal conformations for all ATM structures in this paper are globally similar, but some details differ. Almost half of the fα21/fα22 coiled coil is disordered in the basal conformation of kα9b.4A and Δkα9b (the apical half, which in the WT interacts with the PRD) (Fig. 5 and fig. S16). This disorder suggests that in the basal state of WT ATM, interaction between the kα9b and the apical end of fα21/fα22 mutually stabilize each other, and deletion of kα9b removes this stabilizing effect on fα21/fα22. In addition, disorder of the apical end of fα21/fα22 is not sufficient for ATM to switch to the active state, as shown by the basal conformation of Δkα9b/ADP with disordered apical half of fα21/fα22 (Fig. 5B). This is consistent with the observation that deletion of the apical half of fα21/fα22 (dDH, residues 2408-2450 deleted) maintains the same low basal activity as WT ATM (*40*).

The activated conformations for all ATM structures described here are also globally very similar. Probably the most distinct difference among the active-state structures is the presence of two ordered Mg^2+^ ions in the active-state conformations of Δkα9b and kα9b.4A mutants prepared with the p53 substrate and transition state mimic (ADP/MgF_3_^-^), while only a single Mg^2+^ ion is observed in all other active-state structures of the PRD mutants as well as in all previously published structures of ATM (*7, 10, 28, 34*). In budding yeast Mec1, binding of a second Mg^2+^ ion, which was observed even in the absence of protein substrate, was greatly facilitated by the presence of a second Asp in its DFD motif at the beginning of the activation loop, instead of the DLG motif found in ATM and the DFG motif found in most protein kinases (*23*). For many protein kinases, including CDK2, binding of the second Mg^2+^ is transient but is necessary for the flexible Gly-rich ATP-binding loop to close onto the ATP and for the already activated phospho-CDK2/Cyclin complex to achieve maximal enzyme velocity (*25*).

The hyperactive ATM mutants we have studied helped identify structural changes and ligands required for the mutants to transition from a basal to an active conformation (Fig. 5C). Both hypo- and hyper-activation of ATM are linked to aberrant cell states (*4, 11, 13, 41*). Differential binding of small molecules to constitutively activated mutants compared with the basal state could provide a starting point for developing allosteric ATM activators. The conformational changes that we see associated with ATM activation may be components of ATM activation in biological processes such as DNA damage signaling, senescence, and telomere length regulation. Hyperactivation of the fission yeast Tel1^ATM^, caused by its overexpression, leads to prolonged delays in cell cycle progression, even without exogenous DNA damage, due to persistently increased phosphorylation of Tel1^ATM^ targets such as Rad53 (CHK2 homologue) (*42*). The presence of genotoxic insults exacerbates this cell cycle slowing and can lead to cell death (*42*). Similarly to overexpressed Tel1^ATM^, hyperactive Tel1^ATM^ mutants partially suppress the checkpoint defects of Mec1-deficient cells (Mec1 is the budding yeast ATR orthologue) and hypersensitivity to genotoxic stress (*42–44*). The ability of the upregulated Tel1^ATM^ mutants to phosphorylate checkpoint targets was dependent on the MRX (MRE11-RAD50-XRS2) complex and required MRX complex for recruitment to the DNA damage sites (*44*).

We observe across different cell lines that the expression of activated ATM mutants promotes hyperphosphorylation of a broad range of ATM substrates. Notably, this hyperactivation occurs constitutively, in the absence of DNA damage and does not further increase upon IR-induced DNA damage. This suggests that the ATM activation state reaches saturation in untreated conditions. Furthermore, we have found that MRE11 and NBS1 depletion does not prevent this hyperactivation. This observation is consistent with our cell-free data and indicates that the hyperactive ATM mutants acquire conformations that bypass the requirement for MRN complex for their activation. The structures of the PRD mutant variants show that distinct conformational events accompany ATM activation: PRD ejection, FAT-clamp sliding, N-lobe/C-lobe realignment and fα21/fα22 coiled-coil retraction. In cells, that have high concentration of ATP, PRD ejection alone is likely to be sufficient to drive the full active-state conformational program within a dimeric ATM. It is plausible that the basal-to-active state transition of WT ATM could be initiated by one of these events, with the rest following automatically.

## Acknowledgements

We thank Grigory Sharov of the LMB Cryo-EM facility for support with *cryoSPARC* software installation and maintenance; the LMB EM facility staff for access and support with sample preparation and data collection; Jake Grimmett, Toby Darling, and Ivan Clayson for maintaining the LMB’s scientific computing facility; the LMB Mass Spectrometry facility for analyzing ATM samples; Chris Batters and Stephen McLaughlin of LMB Biophysics facility; LMB media kitchen for providing buffers and media; and the Williams lab members for sharing feedback.

## Author contributions

Conceptualization: O.P., R.L.W., S.P.J., and M.S.I.

Methodology: M.S.I., V. G. G., R.B., O.P., A.S.B.

Investigation: M.S.I., V.G.G., D.B., O.P., and A.S.B.

Visualization: M.S.I., V.G.G., R.B., A.S.B., and O.P.

Data acquisition: M.S.I. and V.G.G.

Data processing: M.S.I., V.G.G, and A.S.B.

Data deposition: M.S.I.

Funding acquisition: S.P.J. and R.L.W.

Supervision: S.P.J., and R.L.W.

Writing – original draft: M.S.I.

Writing – review & editing: M.S.I., O.P., R.B., V.G.G., A.S.B., D.B., S.P.J., and R.L.W.

## Data and materials availability

All plasmids are available from R.L.W.

PDB coordinates are deposited with the PDB under accession numbers 9U0Q, 9TPV, 9TY6, 9TY7, 9TY8, 9TY9, 9TYA, 9TYB, 28XP, 28XQ, 28XS, 28XT, 28YT, 28YU, 29CG, 29CH, 29CQ, 29CR, 29EZ, 29FA, 29FT, 29FU,

Cryo-EM maps are deposited with the Electron Microscopy Database (EMD) under accession numbers EMD-56499, EMD-56116, EMD-56412, EMD-56413, EMD-56414, EMD-56415, EMD-56416, EMD-56417, EMD-56944, EMD-56945, EMD-56947, EMD-56948, EMD-56984, EMD-56985, EMD-57077, EMD-57078, EMD-57081, EMD-57082, EMD-57134, EMD-57135, EMD-57137, EMD-57138.

