## Supplemental Materials for "Insights into mechanisms of ATM activation via constitutively active mutants"

**The PDF file includes:**

Materials and Methods

Figs. S1 to S35

Tables S1 to S7

**Other Supplementary Materials for this manuscript include the following:**

Movies S1 to S5

**MATERIALS AND METHODS**

Cloning, expression, and purification of the ATM variants

Codon-optimized human full-length *ATM* (aa 1-3056, UniProt Q13315) was cloned with an N-terminal FLAG sequence into pDEST12.2-OriP expression vector (*14*). A DNA fragment containing FKBP1A (UniProt P62942) followed by a nine amino acid linker (GGSAGSGSAS) was inserted between the FLAG and ATM sequences using NEBuilder DNA HiFi assembly (New England Biolabs). The ATM mutants were generated using a fragment-assembly based approach. Briefly, two fragments were PCR amplified with overlaps containing either a desired mutation or deletion at one end and overlaps with the digested FKBP-ATM plasmid at the other, so that the combined fragments covered the entire ATM sequence. These fragments were assembled using HiFi DNA assembly.

ATM variants were expressed in mammalian Expi293 cells by following the procedure described previously(*14*). Briefly, the Expi293 cells were grown in Expi293 Expression Media at 37°C, 125 rpm and 8% CO_2_ in 5L Corning flasks. At a density of 2.5 x 10^6^ cells/mL, cells were transfected with DNA and PEI (1.1 mg DNA and 3 mg/mL PEI per 1 L cell culture) and grown for another 50 h, followed by harvesting with 3000 g centrifugation at 4°C for 15 min. Harvested cells were flash frozen in liquid nitrogen and stored at -80°C until further use.

Cell pellets were resuspended in Lysis buffer (50 mM HEPES pH 7.5, 100 mM L-arginine, 150 mM NaCl, 10% glycerol), supplemented with EDTA-free protease cocktail inhibitor tablets (Roche 05056489001), universal nuclease (Cat. 88702, Thermo Fischer), and protease inhibitor PEFA (Cat. A20010, Melford) (*10, 14*). The cells were disrupted by sonication with 10-s on/10-s off for a total of 5 minutes at a 40% amplitude. Next, the cell lysates were cleared by centrifugation in a Ti45 rotor at 4°C, 88000 g for 30 min. ATM was purified in two steps. First, the lysates were incubated with FLAG M2 affinity beads (A2220, Merck) equilibrated in buffer A (50 mM HEPES pH 7.5, 100 mM L-arginine, 150 mM NaCl, and 10% glycerol) for 1 h at 4°C on a roller, followed by elution in Elution buffer (50 mM HEPES pH 7.5, 150 mM NaCl, 10% glycerol) supplemented with 250 µg/mL FLAG peptide (F4799, Sigma-Aldrich). The presence of ATM proteins in the eluted fractions was confirmed by SDS-PAGE analysis. Next, the desired fractions were pooled together, concentrated using a 100 MWCO centrifuge filter (Amicon Ultra, 100K), and purified in a pre-equilibrated Superose 6 16/600 size-exclusion column in Gel-filtration buffer (50 mM HEPES pH 7.5, 150 mM NaCl, 10% glycerol, and 2 mM TCEP). Purified proteins were concentrated using 100 MWCO centrifuge filter, flash frozen in liquid nitrogen, and stored at -80°C.

For cell culture studies, plasmids containing *ATM* variants were diluted in OptiMEM, left for 10 min, and mixed with OptiMEM-diluted TransLT transfection reagent, further incubated for 20 min, then added dropwise to HEK 293T cells at a 70-80% confluency. At 48 h post-transfection, experiments were performed and expression was typically assessed as indicated by Western blotting.

Expression and purification of human MRN complex

Human MRN complex (MRE11, NBS1, and RAD50) was cloned into insect cell expression vectors using the HiFi DNA assembly (10). Briefly, human RAD50 with a C-terminal twin Strep-tag II (TEV cleavable) was cloned into pFastBac, while human NBS1 with a C-terminal FRB domain followed by a twin Strep-tag II (TEV cleavable) and human MRE11 (no tag) were cloned into pFastBacDual expression vector. Expressions were carried out in Sf9 insect cells grown in Sf-900 II SFM medium. Cells were harvested 48 h post-transfection by centrifugation (4000g, 8 min, 4°C), pellets were flash frozen in liquid nitrogen, and the pellets were stored at -80°C until further use.

The MRN complex was purified by following a procedure described previously (10). Briefly, cell pellet from a 2 L culture was resuspended in the Lysis buffer (25 mM Tris pH 8.5, 300 mM NaCl, 100 mM arginine mono-HCl, and 10% glycerol) supplemented with EDTA-free protease cocktail inhibitor tablets (Rosche 05056489001, Universal Nuclease (Thermo Fischer Scientific, 88702), and Pefabloc (Melford, A20010). The cells were disrupted by sonication at 4°C for a total of 2 min 15 sec, with 3-s on/8-s off, at an amplitude of 60%. Cell lysates were cleared by centrifugation at 88,000g for 30 min at 4°C in a Ti45 rotor. The supernatant was further clarified by passing it through a 5 μm syringe filter and loaded onto 3x5 ml StrepTrap columns previously equilibrated with the Binding buffer (25 mM Tris pH 8.5, 300 mM NaCl, 100 mM arginine mono-HCl, and 10% glycerol). After a thorough washing of the columns using the Binding buffer, MRN complex was eluted in 6x15 ml of StrepTrap Elution buffer (25 mM Tris pH 8.5, 150 mM NaCl, 10% glycerol, 10 mM desthiobiotin, and 0.5 mM TCEP). The pooled fractions were diluted in Q Dilution buffer (25 mM Tris pH 8.5, 10% glycerol, and 0.5 mM TCEP) to bring the NaCl concentration to 100 mM, and the diluted sample was loaded onto a 5 ml Q HP column previously equilibrated with the Q Buffer A (25 mM Tris pH 8.5, 100 mM NaCl, 10% glycerol, and 0.5 mM TCEP). After a thorough washing of the Q column using the Q Buffer A, the MRN complex was eluted with a NaCl gradient. Elution fractions containing MRN complex were pooled together, concentrated to 1.5 ml using 15 ml 100 kDa MWCO filter concentrator (Amicon® Ultra-15, Merck Millipore Ltd.), and loaded onto a HiLoad Superose 6 16/600 size-exclusion column (GE Healthcare) previously equilibrated with MRN gel filtration buffer (50 mM HEPES pH 7.5, 300 mM NaCl, 10% glycerol, and 0.5 mM TCEP). Size-exclusion fractions containing MRN complex were pooled, concentrated to 10-15 mg/ml, flash-frozen in liquid nitrogen, and stored at -80°C.

Cloning, expression and purification of p53 protein

A *TP53* gene fragment encoding the N-terminal 1-102 residues (p53^1-102^) was PCR amplified from a human *TP53* construct. The amplified *TP53^1-102^* gene was inserted into a NdeI/BamHI-cut pOPTHM expression vector, with an N-terminal *His_6_-*tag, Maltose-binding protein (MBP), and a TEV cleavage site.

The His-MBP-TEV-p53^1-102^ plasmid was grown in 2xTY media supplemented with 100 µg/mL of ampicillin at 37°C and 200 rpm. Cells were induced with 0.5 mM IPTG at an OD_600_ of 0.8 to start protein expression and continued to grow for another 3 h. Cells were harvested by centrifugation at 6000 g and 4°C, flash frozen in liquid nitrogen, and stored at -80°C until further use.

Cell pellets were resuspended in the Lysis buffer (50 mM HEPES pH 7.5, 150 mM NaCl, 10 mM imidazole, and 10 mM glycerol) supplemented with EDTA free protease inhibitor cocktail tablet (Roche 05056489001), Pierce Universal Nuclease (Thermo Fischer 88702), and PEFA protease inhibitor (Melford A20010). Cells were lysed by sonication for 5 min with 10 s on/off at an amplitude of 60%. Lysed cells were cleared using ultracentrifugation at 4°C for 30 min at 35000 g and the supernatant was further clarified using a 5 µm syringe filter. The lysates were loaded onto 5 mL HisTrap IMAC column equilibrated in buffer A (50 mM HEPES pH 7.5, 150 mM NaCl, 10 mM imidazole, and 10 mM glycerol). Next, the column was washed rigorously with buffer A, followed by elution of p53 in a gradient from buffer A to Elution buffer (buffer A + 1 M imidazole). The eluted fractions were confirmed for His-MBP-TEV-p53 by SDS-PAGE and the desired fractions were pooled. The p53 protein was further purified using a HiTrapQ anion exchange column.

Substrate peptide

An N-terminal 12-residue long peptide of human p53 protein (UniProt P04637) spanning the ATM phosphorylation site Ser^15^ (p53^12^: ^11^EPPL**S**QETFSDL^22^) was purchased from Bankpeptide, China. The peptide was custom synthesized with an N-terminal NH_2_ and C-terminal COOH, in acetate salt, with trifluoroacetic acid removed. The lyophilized powder was dissolved to 100 mM in 50 mM HEPES pH 7.5 and stored at -80°C.

Kinase assays

For the basal kinase reaction (ATM alone), 1000 nM WT ATM (10 nM for the hyperactive mutants, kα9b.4A and Δkα9b) was incubated with 25 μM MBP-p53^1-102^ in the presence of 1 mM ATP and 5 mM MgCl_2_. For MRN/DNA-activated conditions, human MRN complex was expressed in Sf9 insect cells and purified as described previously. MRN complex (final concentration 300 nM) and dsDNA (350 bp fragment, final concentration 25 nM) were added to ATM (final concentration 10 nM for either WT or PRD mutants), 1 mM ATP and 5 mM MgCl_2_. All the kinase assays were run at 30°C on a PCR block thermocycler.

PhosTag gel assays

Phos-tag gels were either cast in-house or obtained as commercially pre-cast gels (FUJIFILM 192-18001) (7.5% polyacrylamide, 17-well, 5 µmol/L phos-tag reagent). A protocol from the Wako Phos-tag™ SDS-PAGE was followed to cast in-house. Briefly, the resolving gel was made by gently mixing 7.5% acrylamide/bis (29:1, 3.3% crosslinker), 350 mM Bis-Tris/HCl (pH 6.8), 50 µM Phos-tag reagent (APExBIO F4002), 100 µM ZnCl_2_, 0.1% TEMED (v/v) and 0.05% ammonium persulfate (APS) (w/v) in a 50 mL falcon tube. The solution (~ 6 mL) was then gently pipetted into the space between two glass plates (83 x 100 x 3.9 mm) that were separated by waterproof spacers and sealed together on three sides by waterproof tape. The resolving gel was left to set with a 100% ethanol overlay. After ~ 60 min, the ethanol was decanted carefully. 4.5% polyacrylamide stacking gel solution was prepared by mixing 4.5% acrylamide/bis (29:1, 3.3% crosslinker), 350 mM Bis-Tris/HCl (pH 6.8), 0.1% TEMED (v/v) and 0.05% APS (w/v) in a 50 mL falcon tube. Approximately 2 mL of stacking gel solution was gently pipetted on top of the resolving gel in the cassette. Gel combs (washed and recycled from commercial gels), were placed into the stacking gel solution soon after pipetting and the gel was left to set for > 60 min before use. Gels were run with 1xMOPS buffer without EDTA for 1 h at 100 V constant voltage and for another 1 h at 150 V constant voltage.

Determination of kinetic parameters

ATM kinase activity was measured by monitoring phosphorylation of the MBP-p53^102^ substrate. Phosphorylated and non-phosphorylated substrate band intensities from Phos-Tag gels were quantified using Bio-Rad Laboratories Image Lab version 6.0.1. Initial rates were calculated by measuring the ratio of phosphorylated substrate to non-phosphorylated substrate, then calculating the amount of substrate phosphorylation at four different timepoints in the linear range and adjusting for the enzyme concentration. Kinetic parameters *k*_cat_ and *K*_M_ were calculated using Prism 11 assuming Michaelis-Menten kinetics.

ATM/MRN binding assay

An in vitro binding assay was conducted to assess the binding of ATM variants to the MRN complex in the presence of DNA. Briefly, 2 μM Flag-tagged ATM (WT, kα9b.4A, and Δkα9b) was incubated with anti-Flag M2-affinity bead suspension (50%, ANTI-FLAG® M2 Affinity Gel, Sigma-Aldrich, A2220) in the presence of 2.5 mM ATP and 5 mM MgCl_2_. Unbound ATM was separated from the bead-bound ATM through centrifugation. The bead-bound ATM was then incubated with MRN/DNA complex (5 μM MRN complex and 2 μM 200 bp dsDNA (IDT Ultrapure)) formed previously in the same buffer containing ATP and MgCl_2_. The ATM/MRN/DNA mixture was washed in buffer to separate the complex from the unbound components, followed by an elution of the Flag-ATM from the anti-FLAG beads using 250 ng anti-Flag peptide (3x, F4799, Sigma-Aldrich). Eluted Flag-ATM and the MRN complex bound to it were resolved in a SDS-PAGE protein gel in 1x MOPS buffer at 200 V. A Bio-Rad Laboratories Image Lab (version 6.0.1) gel imager was used to quantify the protein bands, and Prism 11 software package was used to construct the bar diagrams.

Dephosphorylation of Δkα9b

Global dephosphorylation of Δkα9b was performed by incubation of Δkα9b with λ-protein phosphatase (λ-PP, ~1 μg of phosphatase per 16 μg of Δkα9b) in 1x NEB protein metallophosphatase (PMP) buffer supplemented with 1 mM MnCl_2_. To remove λ-PP, the dephosphorylated Δkα9b was then purified by size-exclusion chromatography (SEC) on a Superose 6^TM^ Increase 3.2/300 column equilibrated in ATM SEC buffer (50 mM HEPES pH 7.5, 200 mM NaCl, 10% glycerol, 2 mM TCEP) (fig. S33). Eluted fractions with Δkα9b were pooled, concentrated and stored as for untreated Δkα9b, and later used in kinase assays and LC-MS/MS analysis.

### Cell Culture, Irradiation and siRNA Knockdown

### HEK293 (WT) were cultured in Dulbecco’s Modified Eagle’s Medium (Gibco, Cat. Nr.: A4192101), and RPE1-hTERT Cas9 and RPE1-hTERT Cas9 *ATM⁻/⁻ TP53⁻/⁻* cells (*24*) were cultured in Dulbecco’s Modified Eagle Medium: Nutrient Mixture Ham’s F-12 (DMEM/F-12, Gibco, Cat. Nr.: 11320033), each supplemented with 10% fetal bovine serum (FBS; Thermo Fisher Scientific), 100 mg/mL penicillin and 100 units/mL streptomycin (GIBCO), and L-Glutamine (2 mM). All cell lines were cultured in a humidified incubator (37°C, 5% CO₂), routinely tested for mycoplasma, and no mycoplasma was detected. To irradiate, an X-ray irradiator was used at RT and a dose of 2 Gy. Afterwards, the plates were returned to standard growing conditions for 1 h recovery, and the samples were further processed. For knockdown experiments, small pooled interfering RNA (siLUC control, siMRE11, and siNBS1 (Horizon, Cat.Nr: D-001100-01-0, M-009271-01-0005, and M-009641-02-0005, respectively) were used in a final concentration of 20 nM. RPE1 cells were seeded to be 70-80% confluent at the time of siRNA transfection. The siRNA and Lipofectamine RNAiMAX (Invitrogen, Cat. Nr.: 13778150) were diluted in OptiMEM (Thermo Fisher Scientific, Cat.Nr: 31985-054), incubated for 10 min, mixed, incubated for a further 20 min, and added dropwise to the cells. After 24 h, a second round of transfection was carried out. 48 h post-second transfection cells were seeded for immunofluorescence assays and sampled for assessing knockdown efficiencies by Western blotting.

### Cloning for Lentiviral Production and Protein Expression

The Gateway cloning system (Thermo Fisher Scientific) was used to clone FLAG-WT or ATM variants from plasmids vectors into a pLIX-403 lentiviral vector as described in the manufacturer’s protocol. Successful clones were selected based on restriction enzyme digestion patterns and confirmed by whole-plasmid sequencing (PlasmidSaurus).

For Lentivirus production, HEK 293T LentiX cells (Takara) were seeded to be grown to 70-80% confluency after 24 h in a 10 cm plate. The TransLT transfection reagent (Mirus, Cat.Nr.: MIR 2305) and OptiMEM were incubated for 5 min, prior to adding them to a mix of 9.3 µg of ATM pLIX-403 vectors with 6.1 µg of capsid vector psPAX2 and 1.8 µg envelop vector pMD2.G. After incubating for 10 min, the plasmid-transfection mixture was then added dropwise to HEK 293T LentiX cells. 72 h post-transfection, the viruses were harvested by filtering the media supernatant through a 0.45 µm filter (Merck, Cat.Nr.: HAWG047500). The viruses were used to transduce RPE1-hTERT Cas9 *ATM⁻/⁻ TP53⁻/⁻* cells at MOI <1, and successful transduction was selected with puromycin (Sigma-Aldrich, 1.5 µg/mL). Stable ATM WT or ATM variant expression was induced for 24 h by doxycycline (0.1 ug/mL), to be further processed.

### Immunoblotting

Samples were lysed in SDS-Laemmli buffer (1M Tris-HCl, pH 6.8, 20% SDS, Glycerol) and denatured for 3 min at 95°C. SDS-PAGE was used to separate proteins, which were transferred onto a nitrocellulose membrane and Ponceau-stained. Membranes were blocked in 5% BSA in PBST (PBS containing 0.1% Tween20) for 60 min at RT. The membrane was then incubated in primary antibody (1:1000) as indicated (table S5) for 1 h at RT or at 4°C ON, washed three times for 10 min in PBST, and incubated in fluorescent secondary antibody (1:10000) for 1 h at RT. Lastly, the membranes were again washed three times for 10 min in PBST. The blots were finally imaged via Odyssey M (Licor).

Immunofluorescence (IF)

For IF experiments, siRNA double knockdown was performed as described above. Then, 100.000 dox-induced RPE1 FLAG-ATM cells/well were seeded in 24-well imaging plates. The next day, cells were treated with EdU (10 µM, 30 min), and were washed in PBS and fixed in 4% PFA for 15 min. Then, the cells were permeabilized by adding PBS-Triton X 0.2% incubation for 5 min at RT. Afterward, the cells were blocked with 5% BSA in PBS for 1 h at RT. Then, the plates were incubated in primary antibody (anti-gammaH2AX (1:1000, Millipore, Cat.Nr.: 05-636) and Cyclin A (1:500, Santa Cruz Biotechnology, Cat.No: sc-751) diluted in 5% BSA-PBS, ON at 4°C), washed three times in PBST and finally incubated in secondary Alexa Flour antibody in darkness for 1 h at RT. After 3x PBS washes, the cells were lastly stained with DAPI (1 µg/mL, 5 min) and washed with PBS. EdU was labeled in Click-iT reaction with Alexa-Fluor 647 Azide (Thermo Fisher, A10277), and the plate was kept in mH2O at 4°C. The plates were imaged on the Opera Phenix spinning disc confocal microscope (PerkinElmer, 40x magnification). The data was analyzed in Harmony software (PerkinElmer).

Cryo-EM sample preparation and data collection

Cryo-EM samples for the ATM variants were prepared in 50 mM HEPES pH 7.5, 100 mM NaCl, and 1 mM TCEP with 3 μM ATM, 2.5 mM nucleotide (ATP or ADP), 5 mM MgCl_2_, and 5 mM p53^12^, unless stated otherwise (see figure legends). For the transition-state samples, the cryo-EM buffer was supplemented with 25 mM MgCl_2_, 15 mM NaF, and 3 mM ADP (*25-27*). An FEI Vitrobot Mark III maintained at 4°C and 100% humidity was used to blot and plunge freeze in liquid ethane 4 μL of ATM sample applied on glow discharged grids, with blotting force 5 and blotting time of 3 s.

Data were collected on various transmission electron microscopes (see tables S1, S3-S4). EPU software was used for automatic data collection in counting Super Resolution mode with binning 2. For all the datasets, the physical pixel size range was 0.73-0.955 Å, total exposure time was 1.5 to 4.2 s with a total dose of 34-45 e^-^/Å^2^ at under focus values between -1.0 to -2.6 µm.

Cryo-EM data processing and deposition

Cryo-EM data were processed using the *cryoSPARC* v4.7 package (*45*). Briefly, blob picker was used to pick ATM particles on the Motion Corrected (*46*) and CTF Estimated (CTFFIND4) (*47*) movies. 2D class averages of good particles were used for template-based particle picking. Initial sorting of the good particles from junk through 2D classification, ab-initio reconstruction, and heterogenous refinement was performed on binned (128^2^ pixels box size) particles (*45*). After iterative cycles of extensive cleaning, homogeneous refinement (C1 symmetry) followed by non-uniform refinements with C1 and C2 symmetry was performed on unbinned particles (see figs. S4-S10, S16-S18, S20-S29, and tables S1, S3-S4) (*48*). To improve local resolution, the maps were locally refined using local mask around the C-terminal FATKIN or N-terminal N-solenoid (Spiral and Pincer) domains (*45*). EM density maps were visualized in UCSF Chimera X (*49*). Previously reported ATM homodimer structures were used for initial model fitting into the EM density maps using UCF Chimera X and Coot (*49, 50*), followed by refinement in PHENIX was carried out until the models were ready for deposition (*51*).

Model building of ATM active sites

ATM homodimer structure of PDB 8OXM (*10*) was split into N-solenoid (Spiral and Pincer) and FATKIN (FAT and Kinase domains) halves, and the two halves were rigid body fitted in Chimera X and Coot into the respective locally refined maps, followed by refinement using StarMap (*52*) and PHENIX (*51*). The active-site model building was carried out against the maps resolved with C2 symmetry. The p53^12^ peptide from previously reported ATM structure (PDB 8OXM) (*10*) was remodeled into the EM densities of the ATM structure reported here through rigid body- and atomic model-fitting. Finally, real-space refinement in PHENIX was run using default parameters in addition to adjusting the weight to 0.1 and targeting rotamer outliers using a sigma of 1.0 (*51*).

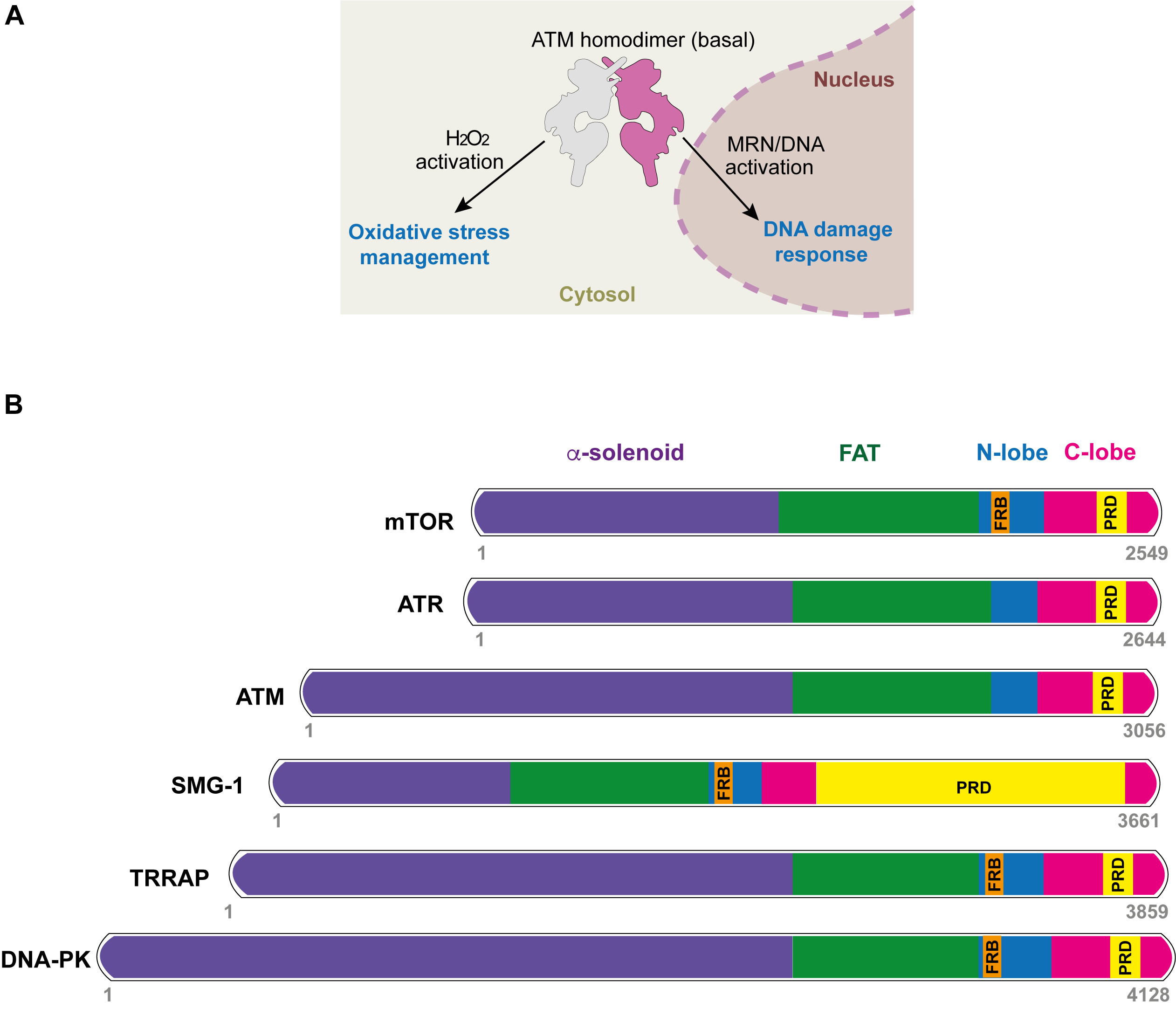

**Fig. S1. ATM belongs to PIKKs family enzyme with two signaling roles: DNA damage and oxidative stress.**

(A) ATM can be activated by two different mechanisms: by MRN/DNA in response to dsDNA breaks and by reactive oxygen species (ROS) in oxidative stress. (B) A schematic representation of the six PIKKs in eukaryotic cells. They share a helical FAT domain and a bilobal C-terminal kinase domain. In addition, there is an insertion in the kinase domain known as the PIKK regulatory domain (PRD).

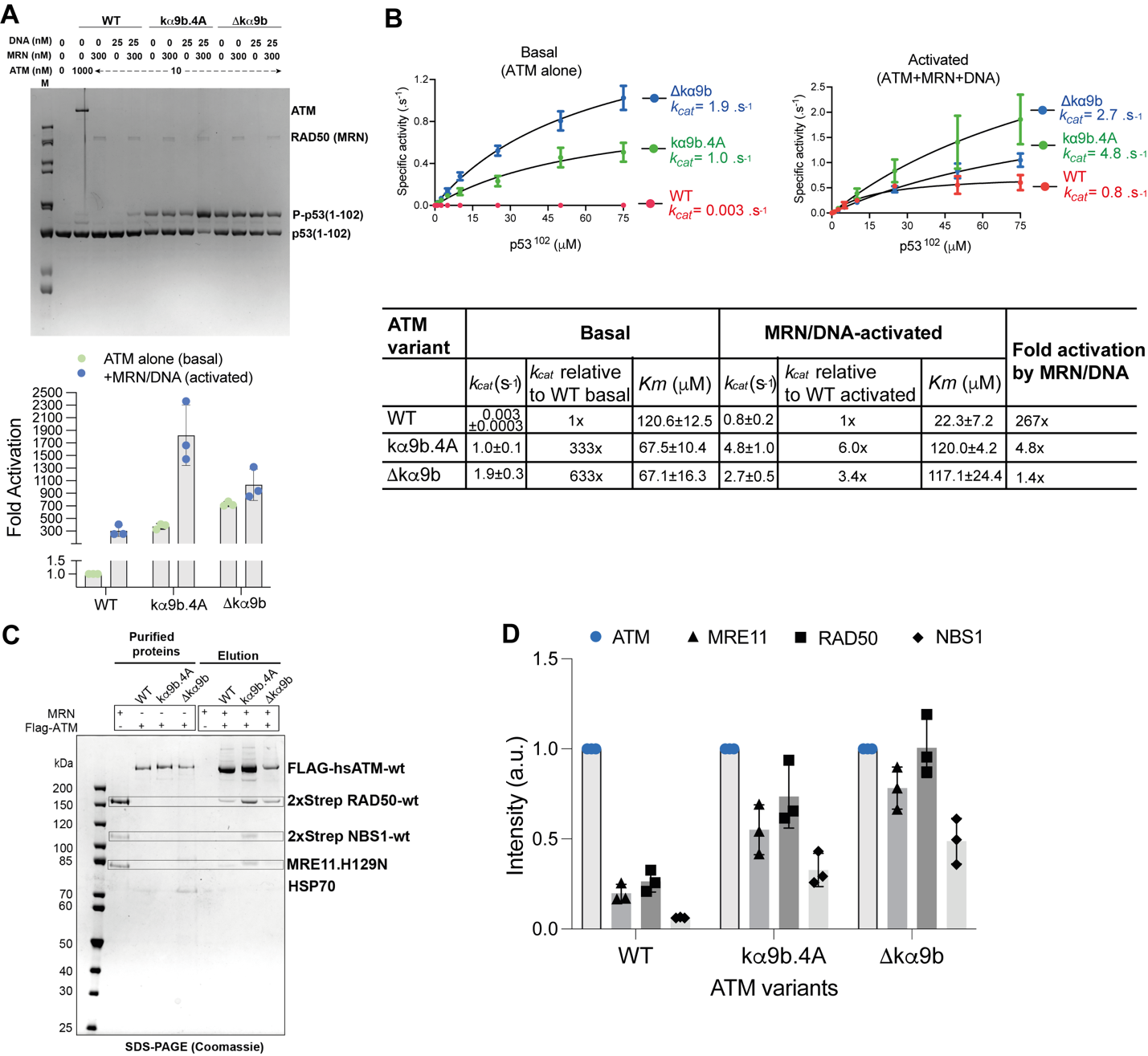

**Fig. S2. In vitro kinase activity of the ATM PRD variants.**

(A) PhosTag gel-based kinase assay of the WT and PRD variants (kα9b.4A and Δkα9b) of ATM, showing kα9b.4A and Δkα9b can constitutively phosphorylate p53^102^ even in the absence of MRN/DNA, whereas WT ATM requires MRN/DNA-mediated stimulation for full activity. Assays were conducted with 10 nM ATM, 25 μM p53^102^, 1 mM ATP, 5 mM MgCl_2_ at 30°C, in the absence or presence of MRN/DNA (300 nM MRN complex and 25 nM 350 bp dsDNA). For measuring the basal activity of WT ATM (ATM alone, no MRN/DNA), a 100-fold more enzyme (1000 nM) was used (lane 3 from left). The lower panel is a quantitation of the three replicates of the PhosTag gel assay as above. (B) Steady-state kinetic analysis (n = 3) of WT, kα9b.4A, and Δkα9b ATM. Graphs show initial specific activity as a function of p53^102^ substrate concentration, for both basal conditions (left) and MRN/DNA-activated (right). The table (lower) summarizes the *k*_cat_, *K*_m_, and fold activation relative to WT ATM. (C) In vitro binding assay of ATM (WT, kα9b.4A, and Δkα9b) to MRN complex in the presence of 200 bp dsDNA, where the MRN complex (untagged-MRE11, 2xStrep-RAD50, and 2xStrep-NBS1) was pulled down on FLAG-tagged-ATM variants using anti-Flag M2 affinity beads; ATM and MRN complex eluted from the anti-Flag affinity beads were resolved in a SDS-PAGE protein gel, stained using Coomassie blue. The pull-down assays were carried out using 2 μM ATM, 5 μM MRN, 2 μM DNA, 1 mM ATP, and 5 mM MgCl_2_. (D) A quantification (n=3) of the band intensities corresponding to ATM, MRE11, RAD50, and NBS1, showing that the PRD mutants, kα9b.4A and Δkα9b, bind to MRN complex more efficiently than the WT enzyme.

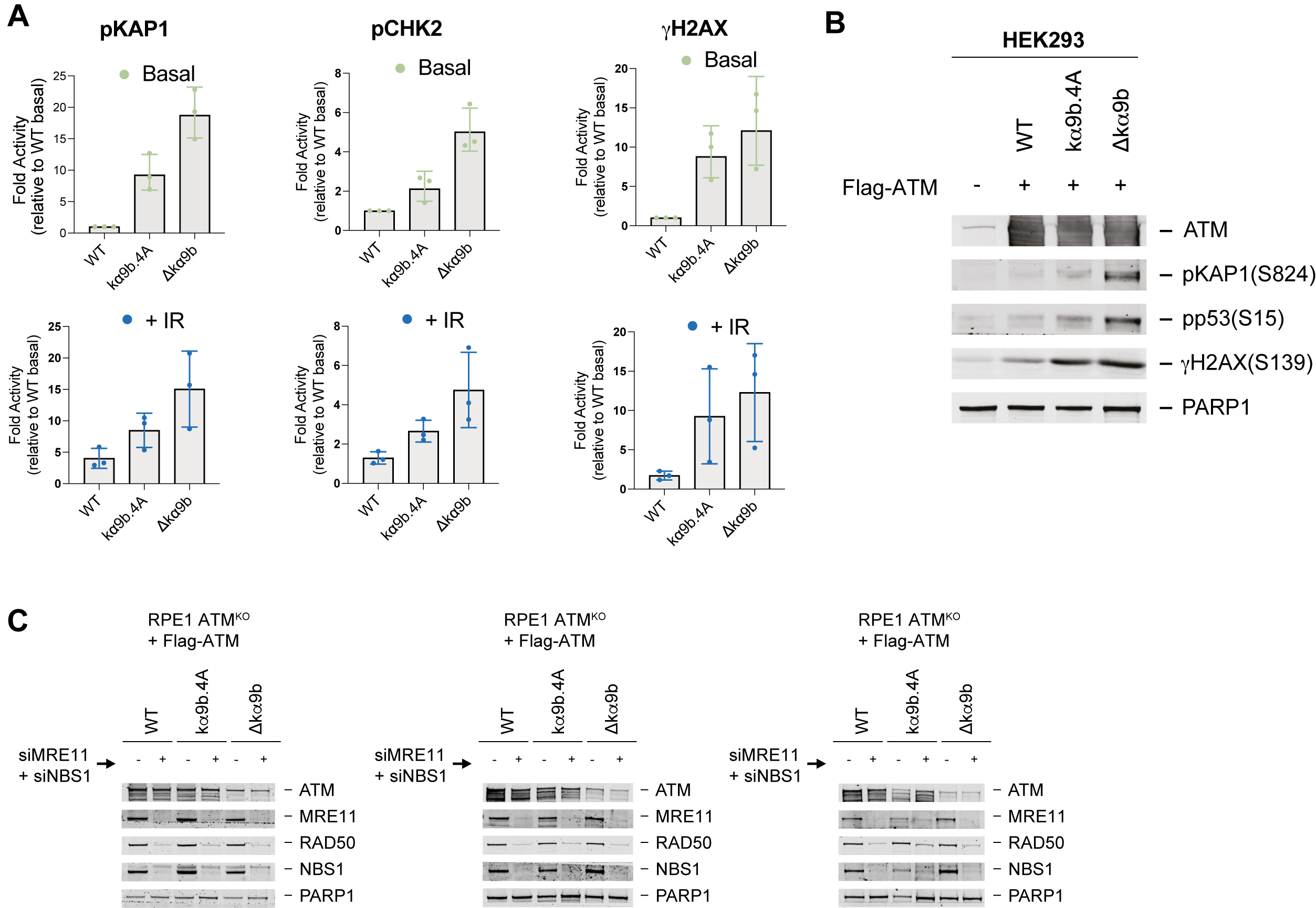

**Fig. S3. Kinase activity of the ATM variants in cells.**

(A) Quantification (from Western blot) (n=3) of the substrate phosphorylation in *ATM TP53* double knockout (KO) RPE1 cells transduced with Flag-tagged ATM variants. The data are normalized to gel loading, kinase expression, and WT activity. Cells were treated with doxycycline prior to the experiment (0.1µg/mL, 24 h). Error bars represent standard deviation (SD). IR=Ionizing radiation of 2 Gy, 1 h recovery. (B) Transient transfection of HEK293 cells with ATM WT, kα9b.4A and Δkα9b, showing phosphorylation of KAP1 on Ser^824^, p53 on Ser^15^, and H2AX on Ser^139^. PARP1 served as a loading control. (C) Western blots for MRE11, RAD50, and NBS1 in *ATM TP53* double knockout (KO) RPE1 cells transduced with Flag-tagged ATM variants, confirming knockdown of MRE11 and NBS1 by siRNAs (siMRE11 and siNBS1). The same *ATM TP53* KO and *MRE11/NBS1* depleted cells were used to measure the γH2AX signals shown in Fig. 1E and 1F (n=3).

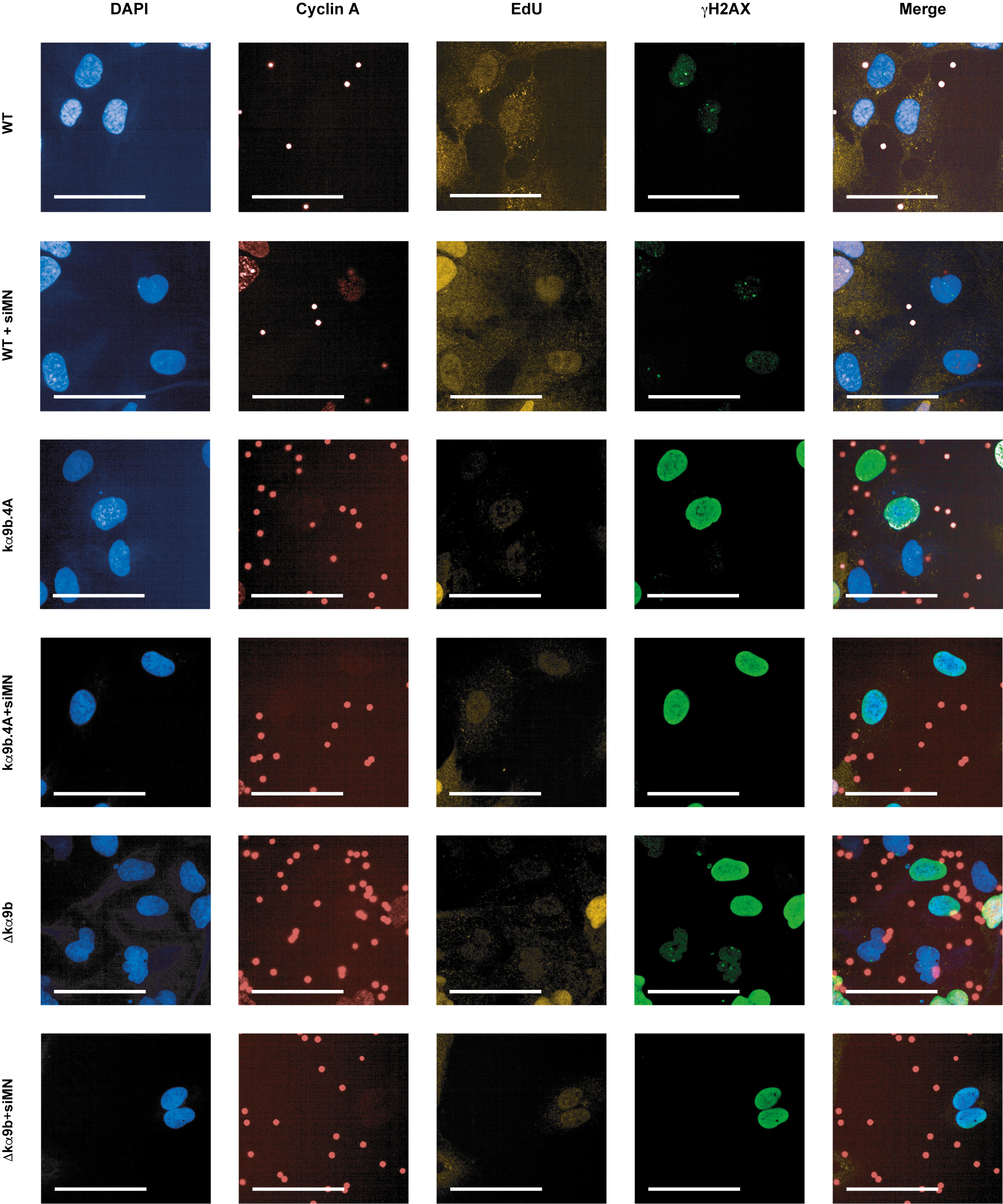

**Fig. S4. Confocal images of the *ATM* and *TP53* double knockout (KO) RPE1 cells.**

Representative images of *ATM TP53* double knockout (KO) RPE1 cells transduced with Flag-tagged ATM variants with and without MRE11/NBS1 depletion (siRNAs). Channels shown: DAPI, Cyclin A, EdU, γH2AX. Scale bar, 50 μm.

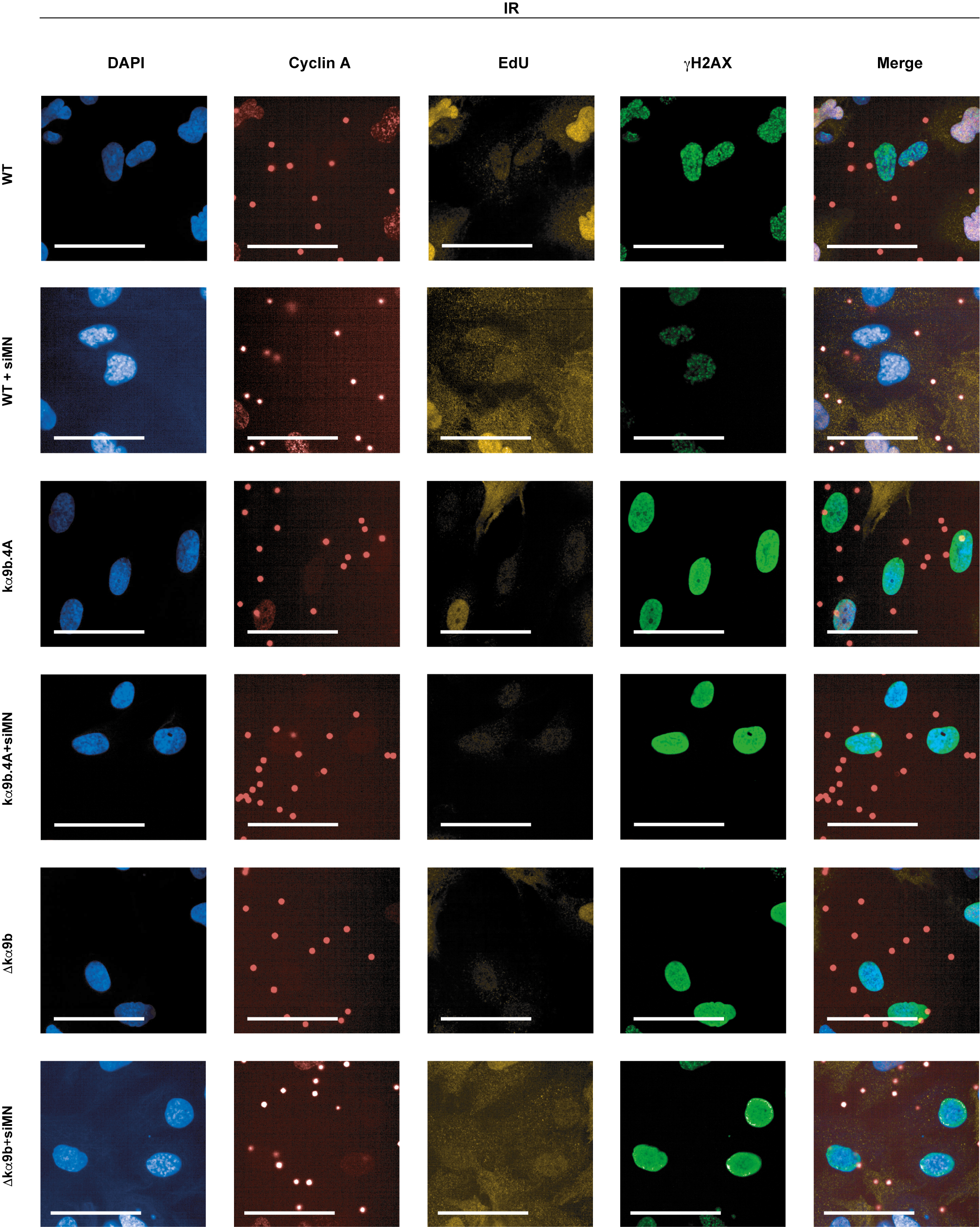

**Fig. S5. Confocal images of the *ATM* and *TP53* double knockout (KO) RPE1 cells after IR exposure.**

Representative images of *ATM TP53* double knockout (KO) RPE1 cells transduced with Flag-tagged ATM variants with and without MRE11/NBS1 depletion (siRNAs). Channels shown: DAPI, Cyclin A, EdU, γH2AX. Scale bar, 50 μm. IR=Ionizing radiation of 2 Gy, 1 h recovery.

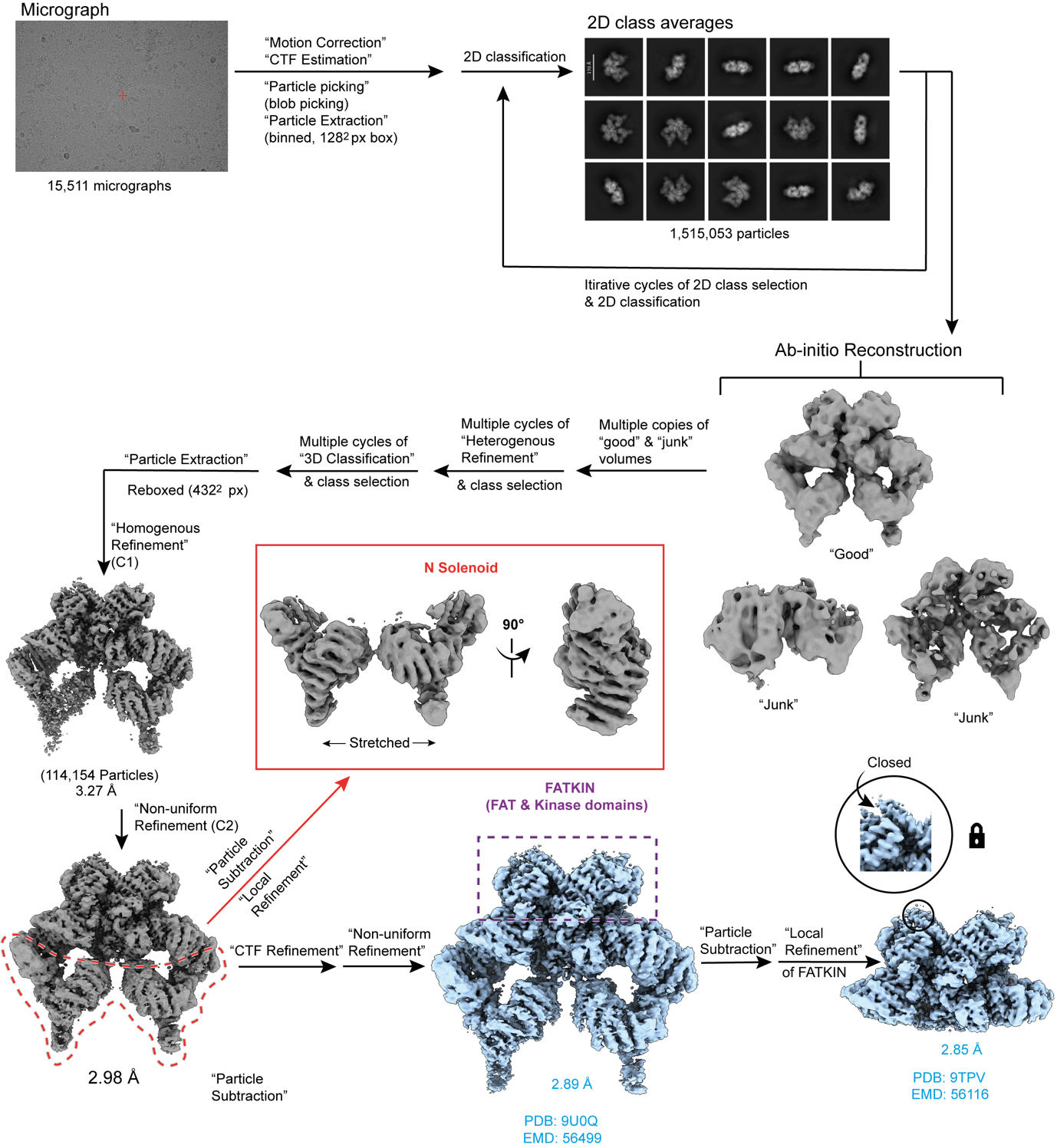

**Fig. S6.** **Cryo-EM single particle analysis data processing workflow of WT ATM in the presence of p53^12^ and TS mimic (ADP/MgF_3_^-^).** Final reconstructions deposited to the EMDB are indicated.

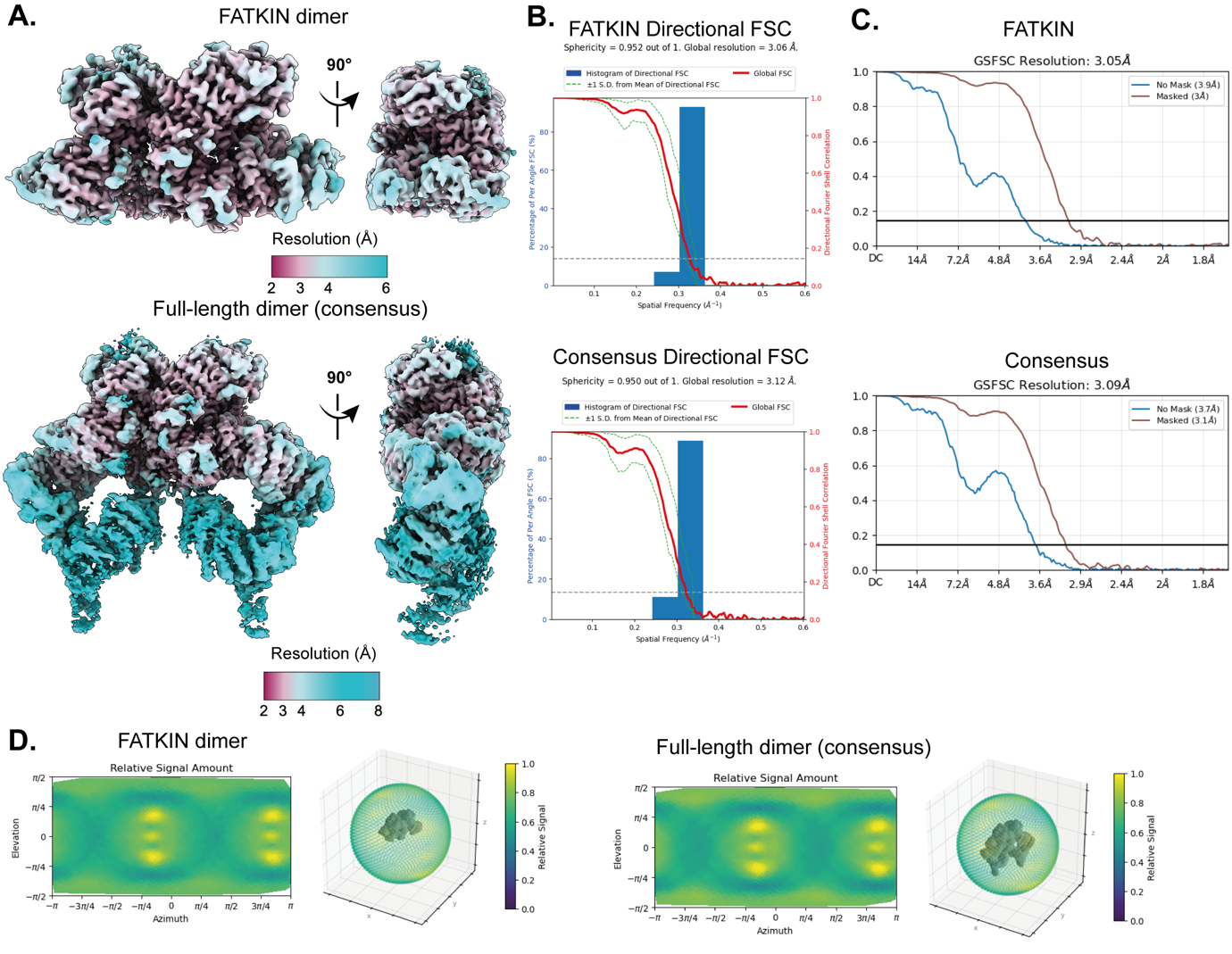

**Fig. S7**. **Cryo-EM map quality metrics for the WT/ADP/MgF_3_^-^/p53^12^ (TS).** (A) FATKIN (local refinement) and consensus maps of the WT ATM bound to ADP/MgF_3_^-^ (TS) colored by local resolution, estimated by *cryoSPARC* v4.7. (B) Directional FSC plots and histograms for the final reconstructions of FATKIN and Consensus maps of WT calculated using 3DFSC (*52*). (C) Gold-standard Fourier Shell Correlation (FSC) plot estimated by *cryoSPARC* v4.7. (D) 2D histograms and 3D view of the directional orientation (Euler angle) of the FATKIN and consensus maps of the WT ATM bound to ADP/MgF_3_^-^ (TS). (E) Consensus and composite maps of WT ATM bound to ADP/MgF_3_^-^ (TS).

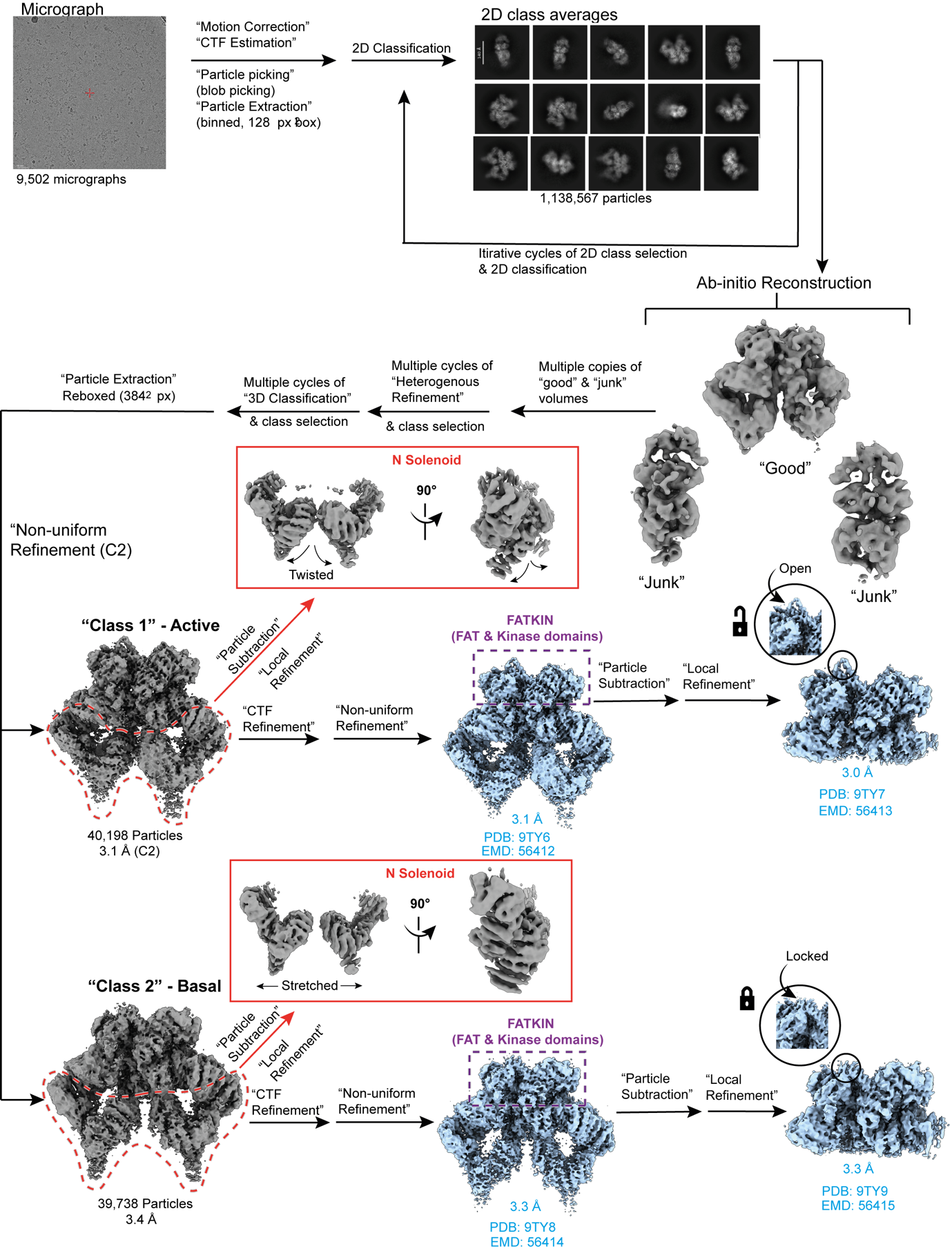

**Fig. S8.** **Cryo-EM single particle analysis data processing workflow of kα9b.4A in the presence of p53^12^ and TS mimic (ADP/MgF_3_^-^).** Final reconstructions deposited to the EMDB are indicated.

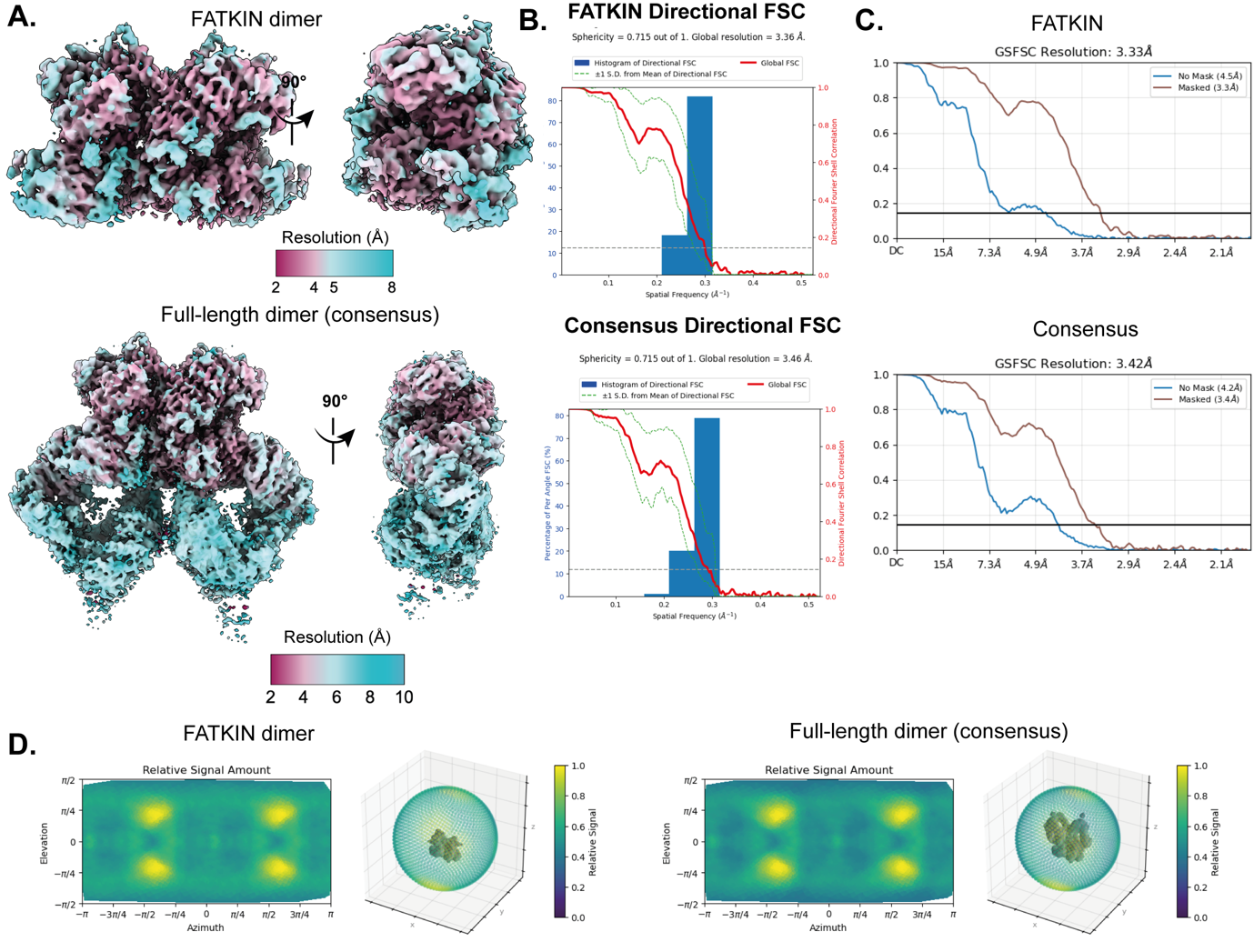

**Fig. S9.** **Cryo-EM map quality metrics for the kα9b.4A/ADP/MgF_3_^-^/p53^12^ (TS) - Class 1.** (A) FATKIN (local refinement) and consensus maps of the kα9b.4A (Class 1) bound to ADP/MgF_3_^-^ (TS) and p53 peptide colored by local resolution, estimated by *cryoSPARC* v4.7. (B) Directional FSC plots and histograms for the final reconstructions of FATKIN and consensus map of kα9b.4A calculated using 3DFSC (*52*). (C) Gold-standard Fourier Shell Correlation (FSC) plot estimated by *cryoSPARC* v4.7. (D) 2D histograms and 3D view of the directional orientation (Euler angle) of the FATKIN and consensus maps of ATM kα9b.4A bound to ADP/MgF_3_^-^ (TS) and p53 peptide. (E) Consensus and composite maps of ATM kα9b.4A bound to ADP/MgF_3_^-^ (TS) and p53 peptide.

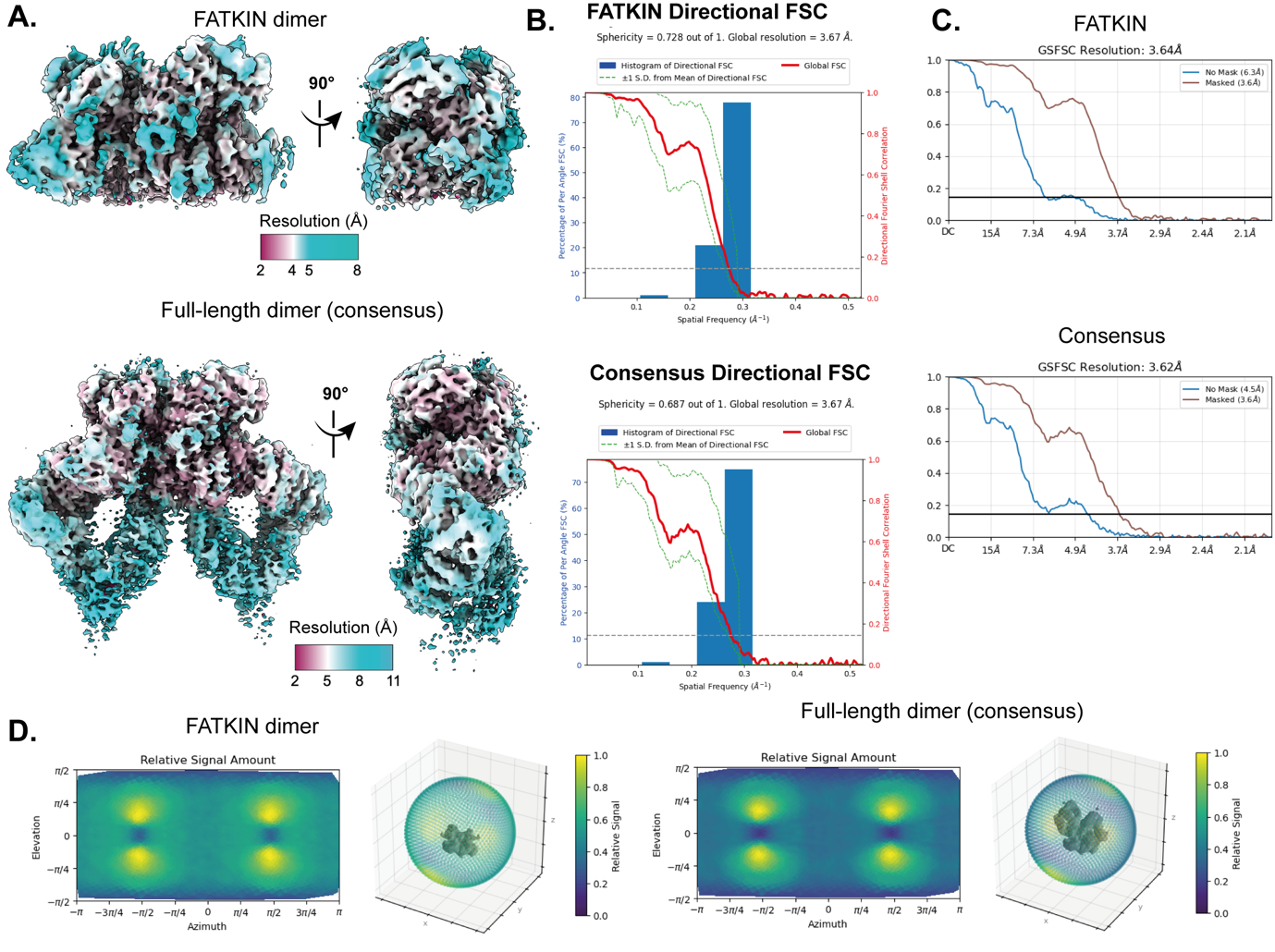

**Fig. S10.** **Cryo-EM map quality metrics for the kα9b.4A/ADP/MgF_3_^-^ (TS) - Class 2.** (A) FATKIN (local refinement) and consensus maps of the kα9b.4A (Class 2) bound to ADP/MgF_3_^-^ (TS) colored by local resolution, estimated by *cryoSPARC* v4.7. (B) Directional FSC plots and histograms for the final reconstructions of FATKIN and consensus map of kα9b.4A calculated using 3DFSC (*52*). (C) Gold-standard Fourier Shell Correlation (FSC) plot estimated by *cryoSPARC* v4.7. (D) 2D histograms and 3D view of the directional orientation (Euler angle) of the FATKIN and consensus maps of ATM kα9b.4A bound to ADP/MgF_3_^-^ (TS). (E) Consensus and composite maps of ATM kα9b.4A bound to ADP/MgF_3_^-^ (TS).

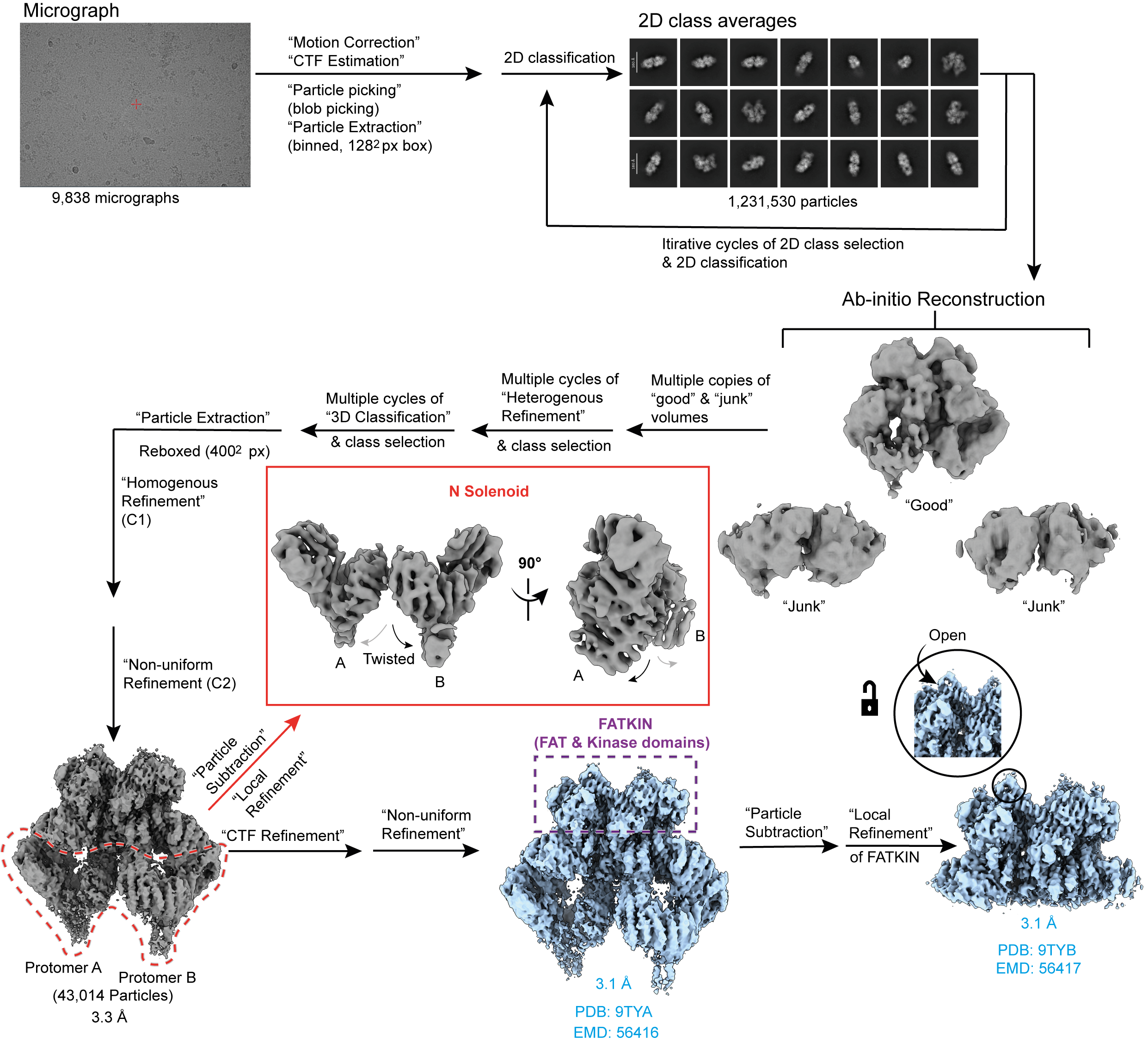

**Fig. S11.** **Cryo-EM single particle analysis data processing workflow of Δkα9b in the presence of p53^12^ and TS mimic (ADP/MgF_3_^-^).** Final reconstructions deposited to the EMDB are indicated.

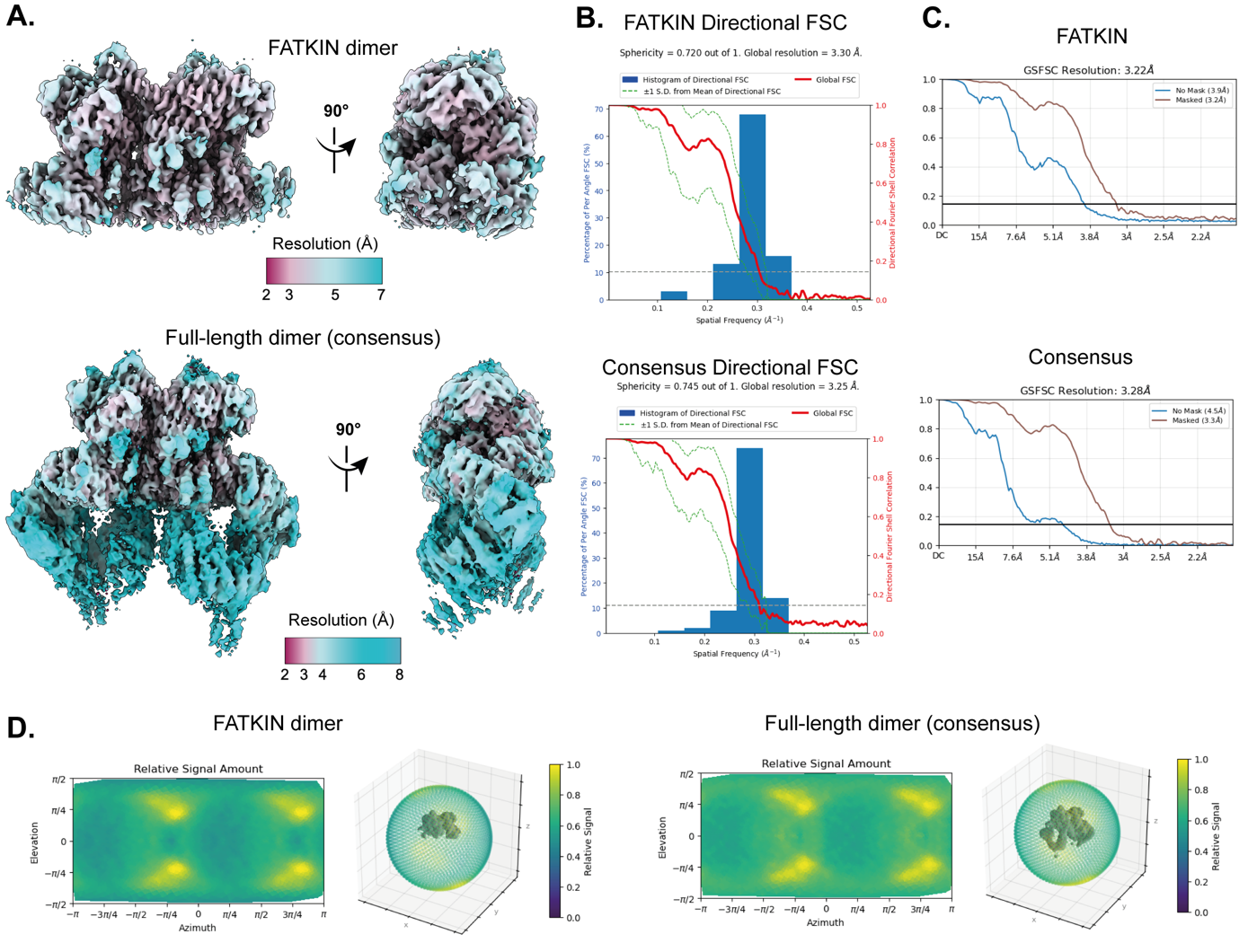

**Fig. S12.** **Cryo-EM map quality metrics for the Δkα9b/ADP/MgF_3_^-^/p53^12^ (TS).** (A) FATKIN (local refinement) and consensus maps of the Δkα9b bound to ADP/MgF_3_^-^ (TS) and p53 peptide colored by local resolution, estimated by *cryoSPARC* v4.7. (B) Directional FSC plots and histograms for the final reconstructions of FATKIN and consensus map of Δkα9b calculated using 3DFSC (*52*). (C) Gold-standard Fourier Shell Correlation (FSC) plot estimated by *cryoSPARC* v4.7. (D) 2D histograms and 3D view of the directional orientation (Euler angle) of the FATKIN and consensus maps of ATM Δkα9b bound to ADP/MgF_3_^-^ (TS) and p53 peptide. (E) Consensus and composite maps of ATM Δkα9b bound to ADP/MgF_3_^-^ (TS) and p53 peptide.

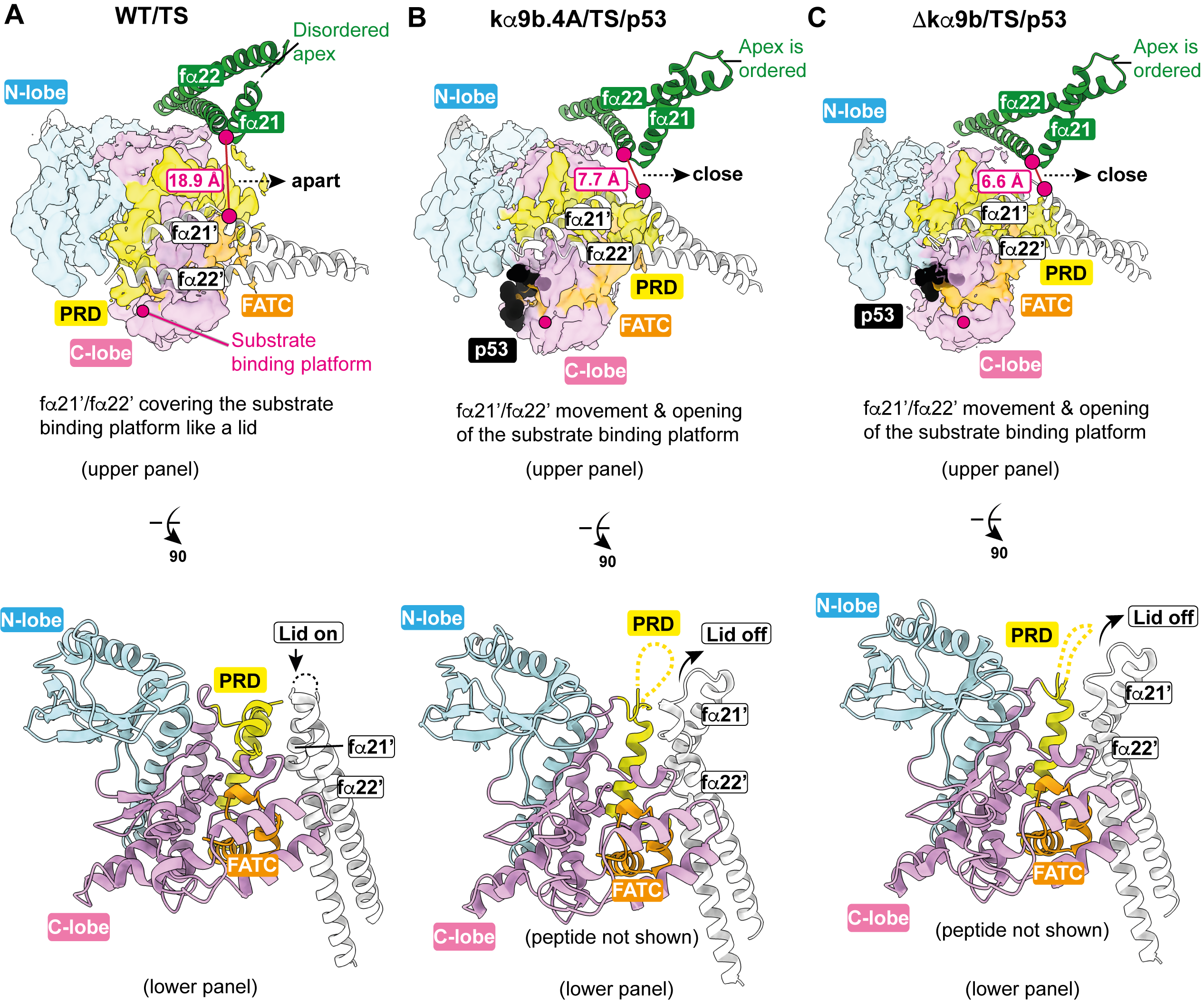

**Fig. S13. Rotation of the two protomers relative to each other in PRD mutant dimers repositions the fα21/fα22 helices across the dimer interface.**

(A, B, C, upper panels) Views from the top of WT ATM (A, upper left panel), kα9b.4A (B, upper middle panel) and Δkα9b (C, upper right panel), with domains colored as in Fig. 1A. The kinase domain is shown as a surface, and two pairs of fα21/fα22 helices are shown as ribbons. The two fα21/fα22 elbows across the dimer interface are distant in the WT ATM conformation (WT basal, left panel), but the rotation of the two protomers in kα9b.4A (middle panel) and Δkα9b (right panel), brings them much closer*.*

(A, B, C, lower panels) View into the active site showing the interaction of the PRD with the kinase and the fα21/fα22 hairpin. In WT ATM, fα21’/fα22’ (from the paired protomer) function as a lid over the substrate-binding platform (Lid on). In the PRD variants, kα9b.4A and Δkα9b, the peptide-binding site is available for peptide binding (Lid off). For clarity, the bound p53^12^ peptide is omitted in the lower panels for the PRD mutants kα9b.4A and Δkα9b.

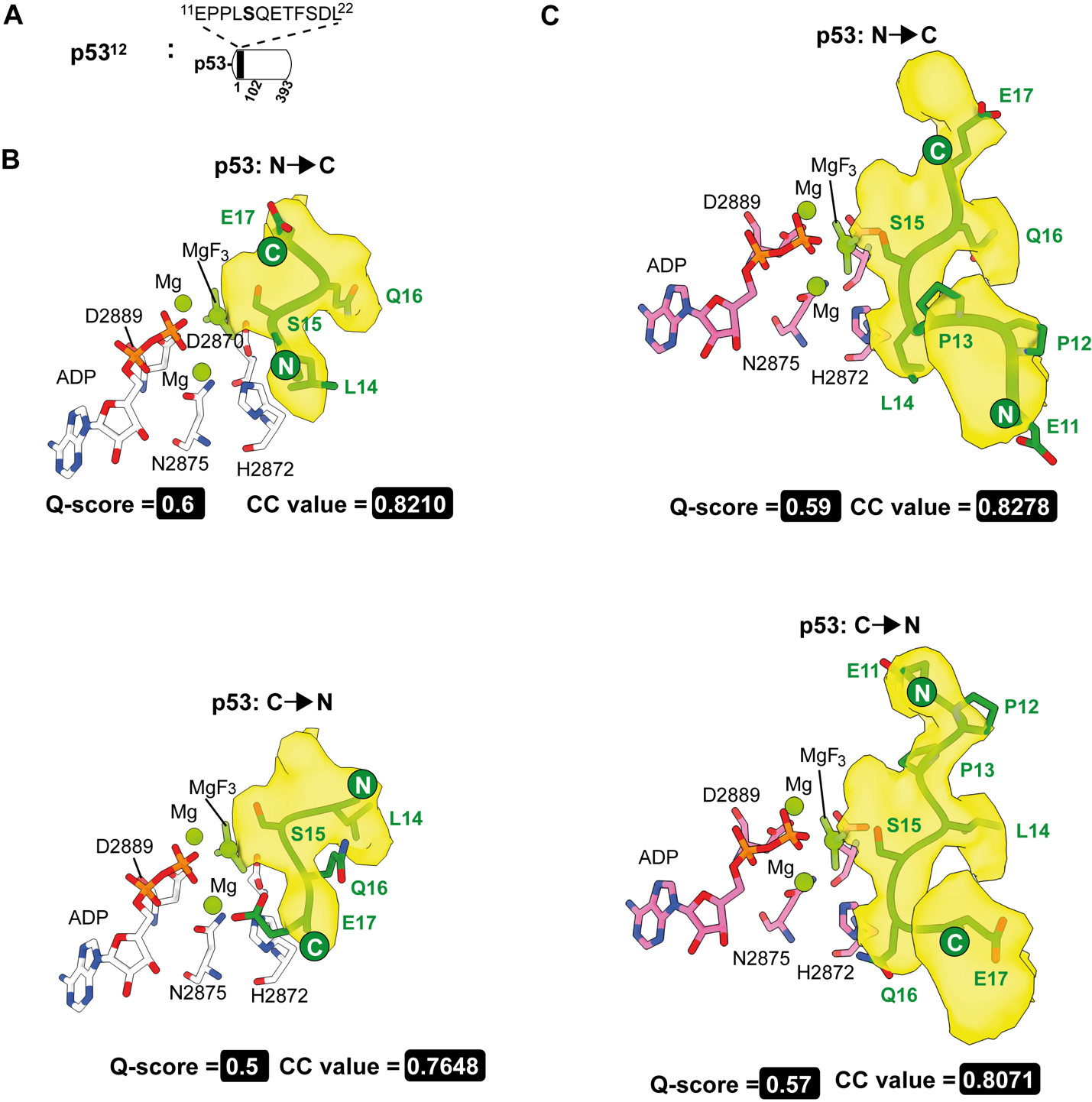

**Fig. S14.** **Cryo-EM density suggests a clear orientation for the bound p53 peptide.**

A p53 peptide model was fit into the EM density in two different orientations for the TS/p53 structures of the two PRD mutants. In both cases the upper panel illustrates the best fit (higher Q-scores) to the density. (A) Sequence of the N-terminal 12 residues of p53 protein (P04637) used as the peptide substrate. (B) The fit of the p53 peptide to the density for the Δkα9b mutant. The Q-score shown is the mean for the p53 peptide as reported by the Q-score plugin for UCF Chimera X (*53*). The correlation coefficient (CC) is as reported by UCF Chimera X fitmap (*54*). The upper panel shows the best fit for the peptide with the N-terminus at the bottom of the figure. The peptide was fit in the reverse direction in the lower panel. (C) The fit of the p53 peptide to the density for the kα9b.4A mutant. The quantitative fit was determined as in (B).

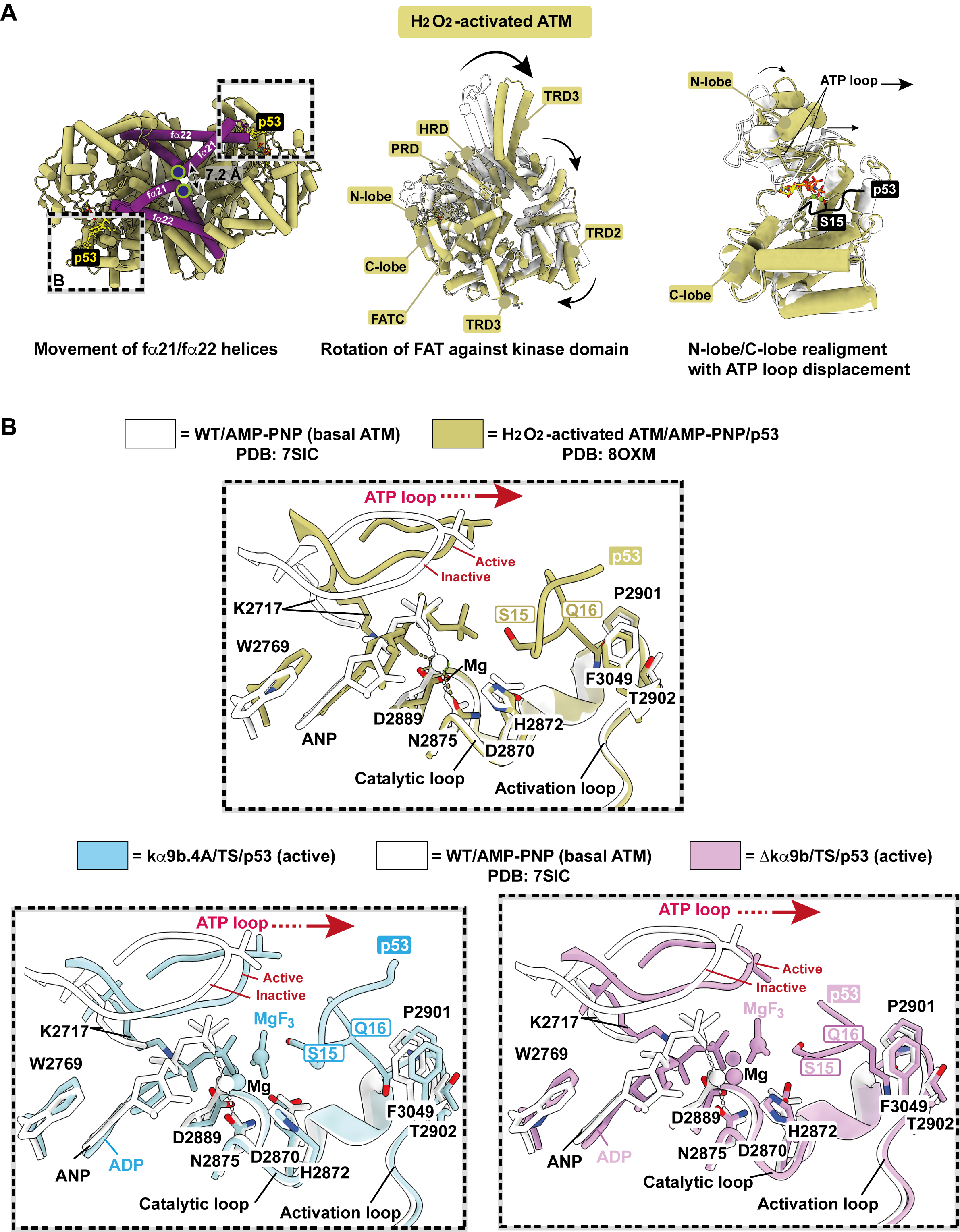

**Fig. S15. TS structures of the PRD mutants** **show activated conformations that are the same as the ATM.Q2971 H_2_O_2_-activated conformation.**

(A) The global conformational change associated with WT H_2_O_2_-mediated activation (olive, 8OXM) compared with the basal WT (white, 7SIC). (B) Upper panel, conformational changes in the active site for WT basal (white) overlaid with ATM.Q2971A H_2_O_2_-activated (olive). Lower panels, conformational changes associated with the kα9b.4A and Δα9b mutants bound to TS/p53 shown overlaid with WT basal (white).

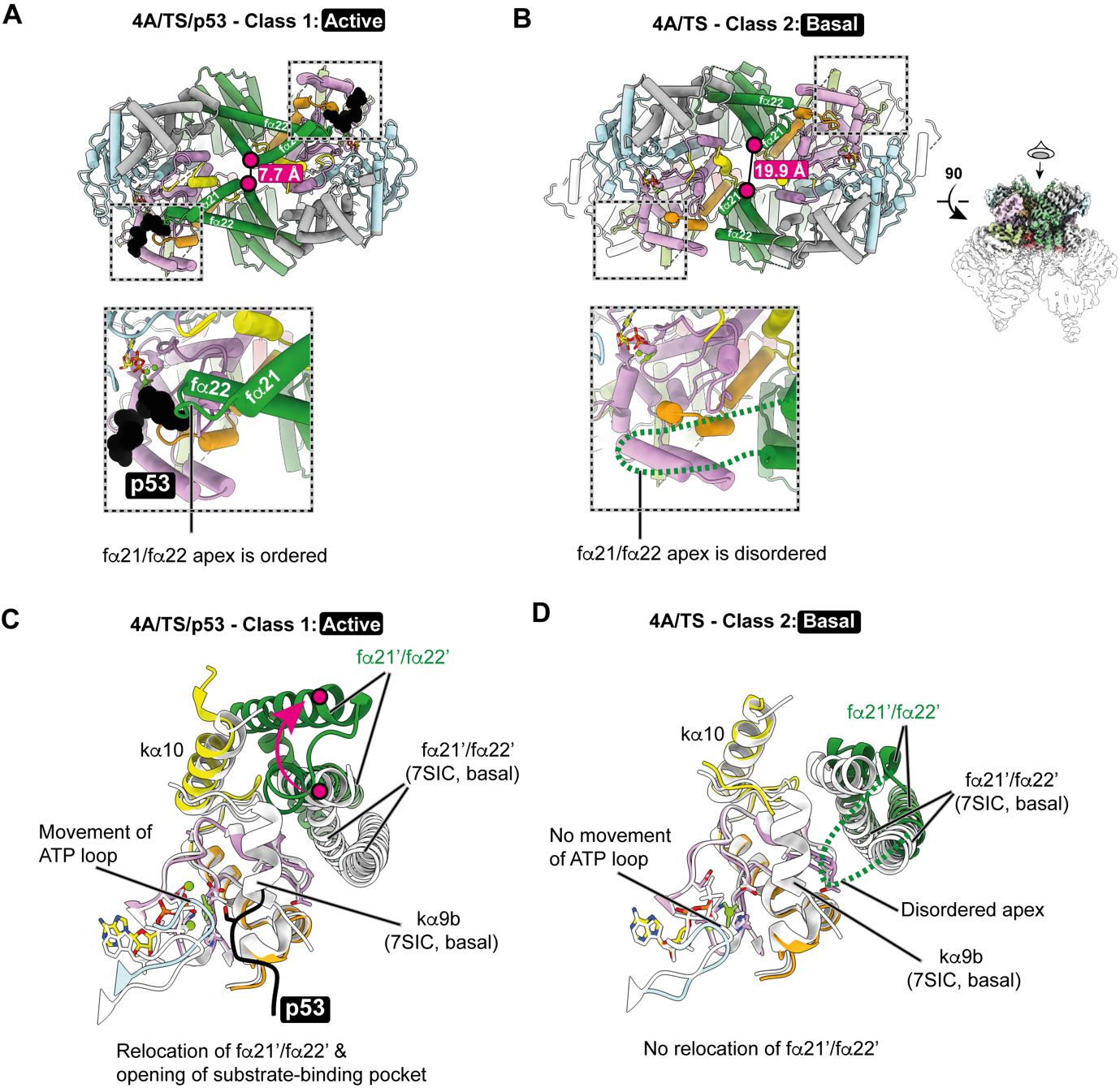

**Fig. S16. Two conformations of kα9b.4A TS structure in the presence and absence of p53 peptide substrate.**

(A) When the TS and p53^12^ peptide are bound (class 1), the shift of the TRD3 relative to the kinase domain produces global conformation changes that move the fα21/fα22 hairpins in the dimer toward each other upon activation (upper panel). Concurrent with this movement, there is a change in order, so that the apex of the fα21/fα22 hairpin is fully ordered (lower panel). (B) When only the TS component (ADP/MgF_3_^-^) is bound, but the peptide is not present (class 2), the fα21/fα22 hairpins in the dimer are distant, characteristic of the basal-state conformation (upper panel). In this state, the apex of the fα21/ fα22 is disordered (lower panel). (C) When the TS and p53^12^ are present, the fα21/ fα22 hairpin, which is part of the TRD3 region of the FAT domain, becomes repositioned relative to the kinase domain. This repositioning is characteristic of activation, which brings the hairpin into contact with kα10 in the C-lobe of the kinase domain and causes the N-lobe of the kinase domain to pivot, optimizing interaction with the nucleotide, relative to the WT ATM in its basal state (7SIC, superimposed in white, superimposed on the kinase C-lobe of the kinase domain). (D) In the absence of p53^12^, the TS/kα9b.4A adopts a conformation that closely agrees with the WT basal ATM (7SIC, in white, superimposed on the C-lobe of the kinase domain).

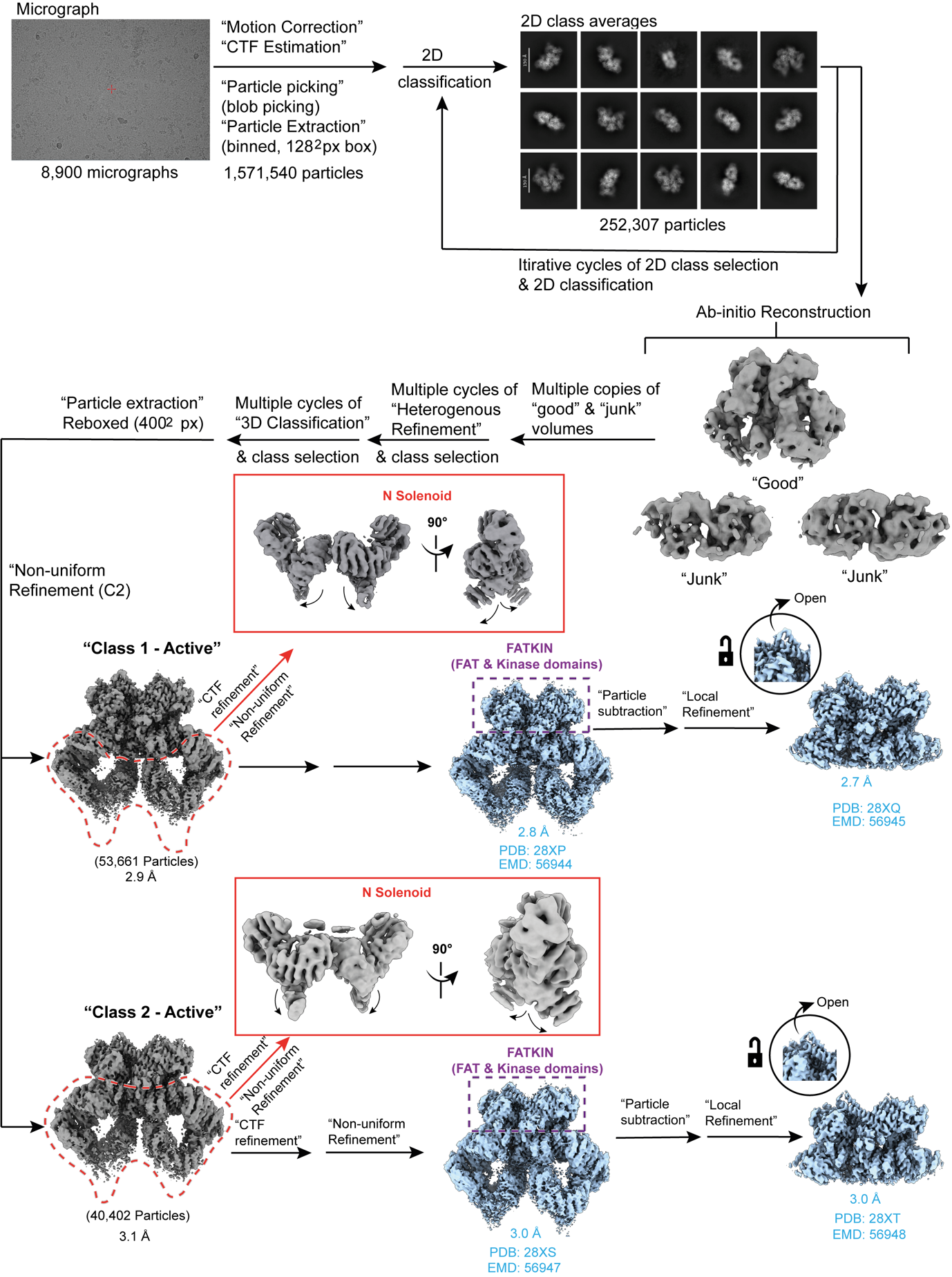

**Fig. S17.** **Cryo-EM single particle analysis data processing workflow of Δkα9b/ATP/p53^12^.** Final reconstructions deposited to the EMDB are indicated.

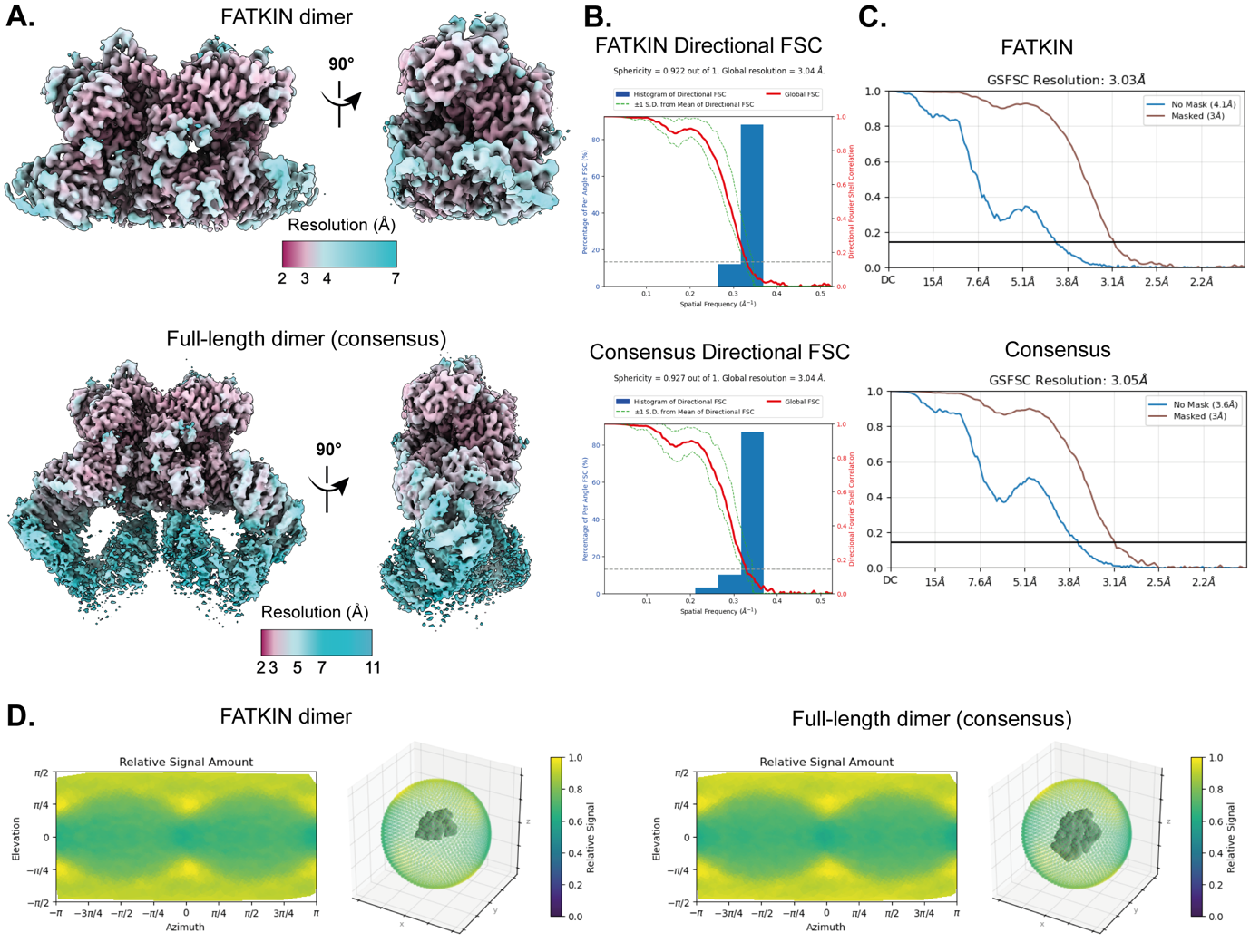

**Fig. S18**. **Cryo-EM map quality metrics for the Δkα9b/ATP/p53^12^ – p53 Conformation 1.** (A) FATKIN (local refinement) and consensus maps of the Δkα9b bound to ATP and p53 peptide colored by local resolution, estimated by *cryoSPARC* v4.7. (B) Directional FSC plots and histograms for the final reconstructions of FATKIN and consensus map of Δkα9b calculated using 3DFSC (*52*). (C) Gold-standard Fourier Shell Correlation (FSC) plot estimated by *cryoSPARC* v4.7. (D) 2D histograms and 3D view of the directional orientation (Euler angle) of the FATKIN and consensus maps of ATM Δkα9b bound to ATP and p53 peptide. (E) Consensus and composite maps of ATM Δkα9b bound to ATP and p53 peptide.

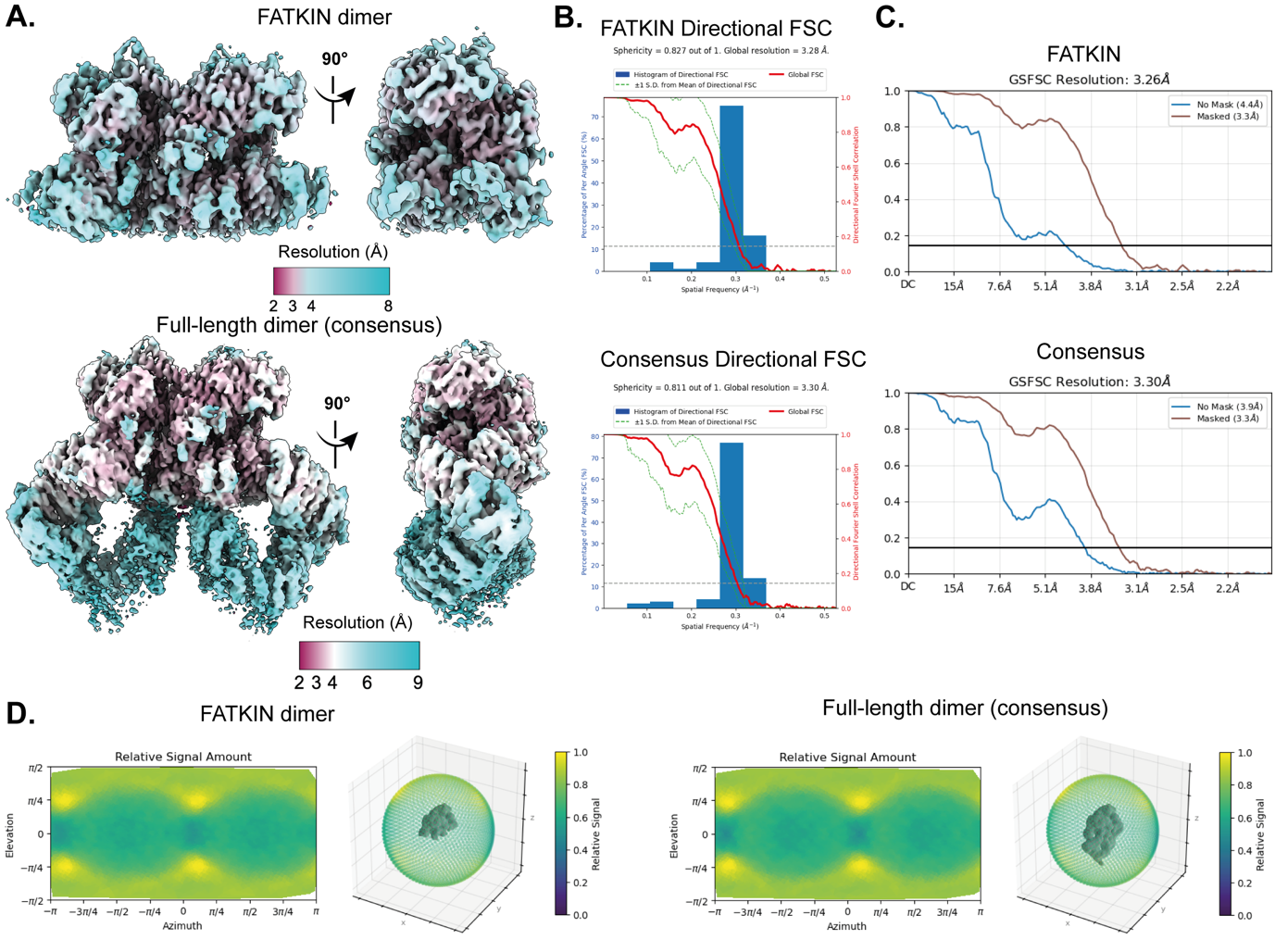

**Fig. S19.** **Cryo-EM map quality metrics for the Δkα9b/ADP*/p53^12^ – p53 Conformation 2.** (A) FATKIN (local refinement) and consensus maps of the Δkα9b bound to p53 peptide and ADP* (symbol “*” indicates ADP formed by in situ hydrolysis of ATP added to the sample), colored by local resolution, estimated by *cryoSPARC* v4.7. (B) Directional FSC plots and histograms for the final reconstructions of FATKIN and consensus map of Δkα9b calculated using 3DFSC(*52*). (C) Gold-standard Fourier Shell Correlation (FSC) plot estimated by *cryoSPARC* v4.7. (D) 2D histograms and 3D view of the directional orientation (Euler angle) of the FATKIN and consensus maps of ATM Δkα9b bound to ADP* and p53 peptide. (E) Consensus and composite maps of ATM Δkα9b bound to ADP* and p53 peptide.

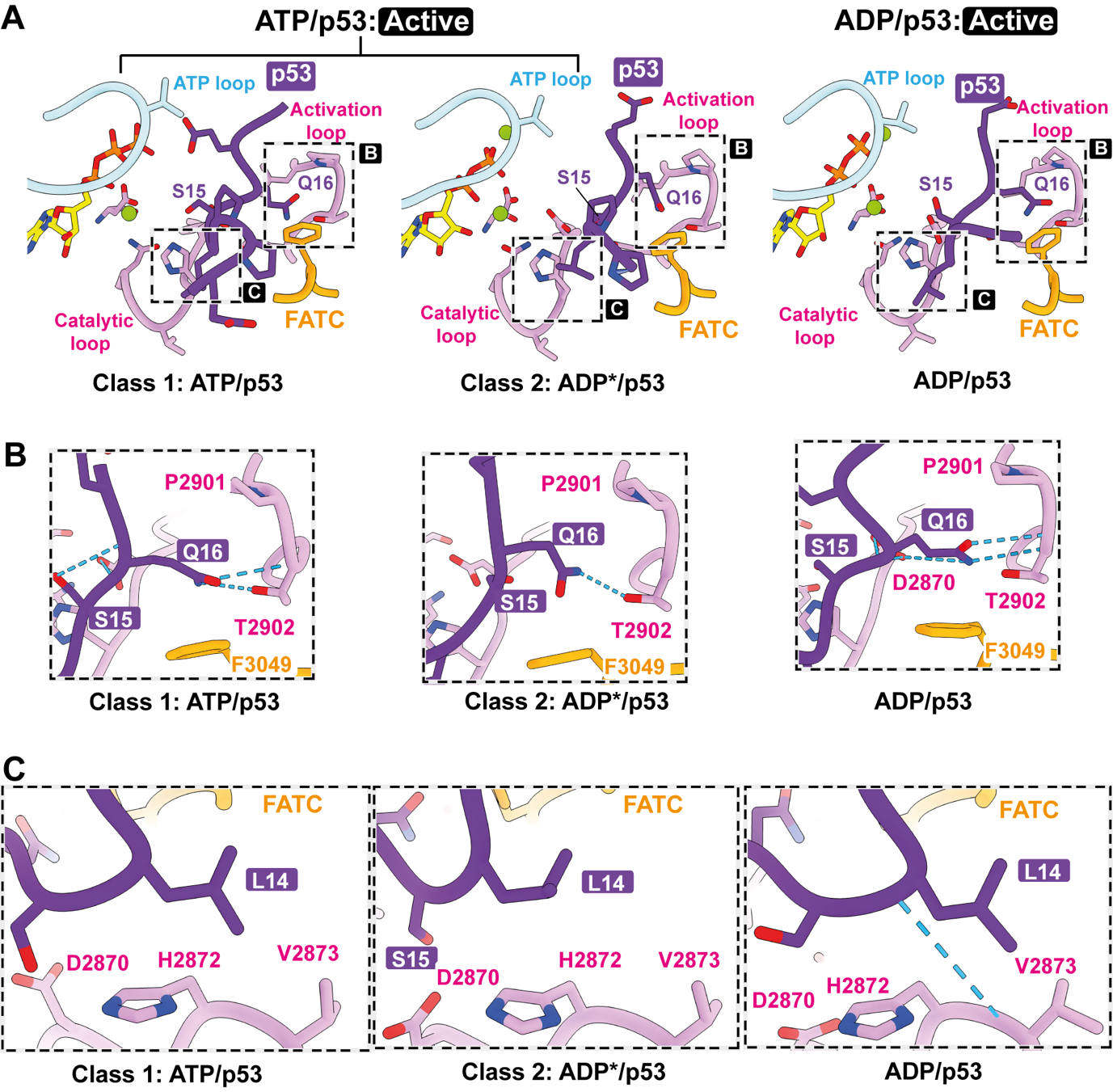

**Fig. S20. p53^12^ peptide interactions in the active site of Δkα9b.**

(A) Models of Δkα9b bound to ATP/p53^12^ (class 1), ADP*/p53^12^ (ADP* denotes ADP formed in situ from ATP) (class 2), and to ADP/p53^12^. The p53^12^ substrate peptide is colored purple, the kinase N-lobe is light blue, C-lobe pink and FATC is orange. (B, C) Close-up views of p53^12^ substrate peptide interactions in the active site.

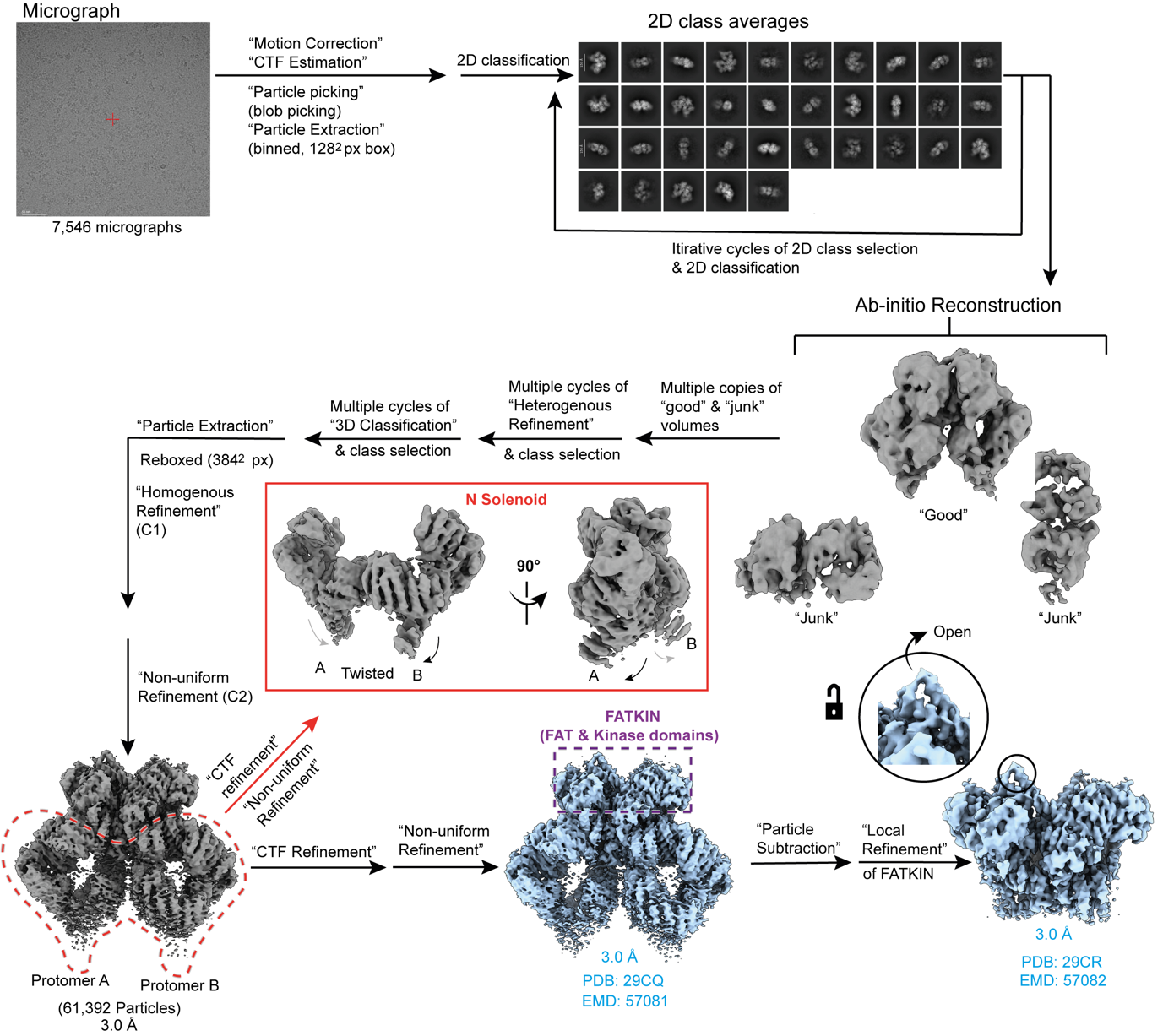

**Fig. S21.** **Cryo-EM single particle analysis data processing workflow of Δkα9b/ADP/p53^12^.** Final reconstructions deposited to the EMDB are indicated.

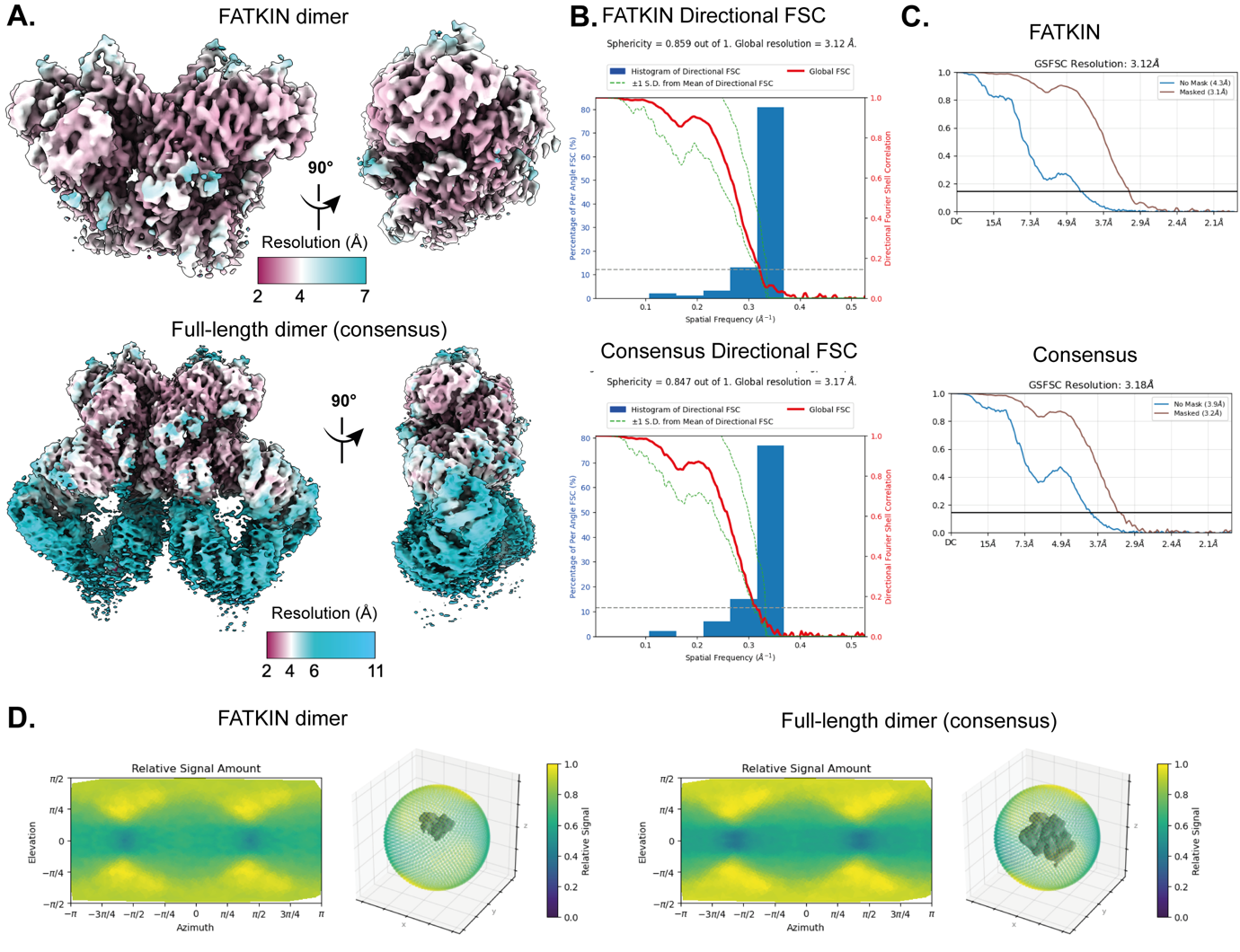

**Fig. S22**. **Cryo-EM map quality metrics for the Δkα9b/ADP/p53^12^.** (A) FATKIN (local refinement) and consensus maps of the Δkα9b bound to ADP and p53 peptide colored by local resolution, estimated by *cryoSPARC* v4.7. (B) Directional FSC plots and histograms for the final reconstructions of FATKIN and consensus map of Δkα9b calculated using 3DFSC (*52*). (C) Gold-standard Fourier Shell Correlation (FSC) plot estimated by *cryoSPARC* v4.7. (D) 2D histograms and 3D view of the directional orientation (Euler angle) of the FATKIN and consensus maps of ATM Δkα9b bound to ADP and p53 peptide. (E) Consensus and composite maps of ATM Δkα9b bound to ADP and p53 peptide.

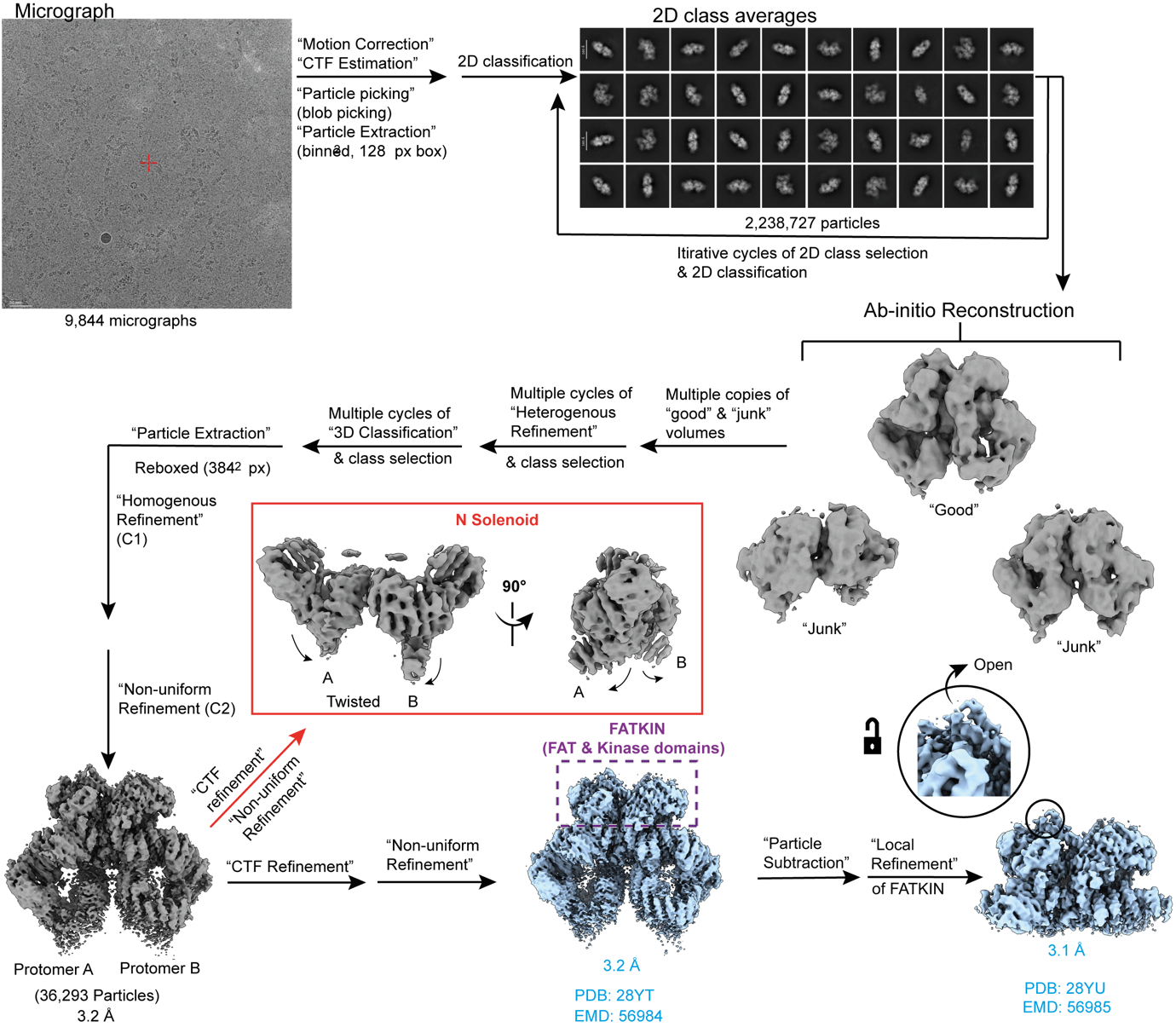

**Fig. S23.** **Cryo-EM single particle analysis data processing workflow of Δkα9b/ATP.** Final reconstructions deposited to the PDB and EMDB are indicated.

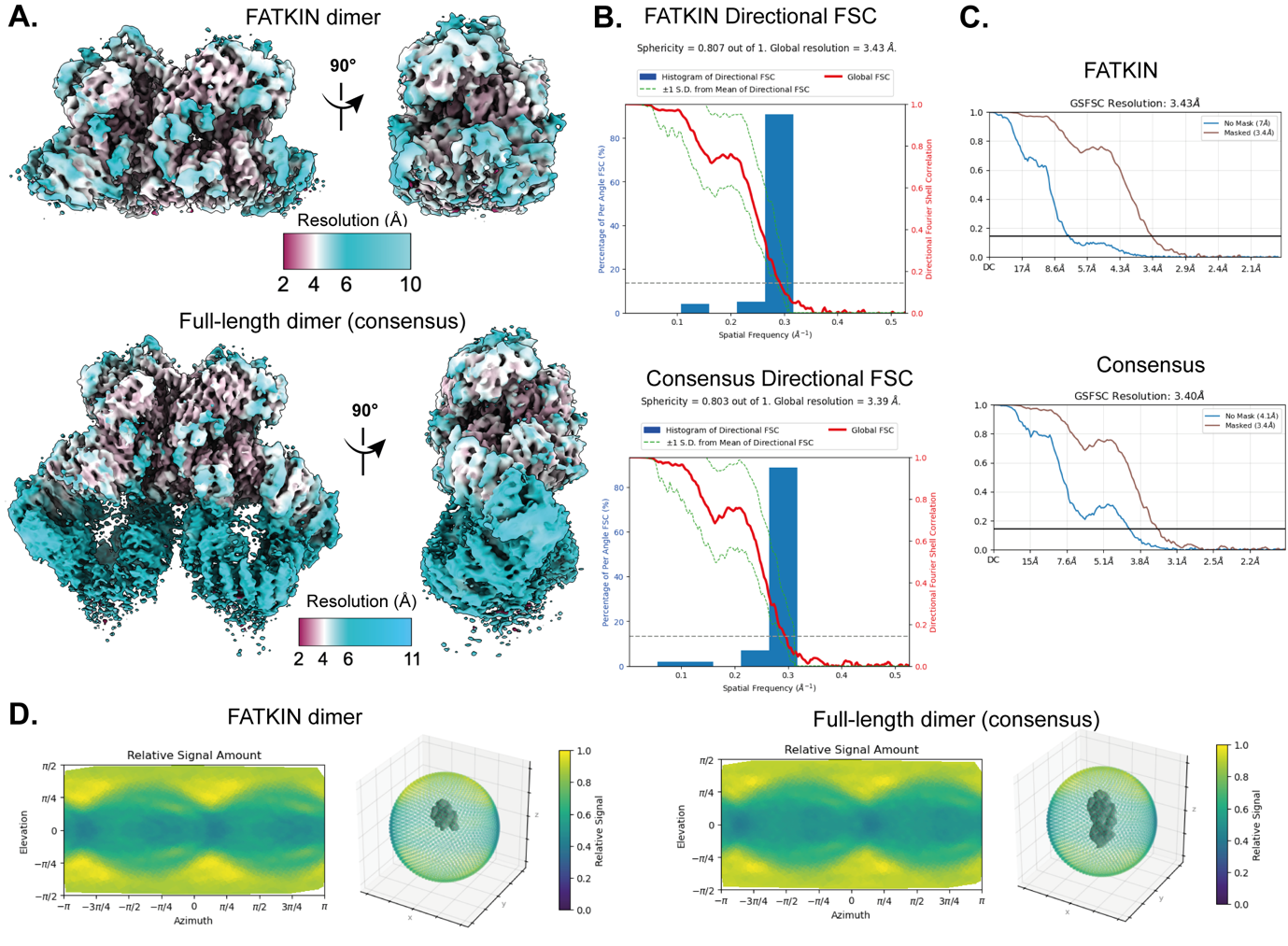

**Fig. S24.** **Cryo-EM map quality metrics for the Δkα9b/ATP.** (A) FATKIN (local refinement) and consensus maps of the Δkα9b bound to ATP colored by local resolution, estimated by *cryoSPARC* v4.7. (B) Directional FSC plots and histograms for the final reconstructions of FATKIN and consensus map of Δkα9b calculated using 3DFSC (*52*). (C) Gold-standard Fourier Shell Correlation (FSC) plot estimated by *cryoSPARC* v4.7. (D) 2D histograms and 3D view of the directional orientation (Euler angle) of the FATKIN and consensus maps of ATM Δkα9b bound to ATP. (E) Consensus and composite maps of ATM Δkα9b bound to ATP.

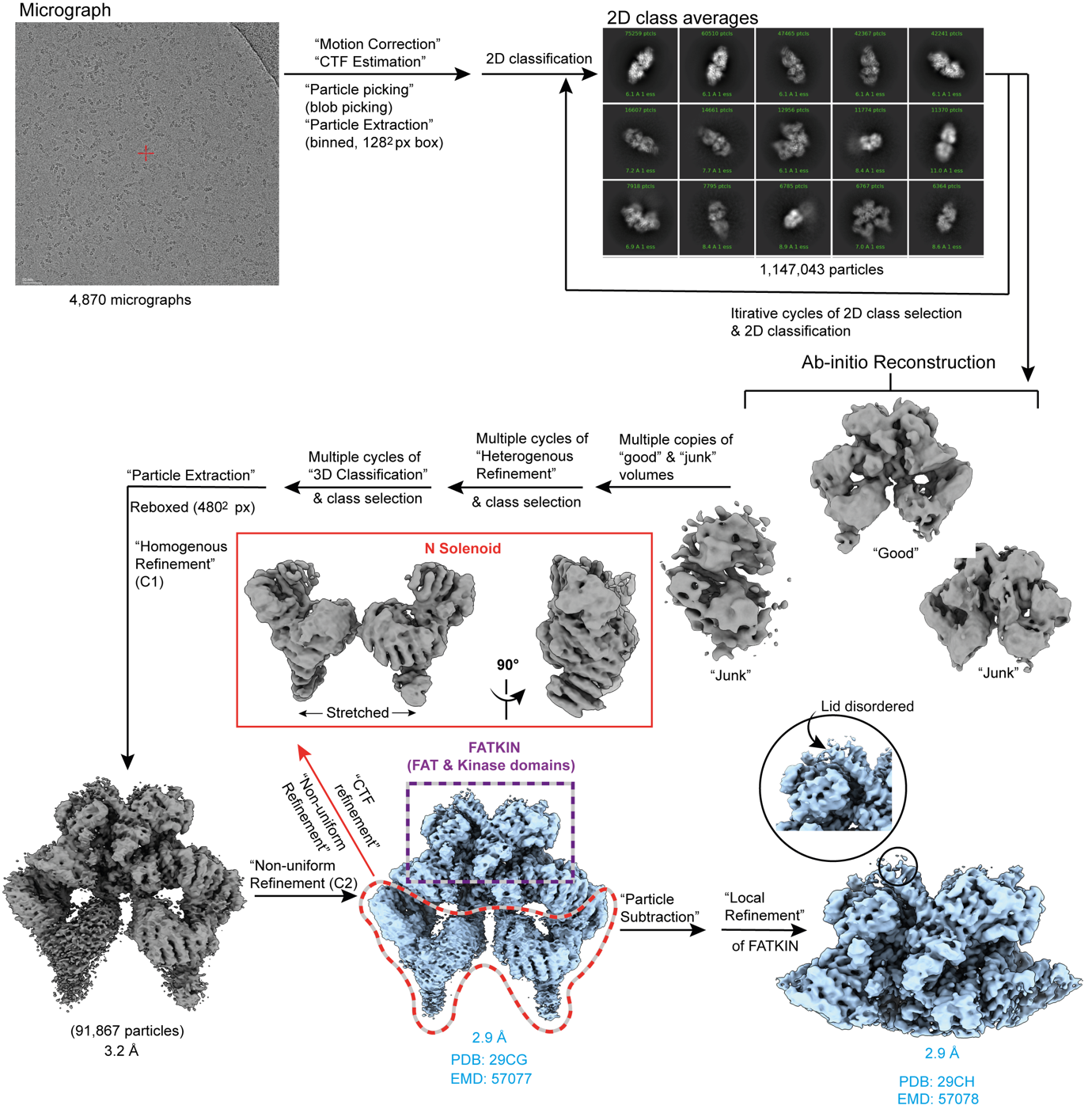

**Fig. S25.** **Cryo-EM single particle analysis data processing workflow of kα9b.4A/ATP.** Final reconstructions deposited to the PDB and EMDB are indicated.

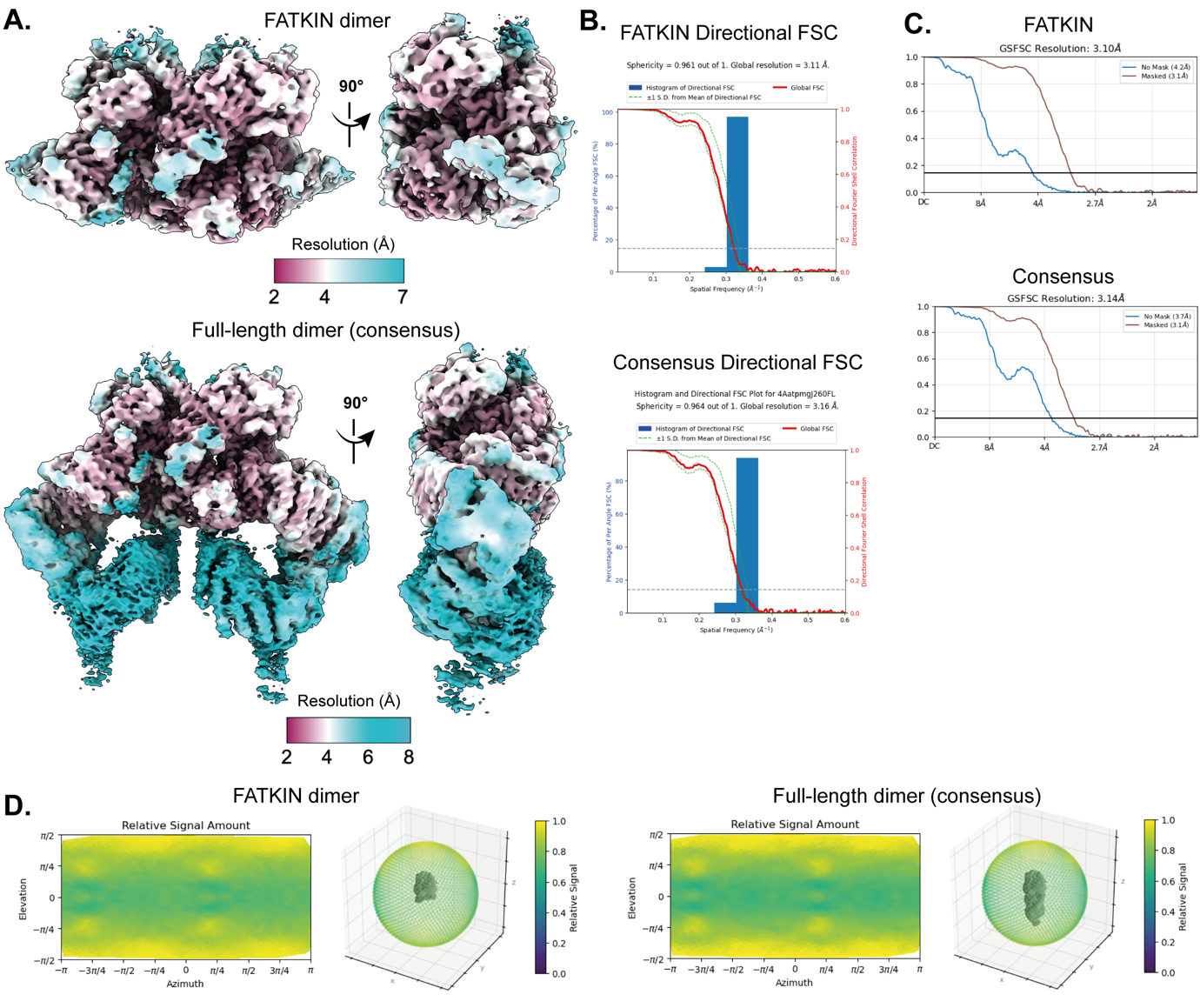

**Fig. S26.** **Cryo-EM map quality metrics for the kα9b.4A/ATP.** (A) FATKIN (local refinement) and consensus maps of the kα9b.4A bound to ATP colored by local resolution, estimated by *cryoSPARC* v4.7. (B) Directional FSC plots and histograms for the final reconstructions of FATKIN and consensus map of kα9b.4A calculated using 3DFSC (*52*). (C) Gold-standard Fourier Shell Correlation (FSC) plot estimated by *cryoSPARC* v4.7. (D) 2D histograms and 3D view of the directional orientation (Euler angle) of the FATKIN and consensus maps of the kα9b.4A bound to ATP. (E) Consensus and composite maps of kα9b.4A bound to ATP.

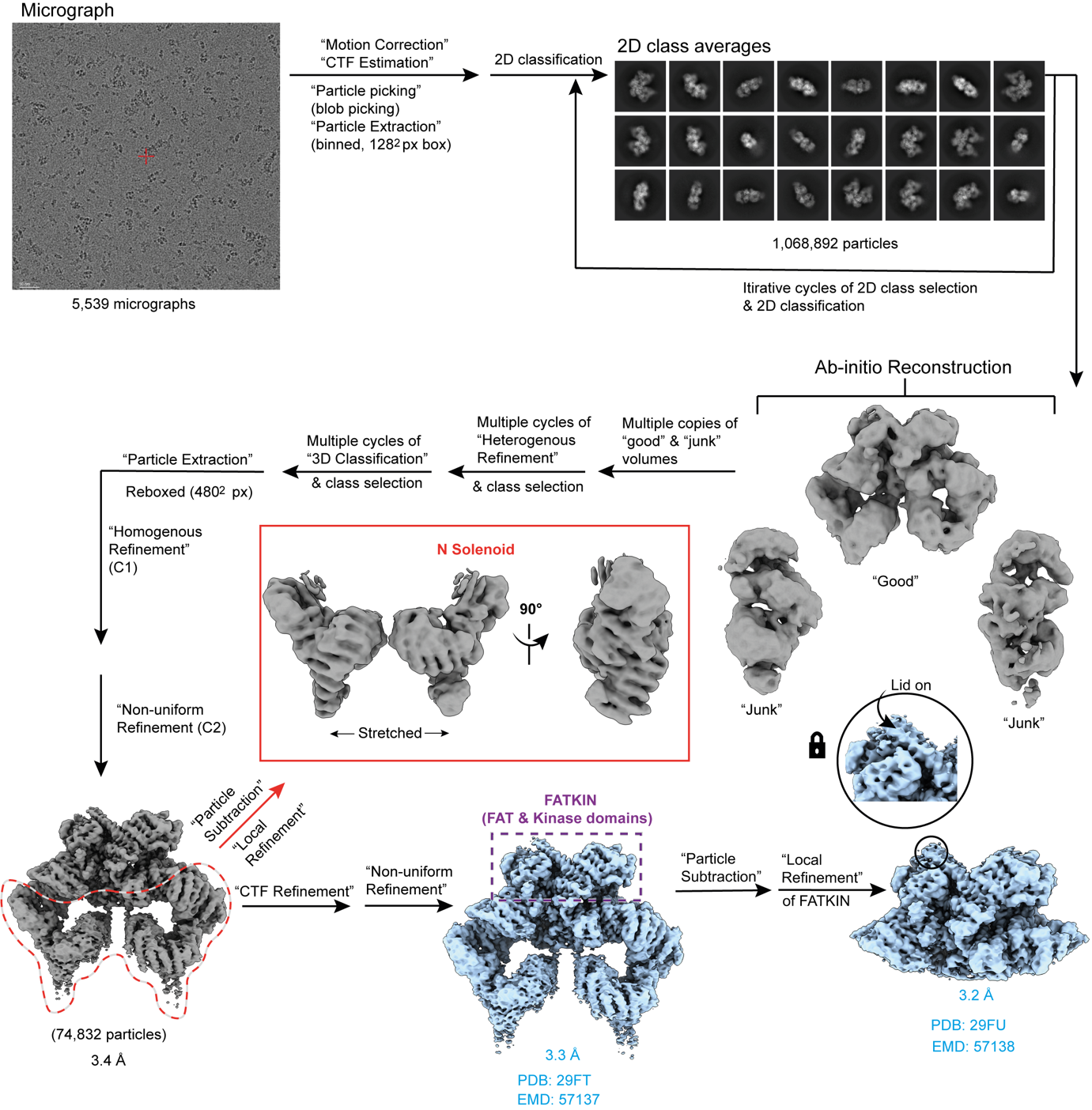

**Fig. S27.** **Cryo-EM single particle analysis data processing workflow of WT/ATP.** Final reconstructions deposited to the PDB and EMDB are indicated.

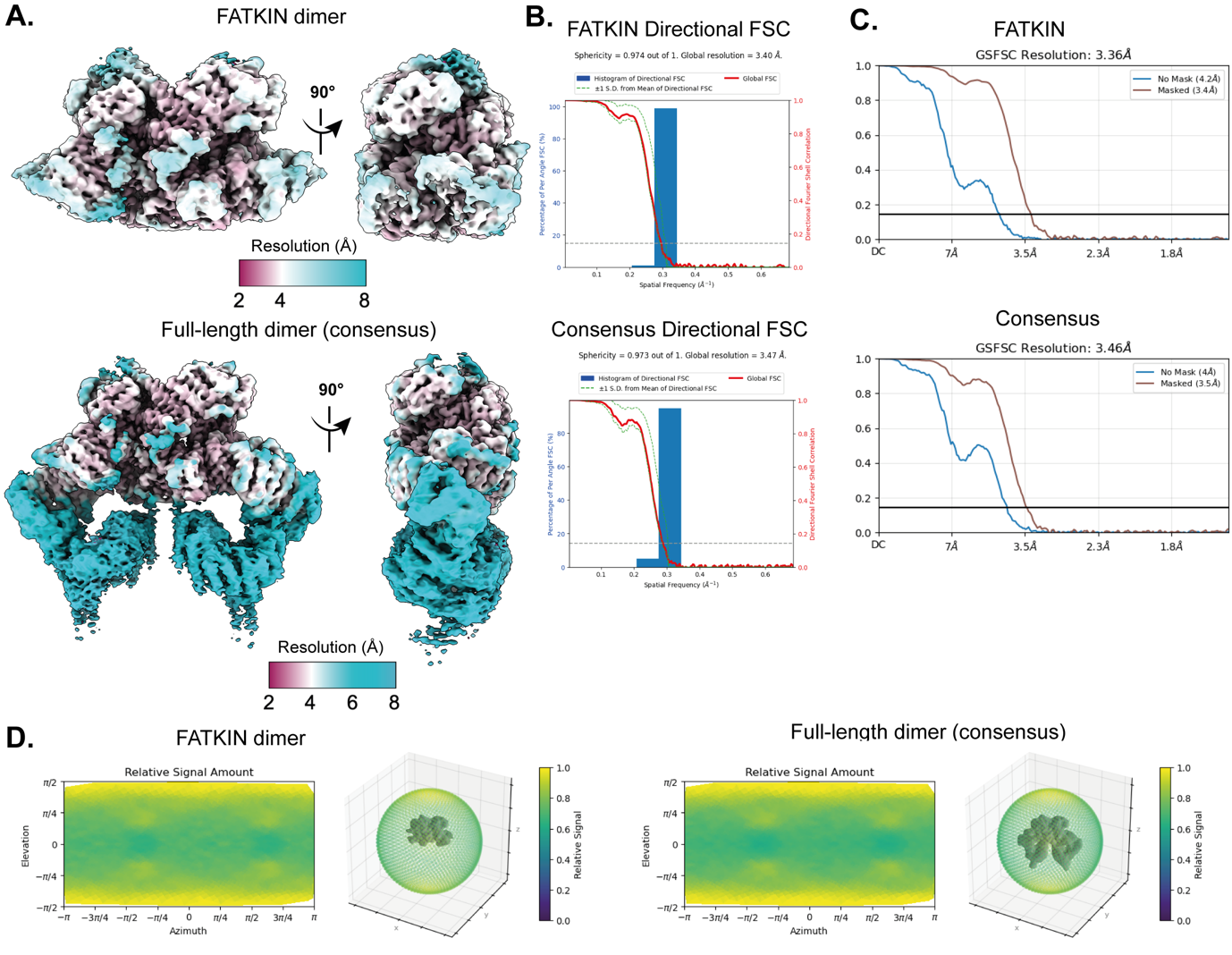

**Fig. S28.** **Cryo-EM map quality metrics for the WT/ATP.** (A) FATKIN (local refinement) and consensus maps of the WT bound to ATP colored by local resolution, estimated by *cryoSPARC* v4.7. (B) Directional FSC plots and histograms for the final reconstructions of FATKIN and consensus map of Δkα9b calculated using 3DFSC (*52*). (C) Gold-standard Fourier Shell Correlation (FSC) plot estimated by *cryoSPARC* v4.7. (D) 2D histograms and 3D view of the directional orientation (Euler angle) of the FATKIN and consensus maps of the WT bound to ATP. (E) Consensus and composite maps of WT bound to ATP.

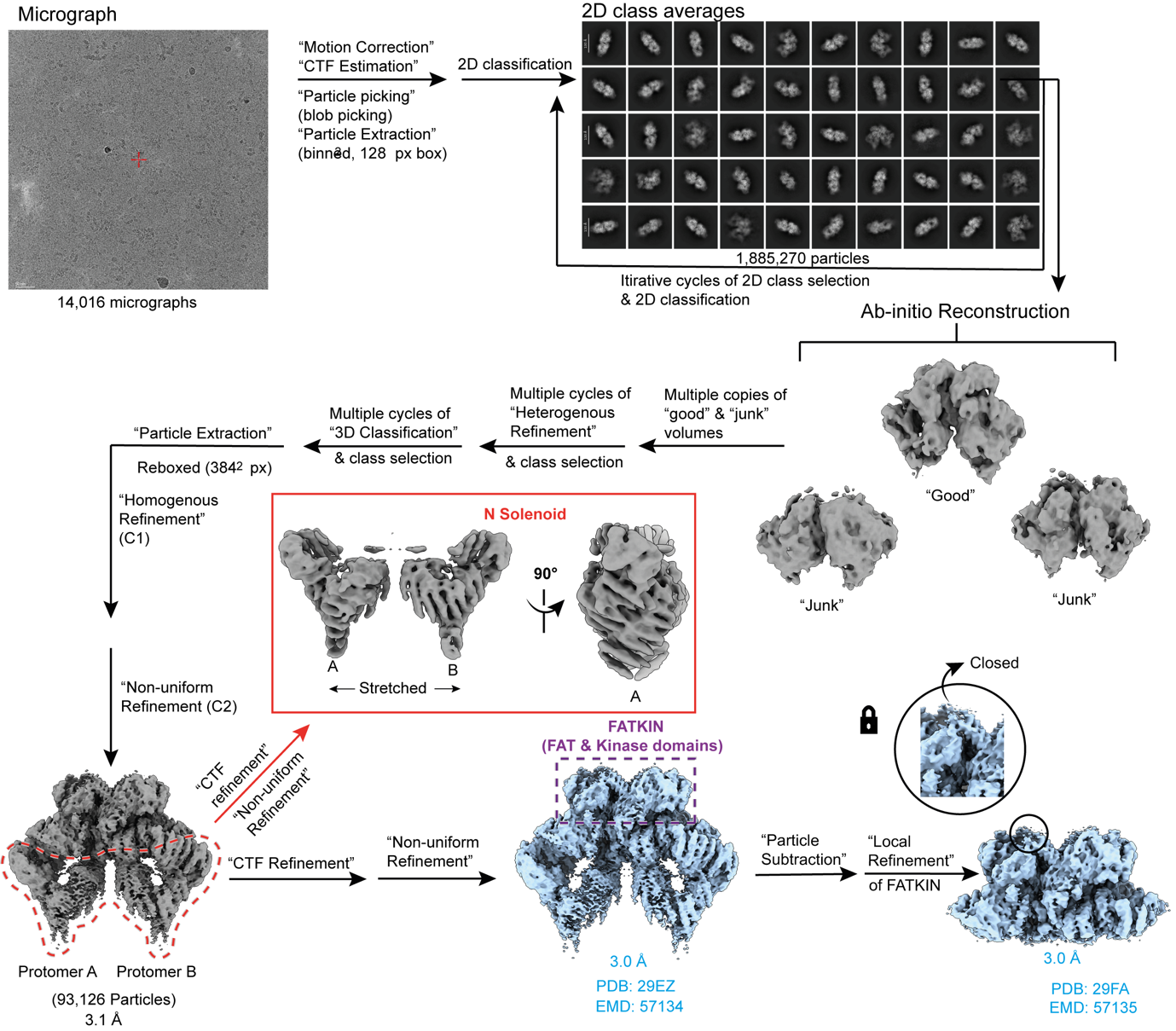

**Fig. S29.** **Cryo-EM single particle analysis data processing workflow of Δkα9b/ADP.** Final reconstructions deposited to the PDB and EMDB are indicated.

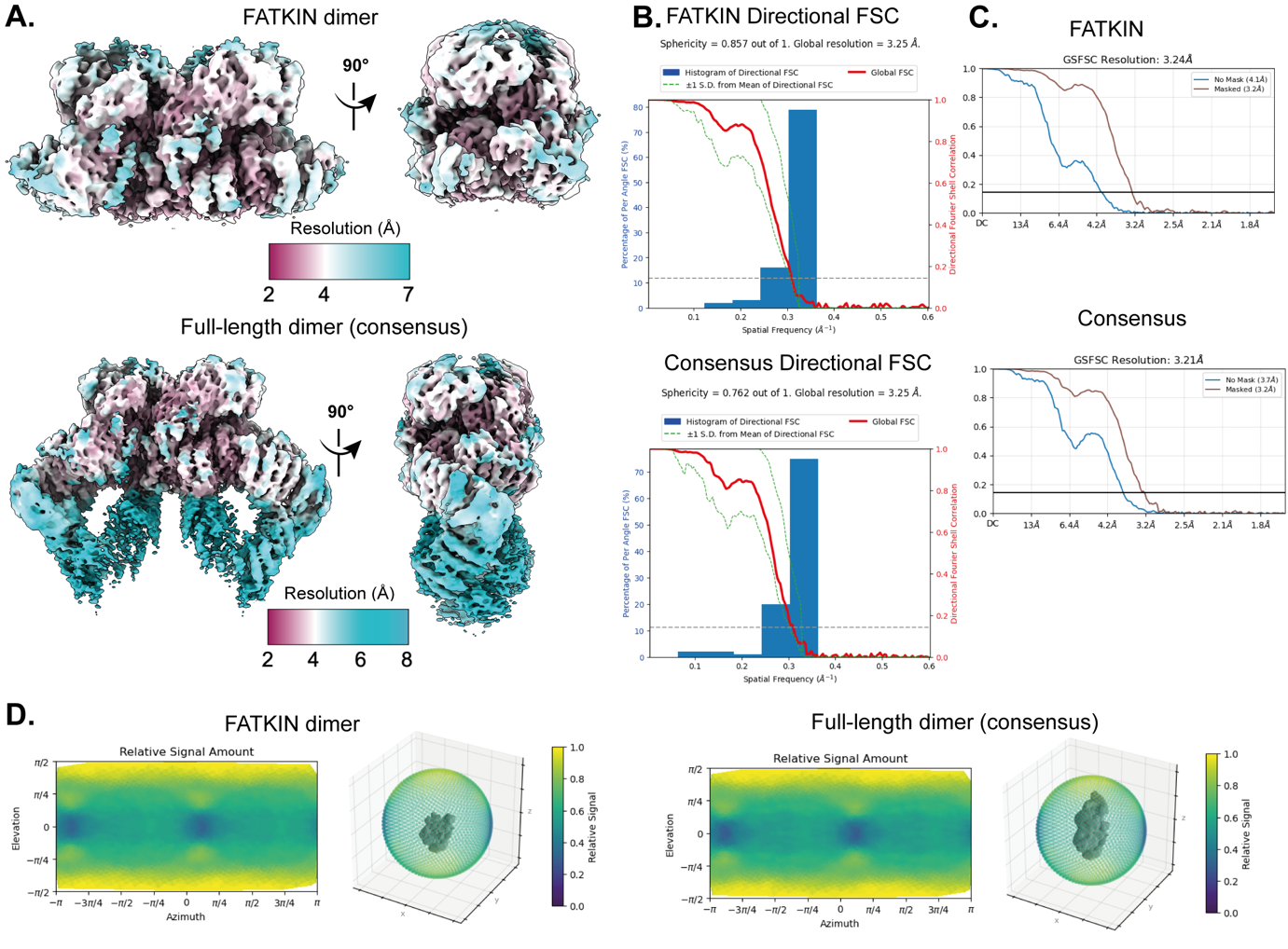

**Fig. S30.** **Cryo-EM map quality metrics for the Δkα9b/ADP.** (A)FATKIN (local refinement) and consensus maps of the Δkα9b bound to ADP colored by local resolution, estimated by *cryoSPARC* v4.7. (B) Directional FSC plots and histograms for the final reconstructions of FATKIN and consensus map of Δkα9b calculated using 3DFSC (*52*). (C) Gold-standard Fourier Shell Correlation (FSC) plot estimated by *cryoSPARC* v4.7. (D) 2D histograms and 3D view of the directional orientation (Euler angle) of the FATKIN and consensus maps of ATM Δkα9b bound to ADP. (E) Consensus and composite maps of ATM Δkα9b bound to ADP.

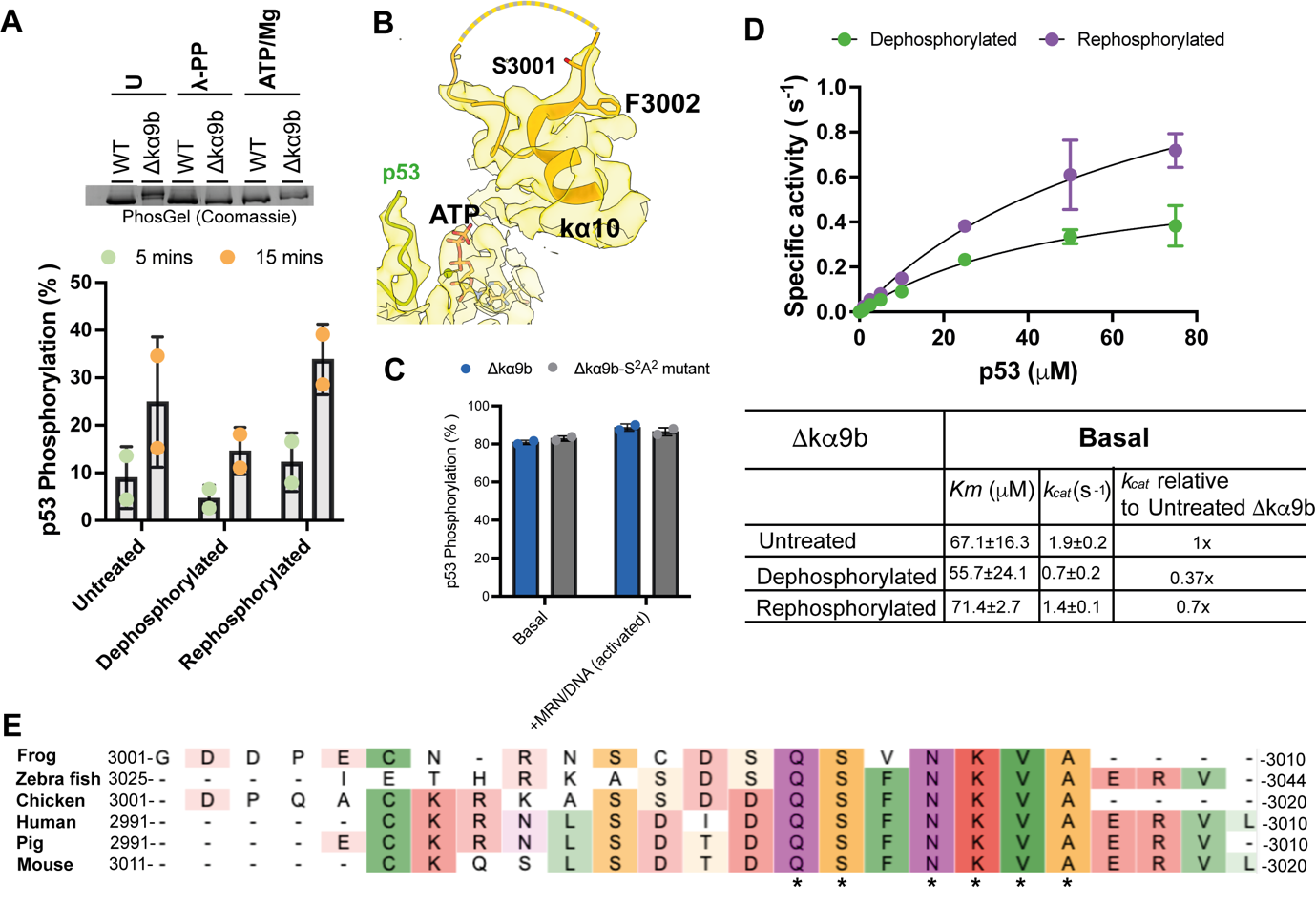

**Fig. S31. Dephosphorylation of hyperactive Δkα9b** **ATM only marginally affects its kinase activity.**

(A) PhosTag gel-based in vitro kinase assays of Δkα9b show only a small decrease in phosphorylation of p53^102^ when Δkα9b was dephosphorylated, and rephosphorylation of Δkα9b regained its kinase activity; inset is a PhosTag gel showing Δkα9b, but not WT, undergoes phosphorylation during protein expression. (B) Cryo-EM model and density map focusing S^3001^ of the PRD linker, which was identified as the autophosphorylated residue by phosphoproteomics (see fig. S32). (C) p53^102^ phosphorylation by Δkα9b and Δkα9b with Ser^2996^Ala and Ser^3001^Ala mutations (Δkα9b-S^2^A^2^). (D) *In vitro* enzyme kinetics showing a ~2.0-fold reduced activity for the globally dephosphorylated Δkα9b (see fig. S33). (E) Ser^2996^ and Ser^3001^ are conserved across lower lineages; hydrophobic residues are green, polar yellow, basic blue and acidic red.

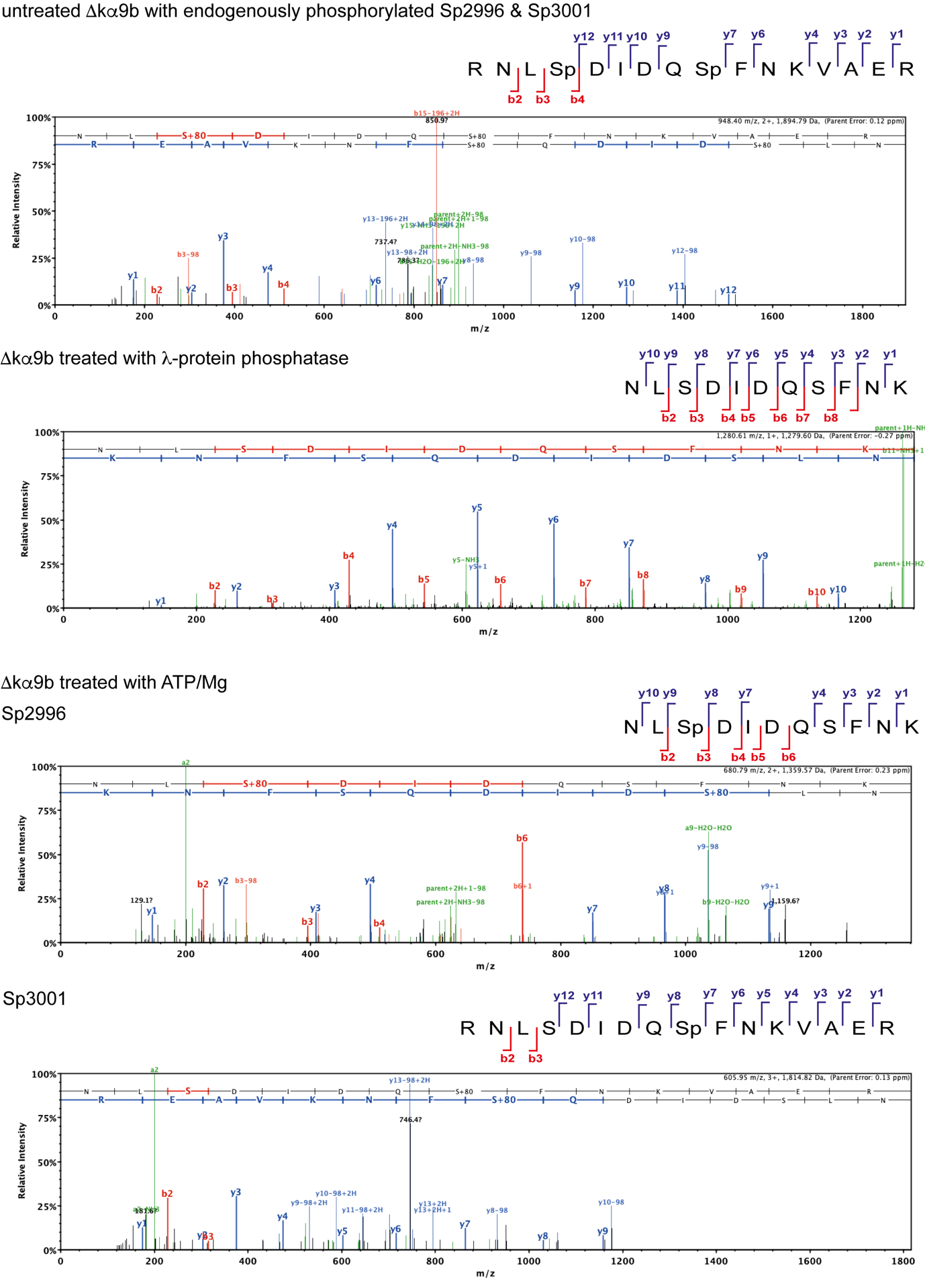

**Fig. S32. Identification of ATM phosphosites by LC-MS/MS**

Three samples of Δkα9b ATM mutant were analyzed for phosphosites by LC-MS/MS: (1) Δkα9b purified from mammalian Expi293 cells was used directly for phosphosite analysis (untreated ATM), or (2) it was first dephosphorylated by incubation with λ-PP for o/n (~16 hours) at 4°C (see fig. S33) or (3) dephosphorylated Δkα9b was autophosphorylated in vitro by incubation with ATP/Mg^+^ at 30°C for 0.5 hour. For each phosphosite, the corresponding peptide sequence is shown with the fragmentation pattern and the MS2 spectrum (b and y ions are shown in red and blue, respectively).

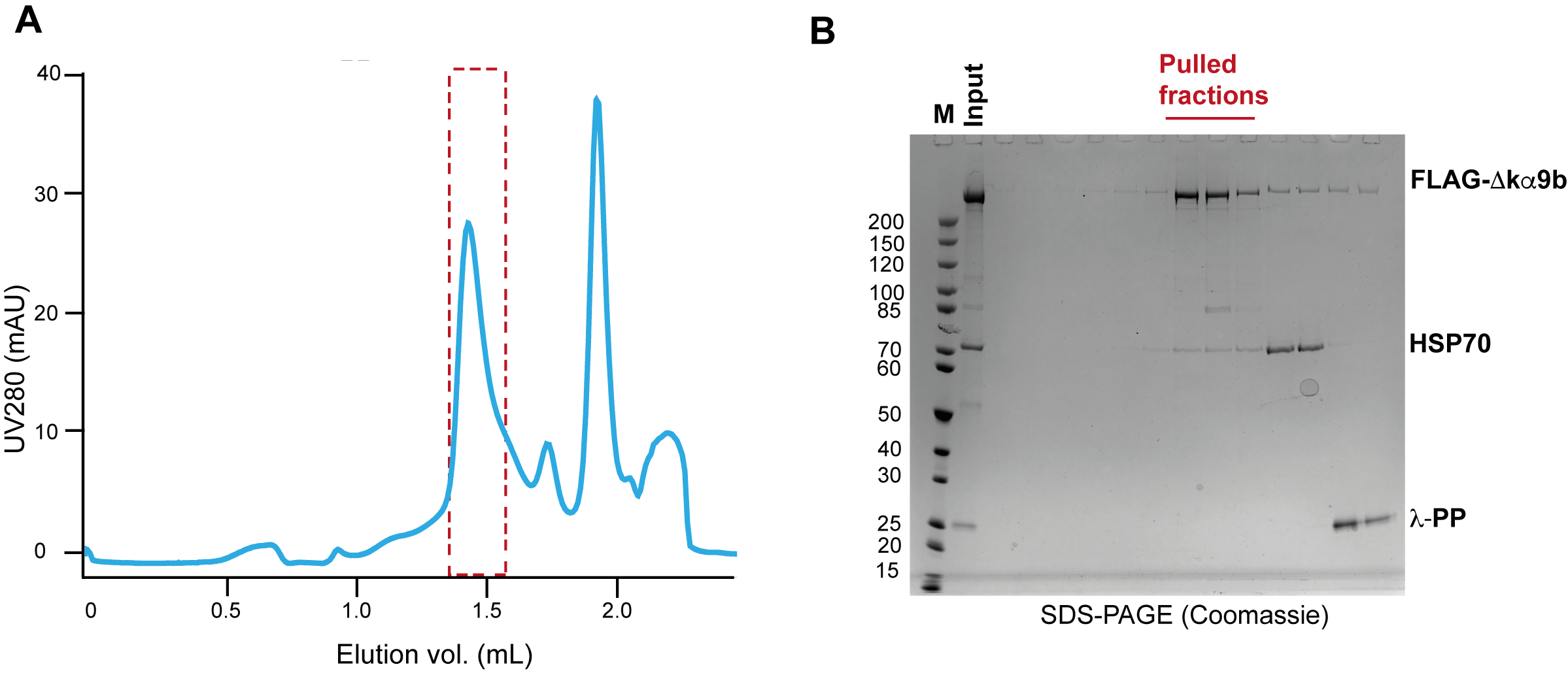

**Fig. S33. Dephosphorylation of Δkα9b.** (A) λ-PP-treated Δkα9b was purified using a Superose 6^TM^ Increase 3.2/300 column to remove λ-PP. (B) SDS-PAGE gel showing Superose 6 elution fractions with the pooled fractions highlighted, which later was used in the kinase assays and for LC-MS/MS analysis of Δkα9b (see figs. S31 and S32).

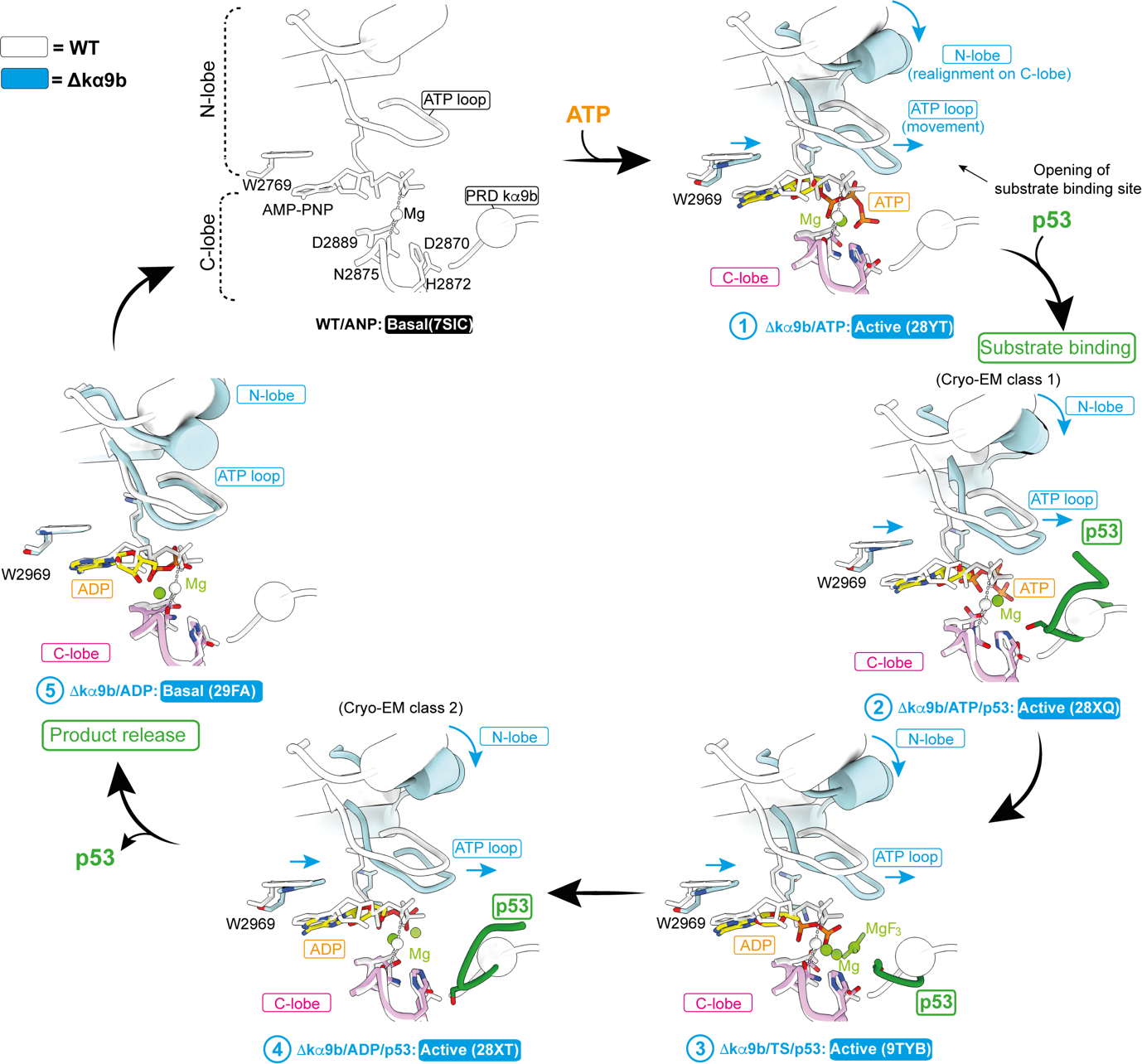

**Fig. S34. A schematic (proposed) showing substrate (p53) phosphorylation by ATM Δkα9b mutant.** Nucleotide/substrate bound cryo-EM structures of Δkα9b reported in this work (cyan), overlayed on ANP bound WT ATM (white, 7SIC), showing its (1) active-state conformation in the presence of ATP/Mg^+2^ only (28YT) and maintains its activated state (2) upon binding to substrate peptide p53 (28XQ), (3) in the presence of p53 and TS mimic (ADP/Mg_3_^-^) (9YTB), and (4) p53 and ADP (formed from ATP hydrolysis)/Mg^+2^ (28XT) but (5) returns to its basal/inactive state in the absence of p53 with bound ADP/Mg^+2^ only (29FA).

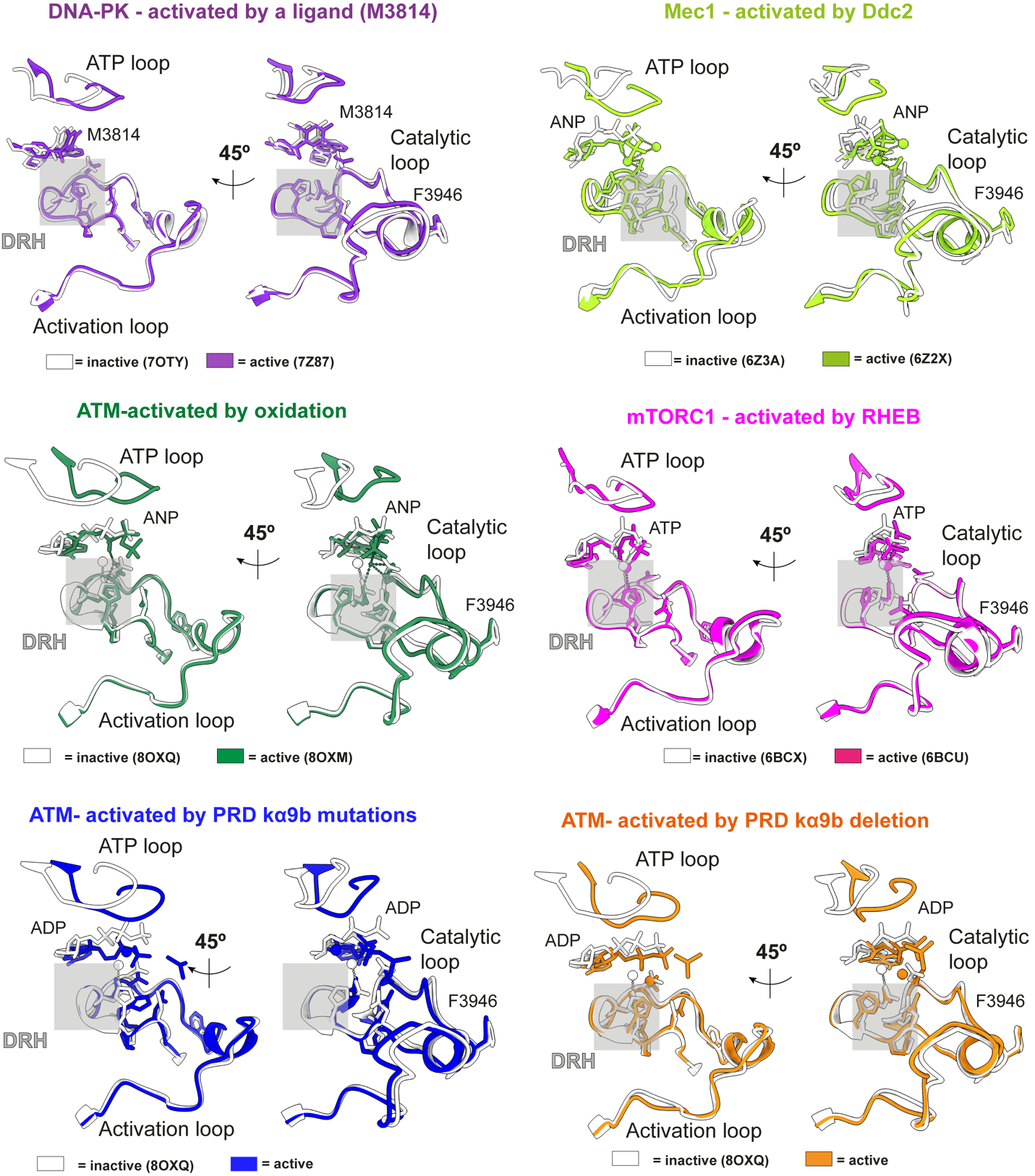

**Fig. S35. Activated cryo-EM structures of PIKKs.** ATP loop movement upon activation of DNA-PK by a ligand (M3814) (purple), Mec1 by Ddc2 (lime), ATM.Q2971A by oxidation (H_2_O_2_) (green), mTORC1 by RHEB (magenta), ATM by PRD mutation ^2968^LYLQ^2917^ to ^2968^LYLQ^2917^ (blue), and ATM by PRD residues 2963-2972 deletion including the helix kα9b (orange).

**Table S1.** **Cryo-EM data collection, refinement, and validation statistics for TS/p53 complexes of WT ATM, kα9b.4A and Δkα9b.**

|  | **WT/**  **ADP/MgF_3_^-^**  **(TS)** | | **kα9b.4A/**  **ADP/MgF_3_^-^/p53^12^ (TS)** | | **kα9b.4A/**  **ADP/MgF_3_^-^**  **(TS)** | | **Δkα9b/**  **ADP/MgF_3_^-^/p53^12^ (TS)** | |
| --- | --- | --- | --- | --- | --- | --- | --- | --- |
|  | **FL** | **FATKIN** | **FL** | **FATKIN** | **FL** | **FATKIN** | **FL** | **FATKIN** |
| PDB | 9U0Q | 9TPV | 9TY6 | 9TY7 | 9TY8 | 9TY9 | 9TYA | 9TYB |
| EMD | 56499 | 56116 | 56412 | 56413 | 56414 | 56415 | 56416 | 56417 |
| **Data collection and processing** |  | |  | |  | |  | |
| Microscope | Titan Krios II | | Titan Krios IV | | Titan Krios IV | | Titan Krios IV | |
| Detector | Gatan K3 (eBIC) | | Falcon 4i | | Falcon 4i | | Falcon 4i | |
| Magnification | 103,000x | | 130,000 | | 130,000 | | 130,000 | |
| Energy filter (eV) | 20 | | 10 | | 10 | | 10 | |
| Voltage (kV) | 300 | | 300 | | 300 | | 300 | |
| Electron exposure on sample (e^-^/Å^2^) | 35 e^-^/Å^2^ | | 43 e^-^/Å^2^ | | 43 e^-^/Å^2^ | | 39.4 e^-^/Å^2^ | |
| Target defocus range (mm) | -1 to -2.6 m | | -1 to -2.6 m | | -1 to -2.6 m | | -1 to -2.6 m | |
| Calibrated pixel size (Å) | 0.83 | | 0.955 | | 0.955 | | 0.955 | |
| Movies (no.) | 15,511 | | 9,502 | | 9,502 | | 9,838 | |
| Extraction box size (pixels) | 432 x 432 px | | 384x384 px | | 384x384 px | | 400x400 px | |
| Initial particle images (no.) | 1,515,053 | | 1,138,567 | | 1,138,567 | | 1,231,530 | |
| Final particle images (no.) | 114,154 | | 40,198 | | 39738 | | 43,014 | |
| Symmetry imposed | C2 | | C2 | | C2 | | C2 | |
| **Refinement** |  | |  | |  | |  | |
|  | FL | FATKIN | FL | FATKIN | FL | FATKIN | FL | FATKIN |
| Map resolution at FSC = 0.143 (Å) | 3.1 | 3.0 | 3.4 | 3.3 | 3.6 | 3.6 | 3.3 | 3.2 |
| Model composition |  |  |  |  |  |  |  |  |
| Non-hydrogen atoms | 89132 | 47020 | 89606 | 21375 | 87887 | 43079 | 88551 | 22028 |
| Protein residues | 5478 | 2898 | 5509 | 2570 | 5403 | 2662 | 5441 | 2726 |
| Nucleotides | 0 | 0 | 0 | 0 | 0 | 0 | 0 | 0 |
| Ligands |  |  |  |  |  |  |  |  |
|  | 2 Mg^+2^ | 2 Mg^+2^ | 4 Mg^+2^ | 4 Mg^+2^ | 2 Mg^+2^ | 2 Mg^+2^ | 4 Mg^+2^ | 4 Mg^+2^ |
|  | 1 MgF_3_^-1^ | 1 MgF_3_^-1^ | 1 MgF_3_^-1^ | 1 MgF_3_^-1^ | 1 MgF_3_^-1^ | 1 MgF_3_^-1^ | 1 MgF_3_^-1^ | 1 MgF_3_^-1^ |
|  | 1 ADP | 1 ADP | 1 ADP | 1 ADP | 1 ADP | 1 ADP | 1 ADP | 1 ADP |
|  | - | - | 2 Zn^+2^ | 2 Zn^+2^ | - | - | 2 Zn^+2^ | 2 Zn^+2^ |
| B factor (Å^2^) |  |  |  |  |  |  |  |  |
| Protein | 208.22 | 131.65 | 227.23 | 126.57 | 206.05 | 196.58 | 219.99 | 149.37 |
| Ligand | 110.33 | 118.65 | 107.24 | 107.4 | 63.7 | 146.53 | 113.47 | 109.19 |
| Bonds (RMSD) |  |  |  |  |  |  |  |  |
| Length (Å) | 0.003 | 0.004 | 0.005 | 0.004 | 0.005 | 0.005 | 0.003 | 0.003 |
| Angle (°) | 0.620 | 0.669 | 0.774 | 0.713 | 0.806 | 0.780 | 0.605 | 0.585 |
| Validation |  |  |  |  |  |  |  |  |
| Molprobity score | 1.01 | 0.99 | 0.88 | 1.25 | 1.02 | 0.94 | 1.12 | 1.24 |
| Clashscore | 1.55 | 1.45 | 1.08 | 2.08 | 1.29 | 1.07 | 2.14 | 2.7 |
| Poor rotamers (%) | 0.0 | 0.0 | 0.0 | 0.0 | 0.0 | 0.0 | 0.0 | 0.0 |
| Ramachandran plot |  |  |  |  |  |  |  |  |
| Favoured (%) | 97.42 | 97.45 | 97.71 | 96.09 | 97.07 | 97.33 | 97.29 | 96.95 |
| Allowed (%) | 2.49 | 2.52 | 2.19 | 3.82 | 2.77 | 2.63 | 2.68 | 3.01 |
| Disallowed (%) | 0.09 | 0.03 | 0.09 | 0.08 | 0.15 | 0.04 | 0.04 | 0.04 |

**Table S2. Quantification of the conformational changes noticed in the basal and activated-state structures of ATM**

|  | **N-lobe/C-lobe Rotation*** | | **TRD1/FAT-clamp*** | | **Protomer Rotation*** | |
| --- | --- | --- | --- | --- | --- | --- |
|  | Angle (º) | Displacement (Å) | Angle (º) | Displacement (Å) | Angle (º) | Displacement (Å) |
| WT/ADP/MgF_3_^-^ | 1.06 | 0.26 | 0.34 | 0.24 | 0.4 | 0.4 |
| WT/ATP | 0.45 | 1.01 | 0.7 | 1.11 | 0.61 | 1.31 |
| kα9b.4A/ADP/MgF_3_^-^/p53 | 9.8 | 2.7 | 12.6 | 5.9 | 14.0 | 7.8 |
| kα9b.4A/ADP/MgF_3_^-^ | 2.5 | 0.72 | 3.2 | 1.1 | 2.8 | 3.4 |
| kα9b.4A/ATP | 2.5 | 0.72 | 1.1 | 0.11 | 2.8 | 3.4 |
| Δkα9b/ADP/MgF_3_^-^/p53 | 10.8 | 2.7 | 12.9 | 6.1 | 13.6 | 7.5 |
| Δkα9b/ATP/p53 | 11.6 | 3.2 | 13.0 | 6.0 | 13.1 | 7.2 |
| Δkα9b/ADP**/p53 | 10.6 | 3.0 | 11.5 | 5.1 | 12.0 | 6.9 |
| Δkα9b/ADP/p53 | 9.8 | 3.0 | 12.5 | 5.5 | 12.5 | 6.7 |
| Δkα9b/ATP | 10.3 | 2.7 | 12.6 | 5.8 | 13.5 | 7.5 |
| Δkα9b/ADP | 1.5 | 0.53 | 1.1 | 0.8 | 1.5 | 2.41 |
| *Relative to the basal structure of wild-type ATM bound to AMP-PNP (PDB 7SIC)(*7*)  ** formed in situ from ATP hydrolysis | | | | | | |

**Table S3.** **Cryo-EM data collection, refinement, and validation statistics for Δkα9b/ATP/p53 (class 1), Δkα9b/ADP*/p53 (class 2), Δkα9b/ADP/p53.**

|  | **Δkα9b/**  **ATP/p53^12^** | | **Δkα9b/**  **ADP*/p53^12^**  **(ADP* formed in situ from ATP)** | | | | **Δkα9b**  **ADP/p53^12^** | |
| --- | --- | --- | --- | --- | --- | --- | --- | --- |
|  | p53 conformation 1  (**Class 1**) | | p53 conformation 2  (**Class 2**) | | | |  | |
|  | **FL** | **FATKIN** | **FL** | **FATKIN** | | | **FL** | **FATKIN** |
| PDB | 28XP | 28XQ | 28XS | 28XT | | | 29CQ | 29CR |
| EMD | 56944 | 56945 | 56947 | 56948 | | | 57081 | 57082 |
| **Data collection and processing** |  | |  | | | |  | |
| Microscope | Titan Krios IV | | Titan Krios IV | | | | Titan Krios IV | |
| Detector | Falcon 4i | | Falcon 4i | | | | Falcon 4i | |
| Magnification | 130,000 | | 130,000 | | | | 130,000 | |
| Energy filter (eV) | 10 | | 10 | | | | 10 | |
| Voltage (kV) | 300 | | 300 | | | | 300 | |
| Electron exposure on sample (e^-^/Å^2^) | 43.11 | | 43.11 | | | | 42.3 | |
| Target defocus range (mm) | -1 to -2.6 m | | -1 to -2.6 m | | | | -1 to -2.6 m | |
| Calibrated pixel size (Å) | 0.955 | | 0.955 | | | | 0.955 | |
| Movies (no.) | 8,900 | | 8,900 | | | | 7,546 | |
| Extraction box size (pixels) | 400x400 px | | 400x400 px | | | | 400x400 px | |
| Initial particle images (no.) | 1,248,403 | | 1,248,403 | | | | 1,020,730 | |
| Final particle images (no.) | 53,661 | | 40,402 | | | | 61,392 | |
| Symmetry imposed | C2 | | C2 | | | | C2 | |
| **Refinement** |  | |  | | | |  | |
|  | FL | FATKIN | FL | | FATKIN | FL | | FATKIN |
| Map resolution at FSC = 0.143 (Å) | 3.05 | 3.03 | 3.3 | | 3.26 ~3.3 | 3.2 | | 3.1 |
| Model composition |  |  |  | |  |  | |  |
| Non-hydrogen atoms | 88591 | 45116 | 67086 | | 22030 |  | | 34078 |
| Protein residues | 5459 | 2792 | 5481 | | 2728 |  | | 2112 |
| Nucleotides | 0 |  | 0 | | 0 |  | | 0 |
| Ligands |  |  |  | |  |  | |  |
|  | 2 Mg^+2^ | 2 Mg^+2^ | 4 Mg^+2^ | | 4 Mg^+2^ | 4 Mg^+2^ | | 4 Mg^+2^ |
|  | 2 ATP | 2 ATP | 2 ADP | | 2 ADP | 2 ADP | | 2 ADP |
| B factor (Å^2^) |  |  |  | |  |  | |  |
| Protein | 207.01 | 146.0 | 218.17 | | 139.33 |  | | 161.44 |
| Ligand | 90.27 | 126.75 | 90.69 | | 110.49 |  | | 126.07 |
| Bonds (RMSD) |  |  |  | |  |  | |  |
| Length (Å) | 0.005 | 0.004 | 0.003 | | 0.004 |  | | 0.004 |
| Angle (°) | 0.719 | 0.611 | 0.629 | | 0.685 |  | | 0.711 |
| Validation |  |  |  | |  |  | |  |
| Molprobity score | 1.13 | 1.00 | 1.48 | | 1.24 |  | | 1.5 |
| Clashscore | 1.04 | 1.21 | 4.13 | | 2.33 |  | | 3.19 |
| Poor rotamers (%) | 0.0 |  | 0.0 | | 0.0 |  | | 0.0 |
| Ramachandran plot |  |  |  | |  |  | |  |
| Favoured (%) | 95.36 | 97.06 | 95.94 | | 96.55 |  | | 94.3 |
| Allowed (%) | 4.51 | 2.9 | 3.91 | | 3.3 |  | | 5.46 |
| Disallowed (%) | 0.13 | 0.04 | 0.15 | | 0.15 |  | | 0.24 |

**Table S4.** **Cryo-EM data collection, refinement, and validation statistics for WT/ATP, kα9b.4A/ATP, Δkα9b/ATP, Δkα9b/ADP.**

|  | **WT/**  **ATP** | | **kα9b.4A/**  **ATP** | | **Δkα9b/**  **ATP** | | **Δkα9b/**  **ADP** | |
| --- | --- | --- | --- | --- | --- | --- | --- | --- |
|  | FL | FATKIN | FL | FATKIN | FL | FATKIN | FL | FATKIN |
| PDB | 29FT | 29FU | 29CG | 29CH | 28YT | 28YU | 29EZ | 29FA |
| EMD | 57137 | 57138 | 57077 | 57078 | 56984 | 56985 | 57134 | 57135 |
| **Data collection and processing** |  | |  | |  | |  | |
| Microscope | Titan Krios I | | Titan Krios II | | Titan Krios IV | | Titan Krios II | |
| Detector | Gatan K3 | | Gatan K3 | | Falcon 4i | | Gatan K3 | |
| Magnification | 105,000x | | 105,000x | | 130,000x | | 105,000x | |
| Energy filter (eV) | 20 | | 20 | | 10 | | 20 | |
| Voltage (kV) | 300 | | 300 | | 300 | | 300 | |
| Electron exposure on sample (e^-^/Å^2^) | 30.14 | | 39.4 | | 37.20 | | 40 | |
| Target defocus range (mm) | -1 to -2.6 m | | -1 to -2.6 m | | -1 to -2.6 m | | -1 to -2.6 m | |
| Calibrated pixel size (Å) | 0.73 | | 0.831 | | 0.95 | | 0.829 | |
| Movies (no.) | 5,539 | | 4,870 | | 9,844 | | 14,016 | |
| Extraction box size (pixels) | 480x480 px | | 480x480 px | | 448x448 px | | 384x384 px | |
| Initial particle images (no.) | 1,068,892 | | 1,147,043 | | 2,238,727 | | 1,885,270 | |
| Final particle images (no.) | 74,832 | | 91,867 | | 35,282 | | 93,126 | |
| Symmetry imposed | C2 | | C2 | | C2 | | C2 | |
| **Refinement** |  | |  | |  | |  | |
|  | FL | FATKIN | FL | FATKIN | FL | FATKIN | FL | FATKIN |
| Map resolution at FSC = 0.143 (Å) | 3.46~3.5 | 3.36~3.4 | 3.1 | 3.1 | 3.2 | 3.1 | 3.25 | 3.25 |
| Model composition |  |  |  |  |  |  |  |  |
| Non-hydrogen atoms | 90162 | 45806 | 88491 | 44880 | 88453 | 22032 | 88491 | 45646 |
| Protein residues | 5539 | 2825 | 5443 | 2772 | 5439 | 2724 | 5443 | 2820 |
| Nucleotides | 0 | 0 | 0 | 0 | 0 | 0 | 0 | 0 |
| Ligands |  |  |  |  |  |  |  |  |
|  | 2 Mg^+2^ | 2 Mg^+2^ | 2 Mg^+2^ | 2 Mg^+2^ | 2 Mg^+2^ | 2 Mg^+2^ | 2 Mg^+2^ | 2 Mg^+2^ |
|  | 2 ATP | 2 ATP | 2 ATP | 2 ATP | 2 ATP | 2 ATP | 2 ADP | 2 ADP |
| B factor (Å^2^) |  |  |  |  |  |  |  |  |
| Protein | 292.95 | 167.44 | 234.48 | 153.58 | 202.8 | 162.05 | 246.27 | 165.06 |
| Ligand | 161.84 | 171.18 | 90.28 | 146.54 | 97.57 | 130.35 | 80.6 | 140.48 |
| Bonds (RMSD) |  |  |  |  |  |  |  |  |
| Length (Å) | 0.003 | 0.004 | 0.006 | 0.002 | 0.005 | 0.002 | 0.005 | 0.005 |
| Angle (°) | 0.634 | 0.748 | 0.947 | 0.542 | 0.753 | 0.449 | 0.927 | 0.787 |
| Validation |  |  |  |  |  |  |  |  |
| Molprobity score | 0.99 | 0.55 | 0.81 | 0.96 | 0.84 | 1.44 | 1.02 | 0.73 |
| Clashscore | 1.7 | 0.13 | 0.1 | 0.58 | 0.63 | 4.92 | 1.35 | 0.42 |
| Poor rotamers (%) | 0.0 | 0.0 | 0.0 | 0.0 | 0.0 | 0.0 | 0.0 | 0.0 |
| Ramachandran plot |  |  |  |  |  |  |  |  |
| Favoured (%) | 97.71 | 98.75 | 96.01 | 96.13 | 97.30 | 96.95 | 97.15 | 97.59 |
| Allowed (%) | 2.01 | 1.18 | 3.76 | 3.76 | 2.66 | 3.01 | 2.81 | 2.37 |
| Disallowed (%) | 0.27 | 0.07 | 0.22 | 0.11 | 0.04 | 0.04 | 0.04 | 0.04 |

**Table S5. Plasmids used for protein expression**

| **Plasmid Number** | **Vector** | **Protein** |
| --- | --- | --- |
| AHp21 | pFB Dual | HsMRE11.H129N (untagged) and HsNBS1-TEV-2xStrep |
| AHp26 | pOPTHM(tev) | 6xHis-MBP(Tev)-Hsp53(1-102) |
| AHp63 | pFastBac | HsRAD50-TEV-2xStrep |
| AHp74 | pDEST12.2-OriP | 1xFlag-FKBP-9AA linker-HsATM-Δ2963-2972 (Δkα9b) |
| AHp75 | pDEST12.2-OriP | 1xFlag-HsATM |
| AHp79 | pDEST12.2-OriP | 1xFlag-FKBP-HsATM_2968-LYLQ-2971 to AAAA (kα9b.4A) |
| VPl15 | pLIX_403 (Addgene, #41395) | 1xFlag-HsATM-Δ2963-2972 (Δkα9b) |
| VPl16 | pLIX_403 | 1xFlag-HsATM |
| VPl17 | pLIX_403 | 1xFlag-HsATM 2968-LYLQ-2971 to AAAA (kα9b.4A) |

**Table S6. Primary Antibodies used**

| **Antibodies** | **Source** | **Identifier #** |
| --- | --- | --- |
| Rb ATM | Epitomics (Distributed by Insight Biotechnology) | ab32420 |
| Ms CHK2 | Millipore (U.K.) Ltd, | 05-649 |
| Rb pCHK2 T68 | Cell Signaling Technology (New England Biolabs) | 2661 |
| Ms FLAG M2 | Sigma- Aldrich | F3165 |
| Rb H2AX | Abcam Ltd | ab11175 |
| Ms γH2AX S139 | Millipore (U.K.) Ltd | 05-636 |
| Rb pKAP1 S824 | Bethyl Laboratories, Inc (Distributed by Universal  Biologicals (Camb) Ltd) | A300-767A-M |
| Rb MRE11 | Bio Techne | NB100-142 |
| Rb NBS1 | GeneTex, Inc. | GTX70224 |
| Rb pP53 S15 | Cell Signaling Technology (New England Biolabs) | 9284L |
| Rb PARP1 | Proteintech Europe Ltd | 13371-1-AP |
| Ms PARP1 | Bio Rad | MCA1522G |
| Ms RAD50 | Novus Biologicals | NB100-147 |

**Table S7. Secondary Antibodies used**

| **Antibodies** | **Source** | **Identifier** |
| --- | --- | --- |
| IRDye 800CW | LI-COR BIOSCIENCES UK Ltd | 926-32212 |
| IRDye 680LT | LI-COR BIOSCIENCES UK Ltd | 926-68021 |

**Movie S1. Human ATM is a homodimer.** Human ATM forms a homodimer where each protomer has Pincer (1-1166), Spincer (1167-1898), TRD1 (1899-2025), TRD2 (2026-2192), TRD3 (2193-2479), HRD (2480-2613), N-lobe (2614-2770), C-lobe (2771-2957), PRD (2958-3026), and FATC (3027-3057) domains. TRD = tetratricopeptide repeat domain, HRD = heat-repeat domain, PRD = PIKK regulatory domain.

**Movie S2. Movement of the FAT-clamp with respect to the kinase domain.** TRD2 (red) and TRD3 (green) as a unit (FAT-clamp) rotates with respect to the kinase domain (white) in the active-state conformation.

**Movie S3. Realignment of N-lobe on C-lobe.** N-lobe (powder blue) realigns on C-lobe (sky blue), which allows the ATP loop (part of N-lobe) and γ-phosphate of the bound nucleotide better register with the ligand-binding residues in the C-lobe.

**Movie S4. Movement of TRD3 fα21/fα22 helices.** Fα21/fα22 helices of TRD3 move away in the active-state conformation, making the substrate-binding pocket more accessible, which helps the incoming substrate (p53, green) bind to ATM. S2047/S2047’ residues are coloured magenta to follow as reference points. ADP, colored as yellow.

**Movie S5. Structural flexibility of ATM.** A morph between ATM’s basal (7SIC) and active state (8OXM) structures, showing the conformational changes in ATM homodimer’s structure in the transition going from basal to active.
